# Transcriptional profile of the rat cardiovascular system at single cell resolution

**DOI:** 10.1101/2023.11.14.567085

**Authors:** Alessandro Arduini, Stephen J. Fleming, Ling Xiao, Amelia W. Hall, Amer-Denis Akkad, Mark Chaffin, Kayla J. Bendinelli, Nathan R. Tucker, Irinna Papangeli, Helene Mantineo, Mehrtash Babadi, Christian M. Stegmann, Guillermo García-Cardeña, Mark E. Lindsay, Carla Klattenhoff, Patrick T. Ellinor

## Abstract

**Background:** Despite the critical role of the cardiovascular system, our understanding of its cellular and transcriptional diversity remains limited. We therefore sought to characterize the cellular composition, phenotypes, molecular pathways, and communication networks between cell types at the tissue and sub-tissue level across the cardiovascular system of the healthy Wistar rat, an important model in preclinical cardiovascular research. We obtained high quality tissue samples under controlled conditions that reveal a level of cellular detail so far inaccessible in human studies.

**Methods and Results:** We performed single nucleus RNA-sequencing in 78 samples in 10 distinct regions including the four chambers of the heart, ventricular septum, sinoatrial node, atrioventricular node, aorta, pulmonary artery, and pulmonary veins (PV), which produced an aggregate map of 505,835 nuclei. We identified 26 distinct cell types and additional subtypes, including a number of rare cell types such as PV cardiomyocytes and non-myelinating Schwann cells (NMSCs), and unique groups of vascular smooth muscle cells (VSMCs), endothelial cells (ECs) and fibroblasts (FBs), which gave rise to a detailed cell type distribution across tissues. We demonstrated differences in the cellular composition across different cardiac regions and tissue-specific differences in transcription for each cell type, highlighting the molecular diversity and complex tissue architecture of the cardiovascular system. Specifically, we observed great transcriptional heterogeneities among ECs and FBs. Importantly, several cell subtypes had a unique regional localization such as a subtype of VSMCs enriched in the large vasculature. We found the cellular makeup of PV tissue is closer to heart tissue than to the large arteries. We further explored the ligand-receptor repertoire across cell clusters and tissues, and observed tissue-enriched cellular communication networks, including heightened *Nppa* - *Npr1*/*2*/*3* signaling in the sinoatrial node.

**Conclusions:** Through a large single nucleus sequencing effort encompassing over 500,000 nuclei, we broadened our understanding of cellular transcription in the healthy cardiovascular system. The existence of tissue-restricted cellular phenotypes suggests regional regulation of cardiovascular physiology. The overall conservation in gene expression and molecular pathways across rat and human cell types, together with our detailed transcriptional characterization of each cell type, offers the potential to identify novel therapeutic targets and improve preclinical models of cardiovascular disease.

## Introduction

Although the mechanics of the cardiovascular system are conceptually simple, with the heart pumping blood through vessels, the physiology is complex. The system responds to multi-faceted hormonal and metabolic cues in seconds, delivering oxygen, nutrients, signaling molecules, and cells. Its intricate physiology parallels its anatomical diversity, with a four-chambered heart and permeable vessels. The anatomical diversity stems from elaborate organization at the cellular level, with cardiac fibers distinctly organized in each chamber, multi-layered blood vessels, and variable extracellular matrix composition.

A long-standing challenge in cellular physiology is detecting and enumerating the functionality of all cell types in a given tissue and comparing this with information from other tissues. Tremendous progress has been made in defining the physiology and pathology of the cardiovascular system over the last 150 years; however, profiling the transcriptional heterogeneity of cells has only become possible in the last decade.

Recently, we identified 15 cell types in the human heart using single-nucleus RNA sequencing (snRNA-seq) ^1^. However, working with human tissue presents many challenges. We therefore undertook an extended effort to study precisely-controlled, wide-ranging anatomical samples from a model organism, including the chambers of the heart, the ventricular septum, the vasculature, and the nodes of the conduction system. Model organisms, and small rodent species in particular, reduce variation from sampling bias, nutritional control, circadian cycles, genetics, and other environmental factors. Here we present an extensive repository of the cell types of the cardiovascular system in healthy Wistar rat, a strain largely used in academic and pharmaceutical cardiovascular research and pharmacology. We uncover a new level of cellular detail, including rare cell types and regional differences across the cardiovascular system.

## Results

### Detailed snRNA-seq atlas of the cardiovascular system in healthy Wistar rat

We collected 89 samples from 10 regions of the heart and major blood vessels: left ventricle (LV), right ventricle (RV), left atrium (LA), right atrium (RA), septum (base and apex), atrioventricular node (AVN), sinoatrial node (SAN), pulmonary vein (PV), pulmonary artery (PA), and aorta (Ao) (**Figure 1a**). We isolated nuclei and performed library preparation and sequencing. After quality control, 78 samples were retained (**Supplementary Table 1**). After additional cell quality control and the removal of doublets, 505,835 high quality nuclei comprised the final dataset (**Figure 1b**). The mean number of unique UMIs (unique molecular identifiers) per nucleus was 1151, with substantial variation across cell types (**Supplementary** Figure 1), which largely reflects previously observed differences in transcriptional complexity across cell types ^2^.

**Figure 1.**
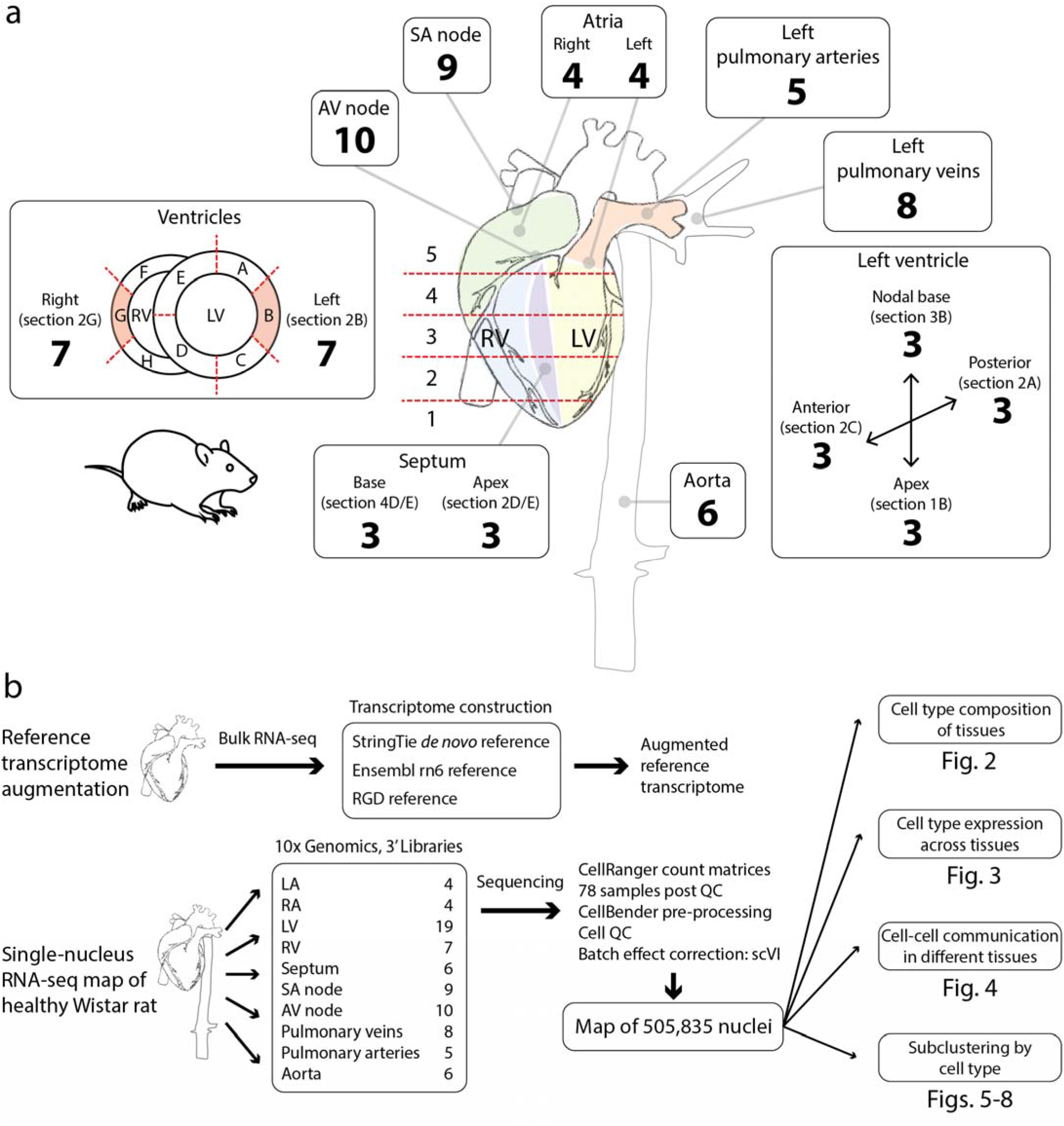
Overview of tissue sampling scheme and study design. **(a)** Extensive tissue sampling of regions of the healthy Wistar rat cardiovascular system. Numbers denote the samples that passed quality control. **(b)** Initial groundwork was laid by augmenting the reference transcriptome for the rat based on Wistar rat heart bulk RNA-seq. Samples were converted to snRNA-seq cDNA libraries using 10x Genomics 3’ mRNA capture. Using the augmented reference transcriptome, cellular transcriptional profiles were quantified and further processed and batch effects were removed, yielding a map of half a million nuclei. Downstream work included an examination of the cellular composition of tissues, cell-type expression profiles, putative cell-cell communication, and subclustering per cell type to achieve a detailed picture of cellular diversity across the cardiovascular system.

### Complete landscape of cell types in the rat cardiovascular system

We identified 27 cell clusters (**Figure 2a-b**), one of which (cluster 20) was omitted from downstream analyses because of the high fraction of mitochondrial transcripts. Among the remaining 26 clusters, we identified all cell types previously identified in the human heart ^1,3,4^, and identified additional rare cell types, such as non-myelinating Schwann cells (NM-SCs), and unique groups of vascular smooth muscle cells (VSMCs), endothelial cells (ECs), and fibroblasts (FBs). In the global UMAP, we observed satellite micro-clusters belonging to other main clusters, suggesting additional cellular complexity, some of which is explored in the following sections.

**Figure 2.**
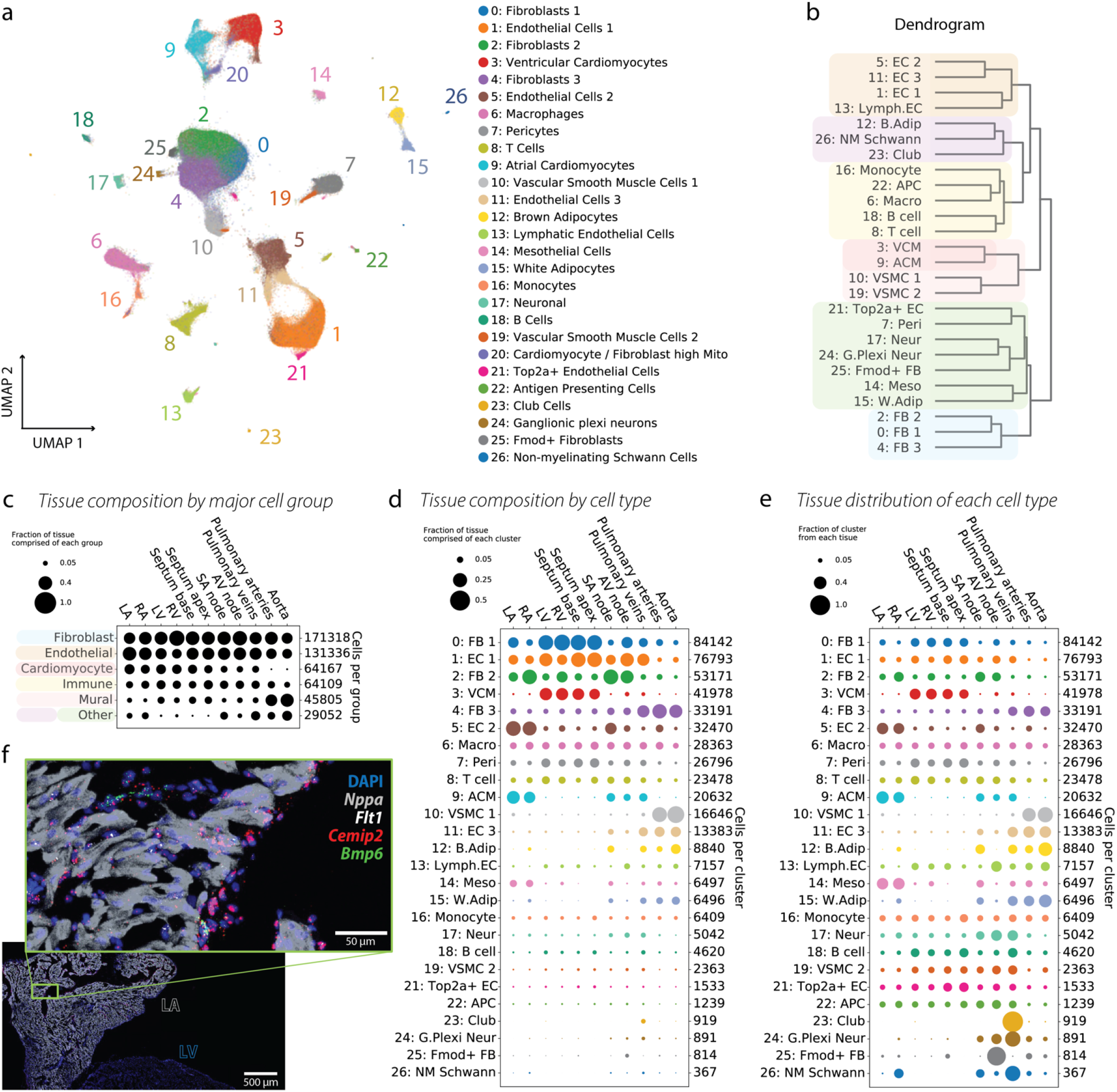
High-level overview of the cardiovascular cell atlas. **(a)** Map of all 505,835 nuclei measured in this study. Clustering was performed using the Leiden algorithm with resolution 1.2, and reveals 27 distinct cell types, as well as several additional micro-clusters that do not separate out at this clustering resolution. **(b)** Dendrogram displaying the hierarchy of cluster relationships. **(c)** Composition of each tissue region in terms of cell type abundance. Cell types are grouped into a few coarse-grained categories. Dot sizes are normalized so that each column sums to 1. **(d)** Composition of each tissue region in terms of the cell types found in **a**. Dot sizes are normalized so that each column sums to 1. **(e)** Location of each cell type. Dot sizes are further normalized so that each row sums to 1. **(f)** RNAscope imaging was performed to validate the presence of cluster 5: EC2 in the LA and its relative absence in the LV, which is expected based on the data in **d-e**. *Cemip2* (red) and *Bmp6* (green) are enriched markers for EC2, while *Flt1* (white) is a general EC marker. Tissue was counterstained with the nuclear marker DAPI (blue) and the atrial CM marker *Nppa* (gray). Inset shows magnification of LA tissue, with EC2 visible in the endocardium.

We calculated marker genes for each cluster (**Supplementary** Figure 2), identifying multiple previously known marker genes (e.g. *Myh6* and *Ttn* in cardiomyocytes (CMs)), and several novel marker genes (e.g. *Opcml* in FBs and *Nuak1* in ECs). Identification of these and other novel markers (**Supplementary Table 2**) may reflect a nuclear enrichment of specific transcripts, or species specificity, or it may reflect true cell type specialization across cardiac tissue and the large vasculature. We also performed gene set enrichment analysis (GSEA) to identify relevant cellular functions across cell clusters (**Supplementary** Figure 2) and between clusters which are more closely related on the cluster dendrogram (**Supplementary** Figure 3).

### Cellular composition of different tissue regions

This scale of the current dataset allowed us to perform an extensive analysis of the tissue architecture of the cardiovascular system. We began by generating super-clusters by combining similar cell types (e.g. all ECs, all FBs, all CMs, see dendrogram in **Figure 2b**) and assessed their distribution across tissues (**Figure 2c**). We find that ECs and FBs are the major constituents of all tissues (with the exception of the PA and Ao) and that there are differences in cellular composition across regions, which are linked to anatomical or functional differences. The ratio of ECs to FBs is higher in the atria compared to ventricles, mostly explained by a greater luminal surface to wall mass ratio in the atria and large enrichment of endocardial ECs.

However in the large vasculature (PA and Ao), the proportion of mural cells (VMSCs and pericytes) is larger. VMSCs are as abundant as ECs and FBs in the PA, and VSMCs are the most abundant cell type in the Ao.

We illustrate the proportion of each tissue composed of each cluster (**Figure 2d**), as well as the proportion of each cluster which comes from each tissue (**Figure 2e**). We observe compartmentalization of certain cell types, including: (1) the presence of two CM populations, which as anticipated map to the atria and ventricles, (2) a large proportion of VSMC1 mapped to the arterial tissues and a second more ubiquitous population, VSMC2, (3) an enrichment of the FB2 cluster in the RA and the embedded/neighboring tissues AVN and SAN, (4) a marked enrichment of FB3 and EC3 in the large vasculature, and (5) a higher pericyte abundance in ventricles and septum, and lower abundance in the large arteries.

Given the enrichment of EC2 in the atrial tissues and the SAN, we identified marker genes that differentiate the EC clusters and used them to perform RNAscope imaging validation. *Ptprb* and *Flt1* were used as ubiquitous EC marker genes, while *Cemip2* and *Bmp6* are mostly enriched in the EC2 cluster (**Supplementary** Figure 4c). By imaging coronal sections of the rat heart, we found that the ubiquitous EC marker *Flt1* is present similarly in atria and ventricles, while *Cemip2* and *Bmp6* are mainly localized to atrial tissue in the endocardial regions (**Figure 2f**). Using information from GSEA, marker genes, and imaging experiments, we have defined the EC clusters as follows: capillary EC (EC1), endocardial EC (EC2), large vessel EC (EC3), lymphatic EC (LymphEC), and cycling capillary EC (*Top2a*^+^ EC). We refer to **Supplementary Section S.1.2** for further discussion of findings related to the cellular composition of each tissue.

### Transcriptional differences across tissues

We then examined tissue-specific differences in transcription for each cell type. Here we highlight findings for FBs, ECs, and VSMCs; however, this same approach can be applied to all cell types, and later sections will discuss the CMs in detail.

A differential expression analysis revealed marker genes that differentiate FBs (aggregated analysis of clusters 0, 2, 4) from all other cell types, separately in each geographical region of the cardiovascular system (**Figure 3a**). We tabulated the number of tissues in which each gene is a FB marker, and identified a small number of “robust” markers which delineate FBs in every tissue, and this list includes the well-characterized markers *Gsn* and *Dcn*. A larger number of genes delineate FBs in specific tissue contexts (**Figure 3b**). We separately performed a differential expression test among only FBs to identify genes whose expression varies significantly across tissues (**Supplementary** Figure 5a), and used this to prioritize a list of tissue-specific FB marker genes, whose expression which we plot alongside the ubiquitous FB marker genes in **Figure 3c**. The “common” marker genes have a varying level of expression across tissues; however, they are FB markers in all tissues. In contrast, the tissue-specific FB markers might be present in one or a few tissues and nearly absent in others, such as *Csmd1* (atrial tissues) and *Col1a2* (arteries) (**Figure 3c**).

**Figure 3.**
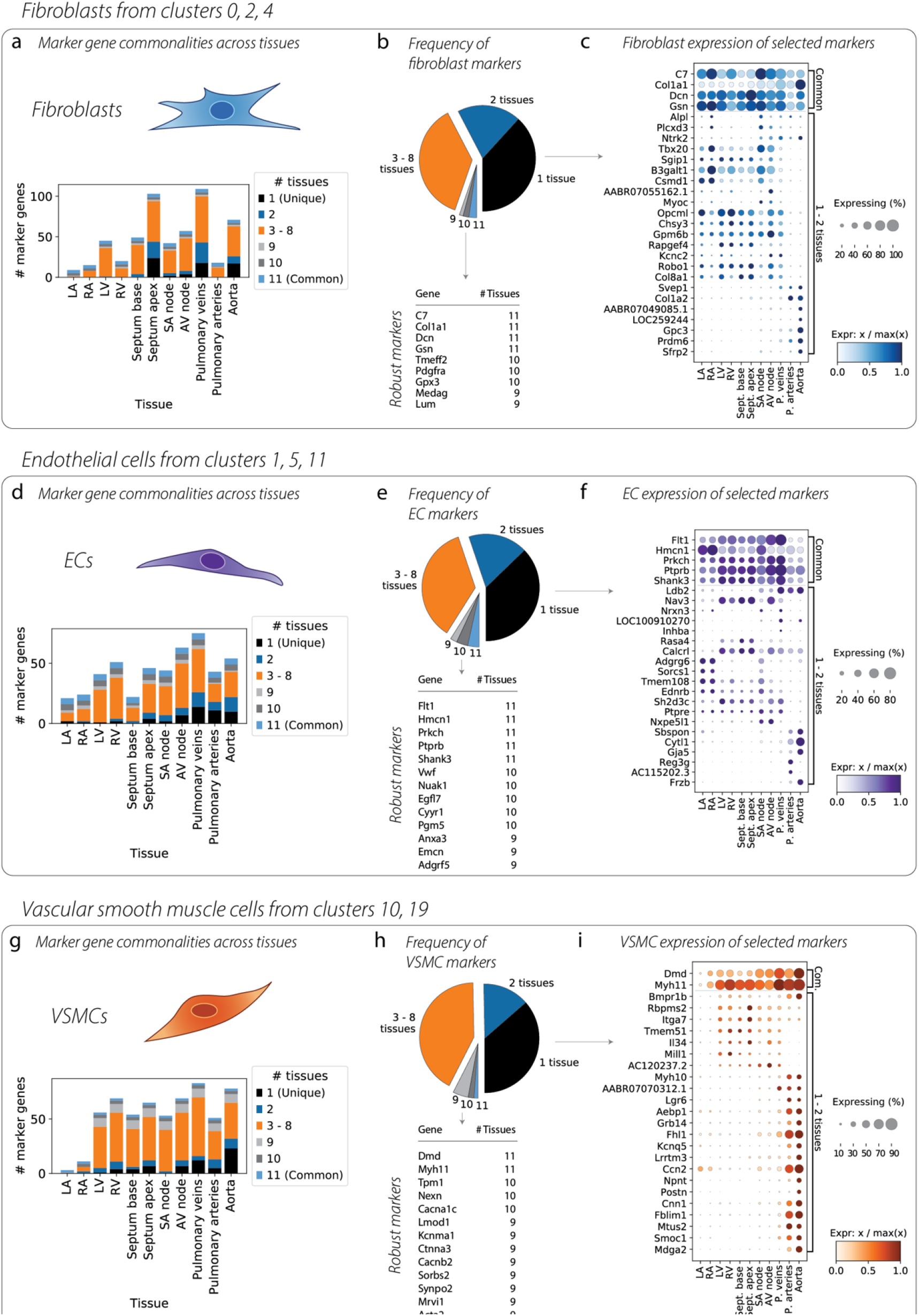
Differences in marker genes across tissues. Differential expression tests for one cell type versus all other cells were conducted separately in each tissue. **(a)** Breakdown of FB marker genes in each tissue, showing the number of unique as well as shared markers. **(b)** Pie chart shows the proportion of markers that were tissue-unique versus common, and lists the marker genes common to nearly all tissues. **(c)** Dotplot contrasts the common marker genes (top) with genes found to be markers in only one or two tissues. Genes are prioritized based on a second differential expression test of one tissue versus all others, within FB cells, and the genes with the most variability across tissues are shown. **(d, e, f)** Same analysis for ECs. **(g, h, i)** Same analysis for VSMCs.

We performed the same analysis for ECs (**Figure 3d-f**), finding the tissue-specific marker *Adgrg6* in atrial tissues, *Nav3* in ventricular tissues, and *Cytl1* in arterial tissues. And again for VSMCs (**Figure 3g-i**), we find that *Mill1* and *Il34* are VSMC markers in ventricular tissues, while *Fblim1* is a marker in the vasculature, and *Postn* is a marker in the aorta. These findings highlight the high degree of molecular diversity in defining cell specialization across tissues.

### Cell-cell communication networks and their tissue specificity

Next, we used the CellPhoneDB database ^5^ to characterize cell-cell communication based on annotated ligand-receptor interactions (**Figure 4a**). Putative interaction strengths for specific cell-type pairs can be computed based on expression of ligands and receptors in our dataset (**Supplementary** Figure 6a-d), and we compute these interaction strengths separately for each biological sample in order to limit interactions to local ones. ECs were responsible for a large share of cell-cell communication. A few of the top paracrine ligand-receptor pairs responsible for the major links depicted in **Figure 4a** are displayed in **Supplementary** Figure 6e.

**Figure 4.**
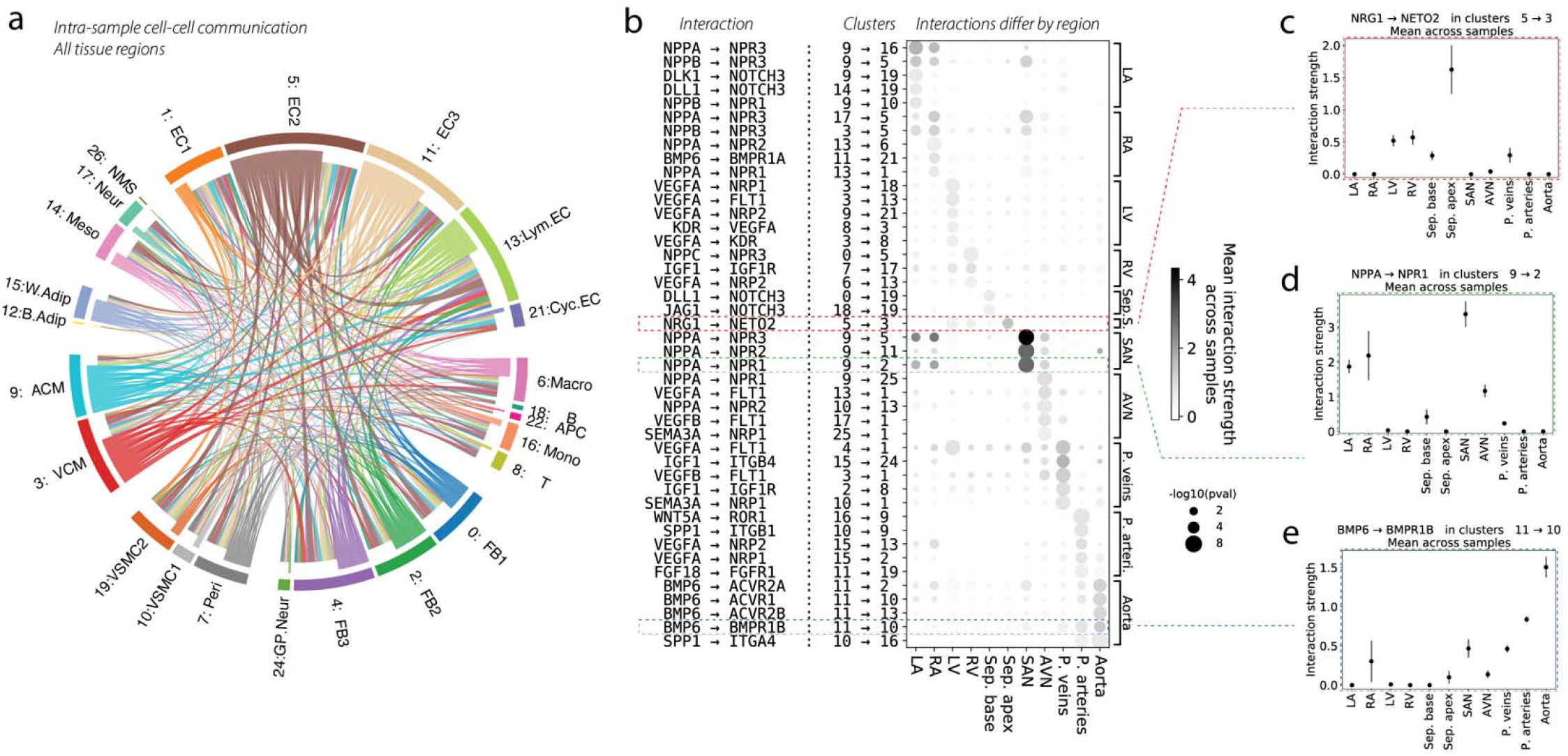
Cell-cell communication and its variability across the cardiovascular system. **(a)** Putative cell-cell interactions, computed separately for each sample and then aggregated over samples, are shown as a chord diagram, where chords are colored by the cell type that secretes the ligand, and the width of each chord is proportional to the number of significant interactions that surpass some minimum interaction strength. **(b)** Dotplot that highlights cell-cell interactions which vary greatly by tissue. Rows represent particular interactions, each of which is annotated by the ligand-receptor interaction as well as the cell type clusters involved. Mean interaction strength for the ligand-receptor interaction is denoted by dot color. Dot size is a p-value for enrichment computed from a Wilcoxon test for differences in interaction strengths across tissues. The top five (or fewer) significant interactions for each tissue are shown, comprising a small part of a much longer list. The red, green, and blue dashed boxes highlight interactions which are examined in panels (c-e). **(c)** The interaction strength of Nrg1 from cluster 5: EC2 → Neto2 from cluster 3: VCM is shown for each tissue. Error bars represent the standard deviation across samples. This interaction is clearly quite enriched in the septum apex. **(d)** Similar plot for the interaction of Nppa from cluster 9: ACM → Npr1 from cluster 2: FB2, which shows a pattern of atrial enrichment. **(e)** Similar plot for the interaction of Bmp6 from cluster 11: EC3 → Bmpr1b from cluster 10: VSMC1, which is enriched in the vasculature and the aorta in particular.

We highlight the relevance of *Vegfa* signaling from VCM and ACM to several other cell types, such as EC clusters and immune clusters, which receive the signal mainly through *Flt1*, *Npr1* and *Npr3* (**Supplementary** Figure 6e). We identify communication between ACM and many other cell types, including EC1, Lymphatic EC, monocytes, neural cells, and cycling ECs via the *Nppa*-*Npr1* pathway.

A few of the top autocrine interactions are displayed in **Supplementary** Figure 6f, where it is shown that EC2 and EC3 each communicate with themselves via *Bmp6*. These figure panels represent a small portion of a much larger dataset, which is included as **Supplementary Table 6**.

Many ligand-receptor interactions were upregulated in specific tissues, and we have prioritized those interactions which differ most across tissues in **Figure 4b**. We observe many tissue-enriched cellular communication networks. For example, there is a particularly high interaction strength in the SAN between *Nppa* in ACM and *Npr1*/*2*/*3* receptors in EC and FB which is highlighted in **Figure 4d**. The enrichment of these interactions in the SAN may have direct implications for the regulation of heart rate; however, the enrichment of *Npr1* in FB2 suggests additional biological functions.

Interestingly, we also identified communication routes specific to the arterial tissues, with a dominant role of the *Bmp6* ligand toward multiple receptors (*Acvr2a*, *Acvr1*, *Acvr2b*, *Bmpr1b*) in the Ao, and with *Wnt5a*-*Ror1* being strongest in the PA. These examples reflect differences in tissue physiology and aid in the interpretation of divergent tissue responses among the large blood vessels. **Figure 4c-e** indicates that there is large variability in interaction strengths across samples from different tissues. The full analysis of all significant ligand-receptor pairs across cell type pairs in each individual tissue is presented in **Supplementary Table 7**.

### Subclustering CMs reveals pacemaker cells and other phenotypic variability

Starting with 53,862 high-quality nuclei from CM clusters (3 and 9, **Figure 5a**), we ran the clustering algorithm at a higher resolution, resulting in 8 distinct subclusters (**Figure 5b**). The tissue distribution of these subclusters is striking, particularly for CMs derived from nodal tissue and PVs (**Figure 5c**, **Supplementary** Figure 7a). Although nearly all CMs from ventricular samples map to the “ventricular” subclusters, there are a very small but nonzero number of “atrial-like” cells in ventricular tissues. We decided to look for these cells by staining for *Nppa* in the ventricles, and indeed were able to identify a very small number of CMs in ventricular tissues expressing a high level of *Nppa* (**Figure 5d**).

**Figure 5.**
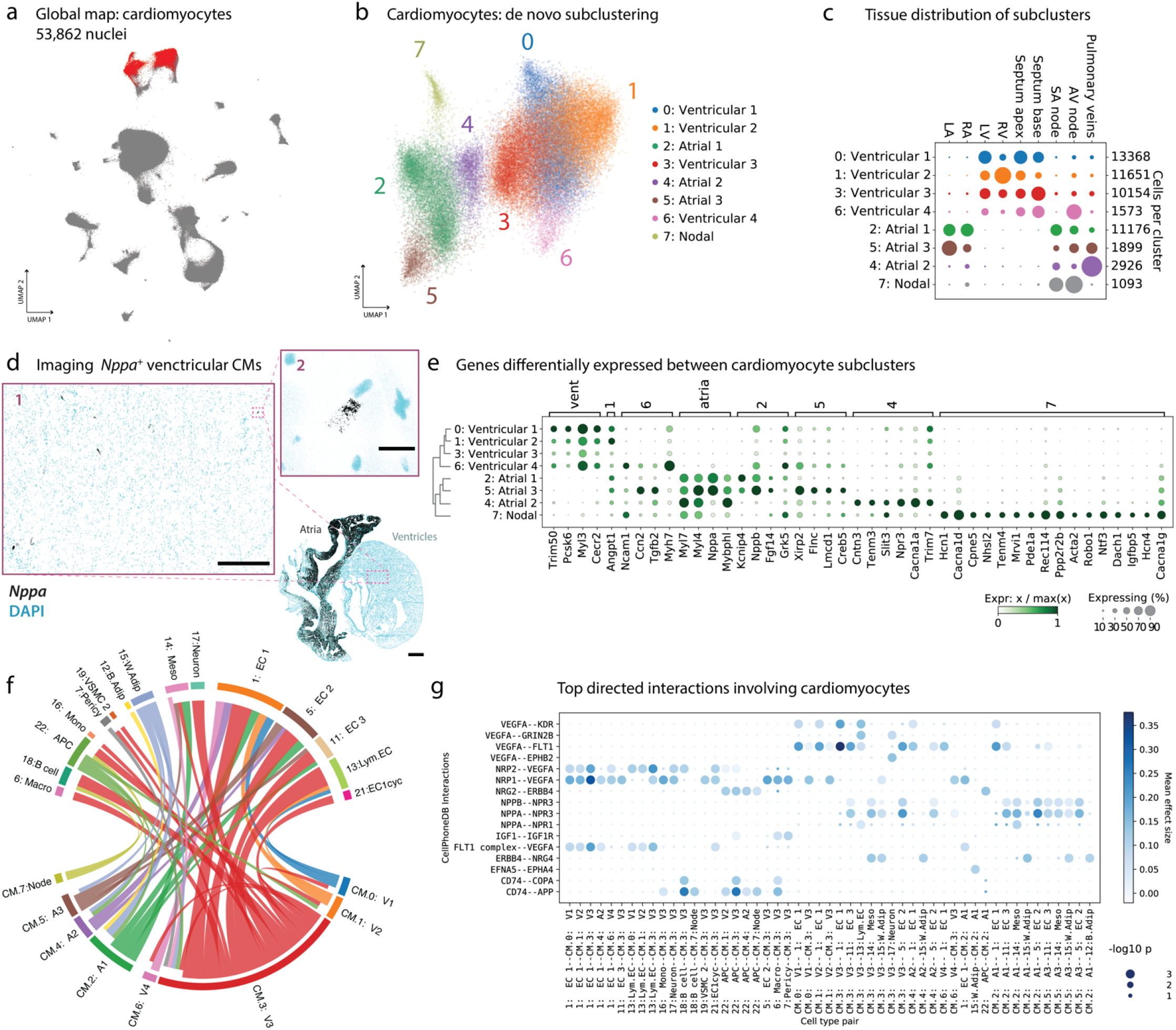
High-resolution subclusters of cardiomyocytes (CMs). (a) All high-quality CMs from the global UMAP are shown in red. **(b)** *De novo* subclustering of the CMs reveals a large amount of transcriptional variability, here shown as a UMAP. **(c)** Distribution of the CM subclusters across tissues. Dot sizes sum to one in each row. **(d)** RNAscope validation showing that there are an exceedingly small number of *Nppa*-positive CMs in the ventricles. Scale bar on full heart image is 1mm. Inset 1 scale bar is 300 microns. Inset 2 scale bar is 20 microns, and shows one such cell. **(e)** Dotplot showing top differentially expressed genes across CM subclusters. The nodal pacemaker cells from subcluster 7 have many marker genes. **(f)** Chord diagram showing significant cell-cell interactions involving CMs, broken down by CM subcluster. Chord width is proportional to the number of significant ligand-receptor interactions, and chords are colored by the cell type which secretes the ligand. **(g)** Dotplot showing the top ligand-receptor interactions involving the CM subclusters. Dot color denotes the interaction strength, while dot size denotes p-value.

Figure 5e shows the results of a differential expression test which identified genes that are enriched in specific CM subclusters. The dendrogram on the left-hand vertical axis of the dotplot shows that there is a clade of ventricular subclusters, a clade of atrial subclusters, and a “nodal” subcluster, distinct from the other two but more closely related to the atrial clade. The ventricular clade is enriched for genes including *Myh7*, *Myl2*, and *Myl3*, while the atrial clade is enriched for *Myl7*, *Nppa*, *Myl4*, and *Mybph1*, among many others. The nodal subcluster contains pacemaker CMs, with cells from both the SAN and AVN, and these cells are enriched for automaticity markers *Hcn1* and *Hcn4*, and neuronal identity markers *Tenm4*, *Robo1*, *Cacna1d*, *Sv2c*, *Ntf3*, and *Vsnl1*.

We also find a subcluster of CMs (“4: Atrial 2” in Figure 5b) which is quite specific to the PV. However, we can be reasonably confident this is not driven by a batch effect due to the presence of other CM subclusters in PV tissue as well (see Figure 5c). This subcluster is not easily characterized by a single, highly-specific marker gene, though it has higher levels of *Cntn3* and lower levels of *Nppa* than other atrial CMs (Figure 5e). We examine differential expression in this cluster versus other atrial CMs in **Supplementary** Figure 7b, where we see that *Cacna1a* is also upregulated in the PV CM subcluster 4 (expression plotted on a UMAP in **Supplementary** Figure 7c). We carried out RNAscope imaging validation in a tissue section which contained both LA and PV. *Cacna1a* shows higher expression in the PV (**Supplementary** Figure 7d), in agreement with the snRNA-seq results. It is known that the PV contains different types of cardiomyocytes with unique electrophysiological properties, which may contribute to the initiation and maintenance of atrial fibrillation ^6^. Whether this novel PV CM subcluster underlies the unique PV electrophysiological phenotype warrants further functional validation.

We re-examined cell-cell communication with CMs subclustered at this fine-grained resolution, and we found that communication differs by subtype (Figure 5f-g). Plots highlighting some of the top interactions involving the CM subclusters are shown in **Supplementary** Figure 7g-k. Nodal pacemaker CMs have the least number of significant interactions, some of which involve *App* receptor response to *Cd74* in immune cells (**Supplementary** Figure 7h).

### Detailed cellular map of the SA and AV nodes

A UMAP of all the cells from AVN and SAN tissue samples is shown in Figure 6a. The distribution of all cell types across these nodal tissues is shown in Figure 6b, and the results of a test for genes differentially expressed between the cluster of pacemaker CMs and the cluster of atrial CMs is shown in Figure 6c. A notable new cluster in this map at this resolution (not annotated in the global map of Figure 2a) is a cluster of nodal pacemaker CMs (cluster 16, containing 1086 cells), which contains the same pacemaker cells identified above via CM subclustering.

**Figure 6.**
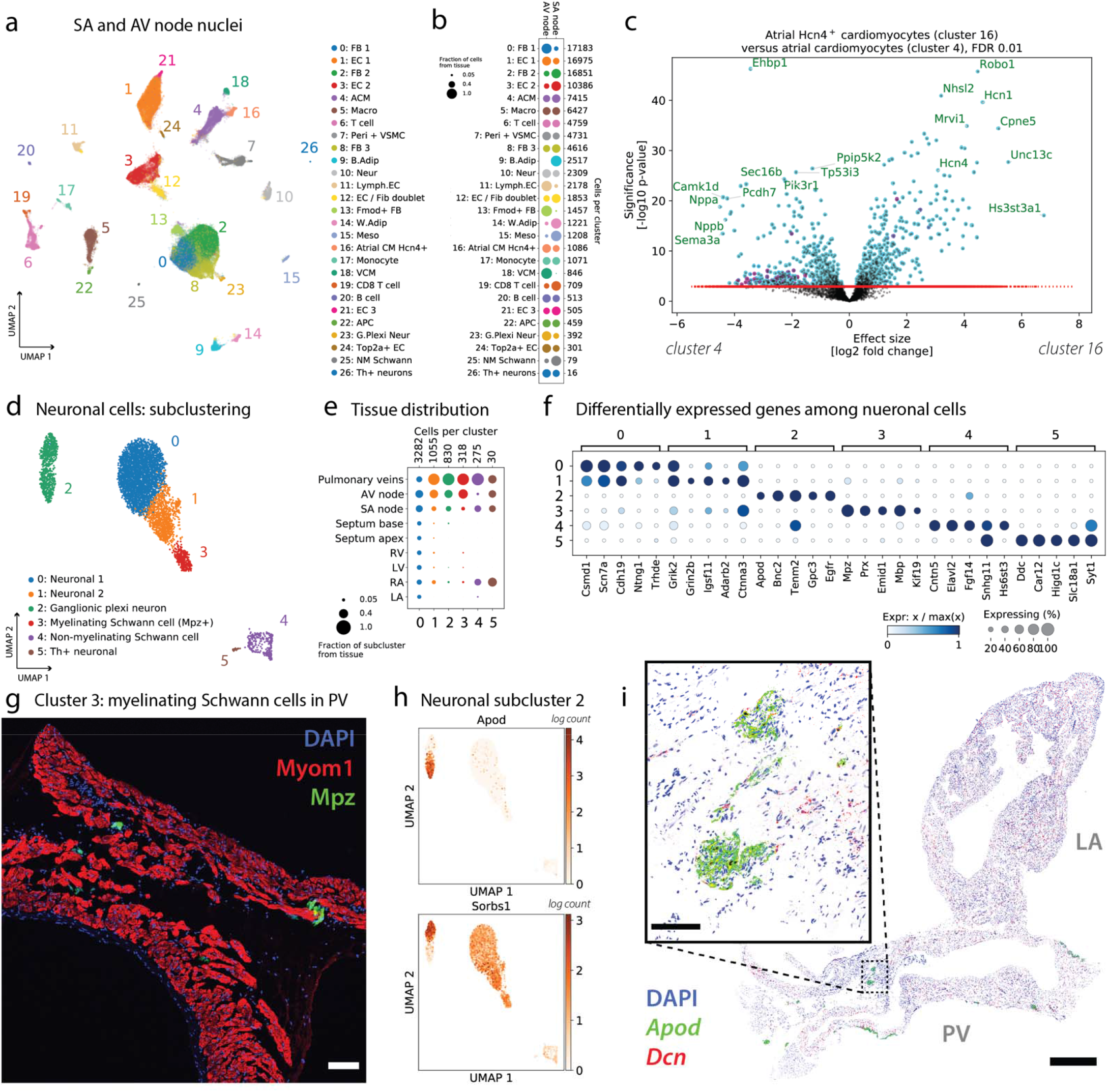
Cardiac conduction: nodal and neuronal cell populations. **(a)** UMAP of all 108,063 nuclei from the SAN and AVN, clustered and annotated. Cluster 16 captures the nodal pacemaker CMs. **(b)** Representation of clusters in the SAN and AVN. **(c)** Volcano plot shows the results of a differential expression test between the nodal pacemaker CMs and atrial CMs. Both cell types are present in the same samples, eliminating confounding batch effects. **(d)** Subclustering of all 5790 neuronal cells from the atlas yields 6 subtypes. **(e)** Distribution of neuronal subtypes across cardiovascular tissues. **(f)** Top marker genes for each neuronal subtype. **(g)** Immunofluorescence imaging of *Mpz*^+^ (green) subcluster 3, the myelinating Schwann cells in PV tissue. Tissue was counterstained with the nuclear marker DAPI (blue) and the CM marker *Myom1* (red). Myelinating Schwann cells make contact with CMs. Scale bar is 100 microns. **(h)** The genes *Apod* and *Sorbs1* are expressed in different subsets of neuronal subcluster 2. **(i)** RNAscope image (white background) of tissue section containing LA and PV, staining *Apod* (green) and *Dcn* (red; FB marker). Tissue was counterstained with the nuclear marker DAPI (blue). Inset shows groups of *Apod*^+^ cells. Full field of view scale bar is 1mm, and the inset scale bar is 100 microns.

Markers strongly upregulated in pacemaker CMs include *Hcn1* and *Hcn4* (see **Supplementary** Figure 8a), *Robo1*, *Cpne5*, *Unc13c*, and *Hs3st3a1*, among others. Many genes are also strongly downregulated in pacemaker CMs as compared to atrial CMs, including *Nppa*, *Nppb*, and *Camk1d*. GSEA results for pathways differentially regulated in atrial CMs and pacemaker CMs are shown in **Supplementary** Figure 8b, and a direct comparison of the expression of genes in pacemaker CMs between the SAN and AVN is shown in **Supplementary** Figure 8c. Among other genes, we see *Shox2* strongly upregulated in the SAN as compared to the AVN, consistent with Liu *et al.*^7^

A similarly detailed map of the PV cells is shown in **Supplementary** Figure 9, along with discussion in **Supplementary Section S.5**. Additionally, separate maps of the AVN (**Supplementary** Figure 10) and SAN (**Supplementary** Figure 11) are discussed in **Supplementary Section S.4**.

### Six neuronal cell types show diverse patterns of tissue specificity

We next sought to explore other facets of cardiac conduction, including the external inputs to the heart and neuronal cells within the heart. Here we characterize 5790 high-quality neuronal cells in our dataset (**Supplementary** Figure 12a) in more detail by clustering them at a higher resolution. Six subclusters are identified in the UMAP shown in Figure 6d, along with a pseudo-bulk PCA plot in **Supplementary** Figure 12b.

The six neuronal cell types include putative ganglionic plexi neurons, two other distinct populations of neurons, myelinating and non-myelinating Schwann cells, and a few *Th*^+^ neurons, which could represent sympathetic peripheral nervous system inputs to the heart. The neurons in subcluster 0 have a fairly uniform distribution across tissues, while the other subclusters are tissue-specific (Figure 6e). The more rare subclusters 2-5 are predominantly found in nodal tissue and the PV.

Marker genes which distinguish the neuronal subtypes are displayed in Figure 6f. While subclusters 0 and 1 are the most similar, they can still be distinguished based on the upregulation of *Trhde* in subcluster 0 and the upregulation of *Grin2b* in subcluster 1 (among other genes). The other subclusters are more transcriptionally divergent, each having highly specific markers: *Apod* and *Bnc2* for the ganglionic plexi neurons, *Mpz* for the myelinating Schwann cells, *Cntn5* for the non-myelinating Schwann cells, and *Ddc* for the *Th*-positive neurons (see also **Supplementary** Figure 12d-e). Canonical marker genes of the sympathetic and parasympathetic nervous system are plotted in **Supplementary** Figure 12f, and enriched pathways from GSEA are shown in **Supplementary** Figure 12g.

We performed imaging validation for two of these neuronal subtypes. The myelinating Schwann cells (subcluster 3) are marked by *Mpz*, and so we used immunofluorescence to image Mpz and Myom1 (a CM marker) in PV (Figure 6g) and LA tissue (image not shown). Myelinating Schwann cells are identified within PV myocardial sleeve but not in LA and appear to make contact with CMs in the PV.

The ganglionic plexi neurons in subcluster 2 comprise two groups of cells which are clearly differentiated by the expression of *Apod*, as shown in Figure 6h (and **Supplementary** Figure 12c). We used RNAscope to image *Apod* (along with *Vit* and *Cacna1a*) in a tissue section containing both LA and PV (Figure 6i). *Apod* was identified in cluster-like structures within non-CM groups of nuclei. These clusters were found in adventitial regions of the PV and not in the LA (in agreement with the expected tissue distribution from Figure 6e). The presence of these nerve fibers and ganglionic plexi at the PV-LA junction in humans, and the implications for heart rhythm disorders, is reviewed by Tan *et al.* ^8^.

### Endothelial cells have discrete subtypes, some unique to the vasculature

In this and the following section, we take up one particular tissue niche – the vasculature (further detail in **Supplementary Section S.6.6**; **Supplementary** Figure 13) – in order to explore cellular transcriptional variability in a specific context.The gene expression of ECs in our study can be summed together to produce a pseudo-bulk measurement per sample. We used principal component analysis (PCA) to quantify the major axes of transcriptional variability across these pseudo-bulked ECs in Figure 7a. The top principal component of variation (x-axis) clearly differentiates between tissues, with ventricular tissues on one side, large arteries on the other, and atrial tissues and PV somewhere in the middle. Thus, overall EC transcription is quite different in these various tissue contexts.

**Figure 7.**
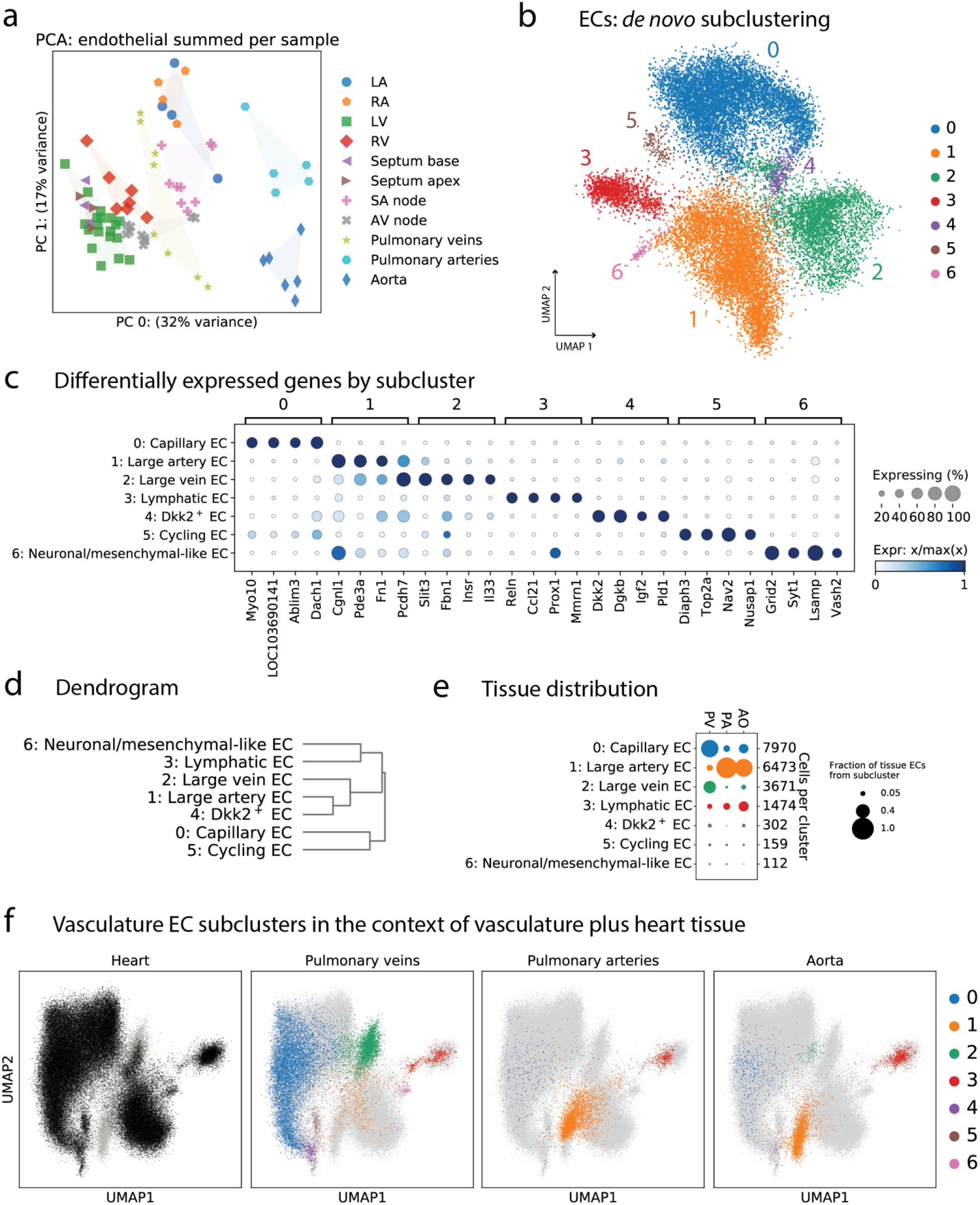
Distinct transcriptional subtypes of ECs in the vasculature. (a) Pseudobulk variation in EC transcription. Each dot is the summed expression of all ECs from one sample. Coloring by tissue shows that the PA and Ao samples clearly separate from other tissues in the top principal component. **(b)** Subclustering of the ECs from PV, PA, and Ao samples reveals 7 subtypes. **(c)** Marker genes of each subcluster show clear distinctions. **(d)** Dendrogram shows that subclusters 1 and 2, the large artery and large vein ECs, are part of the same clade. **(e)** The EC composition of PV, PA, and Ao shows that the arteries are quite different from PV. **(f)** UMAPs where each point is one cell, and all ECs from all tissues are shown. In each panel, all cells are shown in light gray, overlaid by cells present in the tissue specified. Colors correspond to the vascular subclusters from panel b.

To explore what drives these differences, we subclustered ECs in the vasculature at a high resolution (Figure 7b). The ECs display remarkable transcriptional variability, and can be grouped into 7 subclusters with specific marker genes and tissue distribution (Figure 7c-e). Notably, ECs were by far the most heterogeneous cell type across cardiovascular tissues. This fascinating heterogeneity is consistent with previous reports describing EC diversity across tissues in other species.^35,36^, and it suggests complexity beyond tissue origin or blood vessel type or diameter.

We can visualize the EC composition of the PV, PA, and Ao graphically using a UMAP, where the map contains all ECs in the entire dataset, including other tissues (see **Supplementary** Figure 14). This UMAP of all ECs can be used to contextualize the ECs found in the vasculature (Figure 7f; each tissue shown separately in **Supplementary** Figure 14h). In Figure 7e the PA and Ao are principally composed of subcluster 1, while Figure 7f puts subcluster 1 into a broader context and suggests that large blood vessels are composed of ECs with unique phenotypes. These phenotypes could arise from regions of higher shear stress or other physiological adaptations. We were able to validate the existence of these subtypes of ECs in vascular tissues using RNA-scope (**Supplementary** Figure 15, with additional exploration of phenotypically-relevant marker genes in **Supplementary** Figure 16). Indeed, Figure 7f demonstrates that the cellular makeup of PV tissue is closer to heart tissue than PA or Ao, and this explains much of the variability in Figure 7a. For further detail, please see **Supplementary** Figure 14e, which juxtaposes variability due to subcluster and tissue of origin.

Large artery ECs (subcluster 1, Figure 7) (La-EC) comprise the majority of ECs in the PA and Ao, and this subtype is largely absent from other tissues. This subcluster is characterized by high expression of *Eya4* and *Meis1*. In zebrafish, *Meis1* morphants have a reduced expression of dorsal artery markers and an increase in dorsal vein markers.^37^ Thus, we can speculate that different physiology in large arteries and a different developmental origin of aortic EC act as dual drivers in the specialization of *Meis1^+^/Eya4^+^* La-EC.^38^ The small vascular EC subcluster named “neuronal/mesenchymal-like” (subcluster 6, Figure 7, 112 nuclei) could reflect the identification of valvular ECs, due to their tissue distribution and the presence of *Hapln1*, *Selp*, and *Vwf*; or could be a novel subtype of ECs with neuronal-like properties, due to the presence of the glutamate receptor *Grid2*, the Ca^2+^ binding protein *Sty1*, and the neuronal surface glycoprotein *Lsamp*.

Blood vessels are chronically exposed to mechanical forces of various origin and magnitude, which are known drivers of the phenotypic specialization of ECs.^26,27^ We find EC subclusters with an increased expression of *Klf2*, *Klf4*, and *Ass1* both in the arterial and venous vasculature (**Supplementary** Figure 16a), highlighting different mechanical load across vascular beds. Interestingly, the expression of Klf2 targets is different in the venous and arterial EC clusters. Specifically, *Nos3* expression is much higher in La-ECs compared to large vein ECs (subcluster 2, Figure 7) (Lv-EC), which is consistent with a higher NOS3 staining and nitric oxide production found in internal thoracic artery compared to saphenous vein.^40^ It is known that Klf2 activation represses the activity of Nf-kb and the expression of endothelial activation genes, such as *Vcam1*.^28^ However, in the La-EC and Lv-EC we observe co-expression of the Klf2 and Nf-kb responses. Since both transcription factors compete for the p300 transcriptional coactivator, it is possible that p300 presence is higher at the Nf-kb complex in these EC types.^27^ These results demonstrate that a similar activation profile of the mechanical response program is found *in vivo* in blood vessels with different levels of blood pressure and oxygen and CO_2_ levels, suggesting that contributions from shear stress and other forces largely contribute to this phenotype (see also **Supplementary** Figure 26).

### Transcriptionally distinct VSMCs in the large arteries

VMSCs are responsible for maintaining vascular tone through pulsatile contraction and production of extracellular matrix proteins. PCA illustrates the spectrum of VSMC transcriptional profiles across tissues (Figure 8a), and we observe that VSMCs from Ao and PA look similar to each other and distinct from other tissues, based on the first two principal components of variation. Subclustering produced a predominant subcluster 0 and a much smaller subcluster 1 (417 nuclei, 2.8% of total VSMC from the vasculature) (Figure 8b). We refer to subcluster 0 as Large artery VSMC (La-VSMC) and subcluster 1 as Venous/Cardiac VSMC (VC-VSMC).

**Figure 8.**
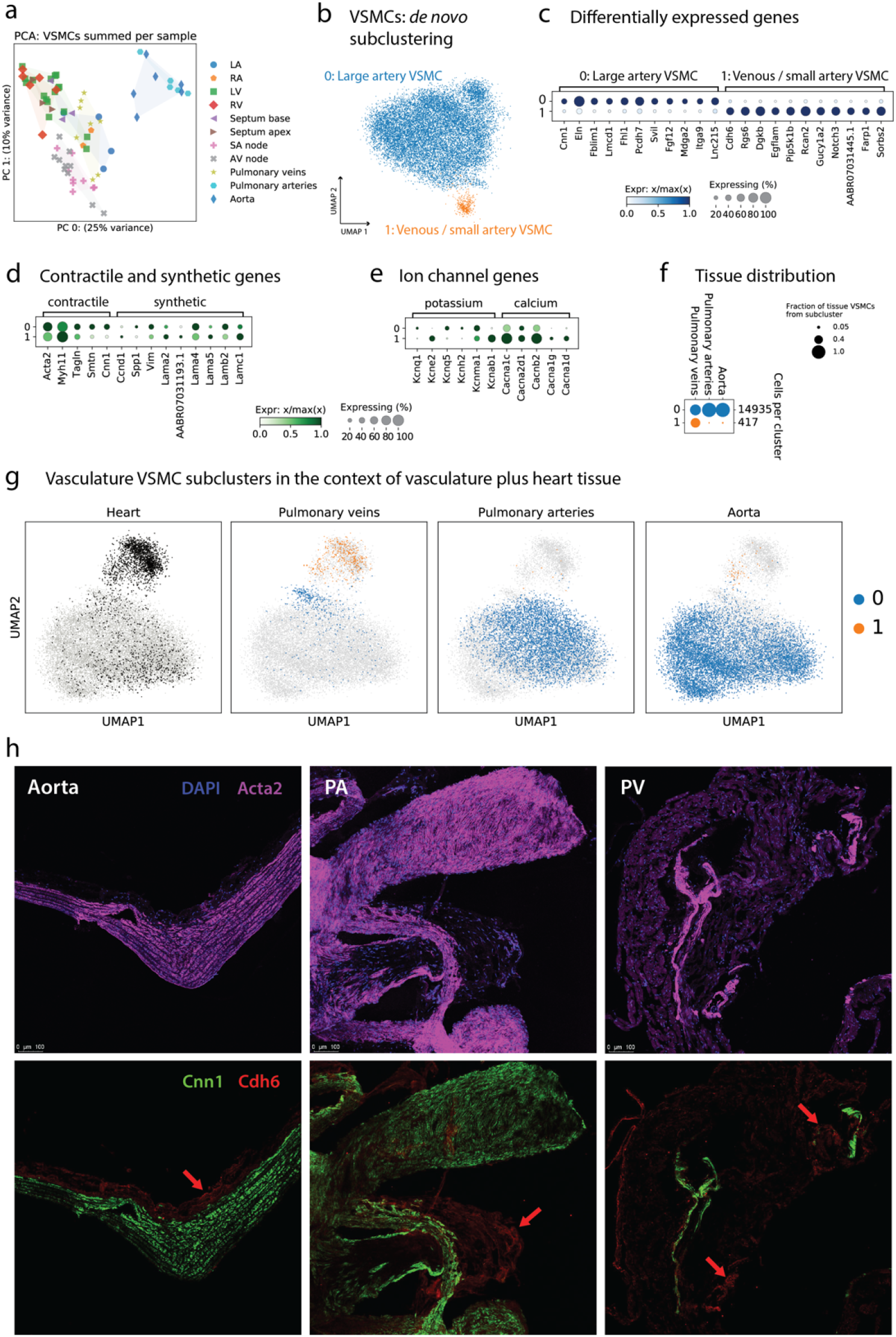
Distinct subtypes of VSMCs in the vasculature. (a) Pseudobulk variation in VSMC transcription. Each dot is the summed expression of all VSMCs from one sample. Coloring by tissue shows that the PA and Ao samples clearly separate from other tissues in the top principal component. **(b)** Subclustering of the VSMCs from PV, PA, and Ao samples reveals 2 subtypes. **(c)** Marker genes of each subcluster show clear distinctions. **(d)** Dotplot showing the expression of a few canonical contractile and synthetic genes in each subcluster. **(e)** Dotplot highlighting a few potassium and calcium channel genes which show different patterns of expression in the two subclusters. **(f)** Distribution of the two VSMC subclusters in PV, PA, and Ao shows that subcluster 1 is nearly absent in PA and Ao. **(g)** UMAPs show all VSMCs from all tissues in light gray, with tissue-specific VSMCs highlighted in black for heart and subcluster color (blue, orange) for PV, PA, and Ao. **(h)** Immunofluorescence imaging of *Acta2* (all VSMCs), *Cnn1* (subcluster 0), and *Cdh6* (subcluster 1) confirms the presence of these cells in PV, PA, and Ao. Subcluster 1 makes up a larger share of VSMCs in PV, in agreement with panel f.

Dotplots in Figure 8c-e quantify the expression of *de novo* subcluster marker genes and canonical genes relevant for VSMC physiology. The subclusters have quite specific marker genes, and additional phenotypically-relevant genes are explored in **Supplementary** Figure 17. La-VSMC have high expression of *Eln* and *Fblim1* among other genes, while VC-VSMC have higher expression of *Ctnna3* and *Rcan2*. VC-VSMC (subcluster 1) is a very small fraction of PA and Ao tissue (Figure 8f). As with the EC analysis above, we used a UMAP to put the vascular VSMCs into the broader context of the entire dataset in Figure 8g, which reveals that the smaller VC-VSMC subcluster is actually the more common VSMC phenotype in myocardial tissue.

As expected based on their biological roles, La-VSMC subcluster 0 shows enrichment of several gene sets related to both cellular and extracellular structural components involved in cell adhesion and tensile strength, while VC-VSMC subcluster 1 shows enrichment of several processes related to ion and transmembrane transporter activity (**Supplementary** Figure 17a; and see the ion channel dotplot in Figure 8e).

We performed immunostaining of PV, PA and Ao sections to validate the presence of these two distinct VSMC subtypes. We identified VSMCs using the general marker *Acta2* (see *Acta2* and *Myh11* in **Supplementary** Figure 17b,e) and co-stained with either Cnn1 antibody for the arterial-enriched La-VSMCs, or Chd6 antibody for the myocardial-enriched VC-VSMCs (Figure 8h). We found abundant double staining of Acta2^+^/Cnn1^+^ cells in both PA and Ao, and fewer such cells in PV (consistent with expectations from snRNA-seq), located mostly in the media and to a lesser degree in the intima layers. Acta2^+^/Cdh6^+^ cells were scarce in all three tissues, and were found mostly in the adventitia layer (Figure 8h, red arrows).

To better characterize the phenotypes of the La-VSMC and VC-VSMC, we focused on several well-defined biological concepts relevant to VSMCs. The large variability in physical forces and environmental cues present in different blood vessels can be linked to a differential expression of contractile and synthetic marker genes (**Supplementary** Figure 17), underpinning a specific cellular state. We did not identify a clear signature for the “synthetic state” of VSMCs (likely consistent with the healthy state of the rats in this study), but we identified a distinctive profile for contractile genes across the two subclusters (Figure 8d). La-VSMCs have higher expression of most contractile markers (*Acta2*, *Tagln*, *Smtn*, *Cnn1*) with the exception of *Myh11*, which is expressed at somewhat higher levels in VC-VSMC. It is well established physiologically that VSMC contraction and proliferation are regulated by potassium channels (Kcn) ^9^. We examined the expression of ion channels to understand whether these two VSMC phenotypes differ in this regard. We observe that *Kcnab1* and *Kcne2* are clearly upregulated in VC-VSMC while several other Kcn genes are upregulated in La-VSMC, but only modestly (Figure 8e). Similarly, the calcium channel genes *Cacnb2*, *Cacna1c*, *Cacna1g* and *Cacna1d* are upregulated in VC-VSMC. Finally, we observe striking differences in the ECM-producing potential of the VSMC subclusters. La-VSMC have a higher expression of *Eln*, and also fibril-forming collagens, beaded-filament collagens, and unique domain organization collagens, while VC-VSMC have higher expression of basement membrane collagens (**Supplementary** Figure 17c).

We refer the reader to the Supplementary Information for similar subclustering analyses of vascular pericytes (**Supplementary Section S.6.6.4**; **Supplementary** Figures 18-19), myocardial mural cells (**Supplementary Section S.6.7**; **Supplementary** Figure 20), vascular FBs (**Supplementary Section S.6.6.2**; **Supplementary** Figures 21-23), and myocardial FBs (**Supplementary Section S.6.4**; **Supplementary** Figure 24).

## Discussion

The rat has been used as a model organism for over a century, and its use for cardiovascular research and disease pathology dates back to at least 1938 ^10^. The rat is a common pharmacological and toxicology model system in the pharmaceutical sector ^11^ due to size, cost, and overall similarity with human physiology ^12^. Here we present an snRNA-seq datasetof over half a million cells from several tissues across the cardiovascular system. We identified cell types and their marker genes at a higher resolution than existing studies in the human heart ^1,3,4,13^, and we inferred a spectrum of phenotypes and sub-phenotypes, as well as cell-cell interactions. We provide an overview of the tissue distribution of multiple cell types, validate the existence of individual populations, and identify differences in gene expression across tissues of the cardiovascular system. Our analyses led to several primary observations.

First, we found unexpected differences in the proportion of individual cell types across tissue regions. We validated these findings using known differences in cellular composition, such as a disproportionate abundance of VSMCs in PA and Ao compared to PV, which is explained by the presence of the intima layer in arterial vessels. We identified a greater proportion of endocardial ECs (EC2) in the atria compared to the ventricles, which may reflect the larger luminar surface in the atria. We also identified a larger proportion of capillary-like ECs (EC1) in the LV compared to the RV, likely reflecting a higher capillary density in the LV of Wistar rats as previously reported ^14^. We found a larger proportion of lymphatic ECs in the ventricles compared to the atria, which could reflect a lower density or smaller diameter in the atria, as seen in dogs ^15^. We also found an enrichment of several neuronal cell types in the PV and nodal regions, representing cell types responsible for the modulation of cardiac function in normal physiology.

Second, we find that several cell subtypes are tissue-specific. We identified nodal pacemaker CMs characterized by canonical markers such as *Hcn4*, *Hcn1*, *Cacna1g*, *Cacna1d*, and *Tbx3* ^16^ as well as *Cpne5,* also seen in mouse ^17^, and novel markers including *Unc13c*, *Mrvi1*, *Robo1*, and *Hs3st3a1*, among many others (Figure 6c; **Supplementary** Figures 10-11). *Hst3st3a1* has been shown to be upregulated in human studies of LA CMs from failing hearts with atrial fibrillation versus non-failing hearts ^18^. Certain genes mark a specific node, with *Shox2*, *Tenm3*, and *Gramd1b* marking the SAN, and *Hmgcll1*, *Abi3bp*, *Arhgap* marking the AVN. We also find that ECs have many discrete, phenotypically distinct cell subtypes, some of which can be found almost exclusively in the arterial or venous blood vessels. Other EC subtypes are mostly found in the heart, with its diverse vascular network (**Supplementary** Figure 14). On the other extreme, FBs exist as a continuum of phenotypes across different tissue niches. An arterial-enriched FB phenotype expresses large amounts of *Col1a2* and *Eln*, required to provide mechanical structure to the PA and Ao, while another population of *Fmod*^+^ FBs resides only in the AVN region. La-VSMCs are enriched in PA and Ao but also present in the heart (vascularized by coronary arteries), while VC-VSMCs are found in the heart and PV but almost absent from the PA and Ao. Increased endopeptidase activity in VC-VSMCs aligns with previous observations of higher *Mmp2/9* and *Timp* activities in rabbit venous versus arterial VSMCs ^19^. Altogether, these results highlight the existence of tissue-specific cellular subtypes which reflect phenotypic plasticity.

Third, we identify several rare cell types. These include *Mpz*^+^ myelinating Schwann cells as well as ganglionic plexi *Apod*^+^ neurons in the PV, which we validated using imaging experiments. We provide a transcriptional profile of *Th*^+^ neuronal cells, a population that consists of only 30 cells in an atlas of over half a million. Since these *Th*^+^ neurons do not express *Npy* but do express *Slc18a2* (vesicular monoamine transporter 2), we speculate that these neurons represent noradrenergic nerve fiber inputs to the heart. We also validated the existence of atrial-like *Nppa*^+^ cardiomyocytes in the ventricles, a finding that is supported by previous work ^20^.

Fourth, we describe predicted cellular communication networks, some of which are differentially enriched by tissue. We identified a strong enhancement of the *Nppa*-*Npr1/2/3* interaction in the SAN between atrial CMs and FBs/ECs, a novel finding that warrants further investigation. *Npr1* and *Npr2* transduce signals from natriuretic peptides through a kinase domain, while *Npr3* does not possess a kinase domain and functions to clear natriuretic peptides from circulation ^21^. The high level of expression of *Nppa* in SAN cardiomyocytes and *Npr1/2/3* in ECs and FBs may suggest an intrinsic ability of the SAN to co-regulate heart rate and blood pressure.

This study was subject to several limitations. First, while the Wistar rat is used as a preclinical model of cardiovascular disease, we should be cautious when translating these findings to human data. However, we do see general agreement between cell types and transcriptional profiles in rat and human heart, and we can use this dataset to determine which cell types are most similar transcriptionally (**Supplementary** Figure 27). Although the Wistar rat is an outbred strain which is more readily translatable to human research than an inbred strain,^30^ both the age of rats (∼17 weeks) and the controlled environmental conditions limit translatability to diverse human populations, where the compound effects of genetic diversity, age, and environmental factors lead to greater transcriptional variability. Second, we characterize the transcriptome using snRNA-seq, where data is derived from nuclear mRNAs at various stages of maturation through the splicing process. Thus, it is unknown how much of a given transcript is spliced and translated (for protein-coding genes). Third, the rat transcriptome is relatively incomplete compared to human or mouse: in this study, we made an attempt to improve the rat reference by combining two data sources as well as our own bulk RNA-seq experiments using rat heart. Another rat reference transcriptome that includes 52,807 genes was recently assembled by others,^31^ yet was not available until after we completed our analyses. Even with our most recent efforts, the state of the rat transcriptome lags behind that of human or mouse.

The analyses and conclusions drawn in this study represent a small fraction of the possible analyses enabled by this rat cardiovascular cell dataset. We continue to explore questions related to novel cell subtypes in PV tissue and the overall distribution of immune cells in the cardiovascular system. The collection of this large dataset under carefully controlled conditions has led to minimal batch effects, and the acquisition of technical replicates and multiple tissue samples per individual rat allows for accurate cross-tissue comparisons. We envision this dataset to be useful not only to the cardiovascular research community and those interested in pharmaceutical development, but also to data scientists pursuing a wide variety of single-cell-related analysis tasks, particularly tasks in need of high quality training data for machine learning applications.

## Methods

### Rat tissue sampling

Animal experiments were approved by the institutional IACUC at Broad Institute. Wistar rats (Charles River, MA) were acclimated for 2-3 weeks to the Broad vivarium, with *ad libitum* access to water and chow diet. 17-week-old animals were euthanized between 10am and 12pm using CO_2_, followed by perfusion with PBS to remove excess blood. Right ventricle, left ventricle and septum were immediately rinsed with ice cold PBS, then minced on ice by sterile razor blade, finally mixed with 4mL of nuclei isolation buffer (NIB). Left atria, right atria, SA node, AV node, aorta (from aortic root to iliac bifurcation), pulmonary vein, and pulmonary artery were immediately frozen in LN_2_ and stored at −80°C until use. Since AV nodes from rat have an approximate volume of 0.5mm^3^, we pooled 3-4 rat samples to obtain 1 sample for further processing. Same approach was used for SA node. See **Supplementary** Figure 28 for details on dissection and validation of *Hcn4*^+^ cells in nodal regions. In total, 89 libraries were prepared and sequenced, of which 78 passed QC (see below). Tissue sampling was performed in such a way that several tissues were collected from each rat. The details of which tissues (and tissue-pools) were collected from each rat are shown in **Supplementary Table 1**. A total of 16 rats were used for snRNA-seq sample collection. In order to retain the transcriptional identity of nuclei, we chose to optimize our nuclei isolation protocol to achieve highly enriched nuclei preparations, to avoid sorting or gradient centrifugation. Cytoplasmic fragments or suspected doublets that may be retained by this strategy were removed during data analysis. A detailed experimental protocol is contained in the Supplementary Methods.

### Single-nucleus RNA-seq

7,000 nuclei input (5,000 calculated recovery) per sample were used for droplet generation and library construction according to the manufacturer’s protocol (10x Genomics, single-cell 3-prime V2 chemistry), with minor modifications (see Supplementary Methods). Libraries were multiplexed at an average of 4-5 libraries per flow cell. Sequencing was performed on an Illumina Nextseq550 in the Broad Institute’s Genomics Platform (genomics.broadinstitute.org).

### Reference transcriptome augmentation

The rat transcriptome from Ensembl (Rattus norvegicus, Rnor_6.0.96) ^22^ lacks full-length *Ttn* as well as large stretches of other important cardiac-related transcripts including *Ryr2*. Many other transcripts are annotated with extents shorter than the read alignment would suggest, resulting in low read-mapping to the Ensembl transcriptome. We therefore created an augmented reference transcriptome for the rat, starting with Ensembl Rnor_6.0.96 as the foundation, which was used for this study. Transcript definitions were augmented in part by performing bulk RNA-seq by isolating poly-adenylated RNA from the aorta, AV node, and all four cardiac chambers of two male Wistar rats and converting it to sequencing-ready Illumina TruSeq libraries. See Supplementary Methods for details. The transcriptome GTF file is available as Supplementary Data. Compared to the Ensembl Rnor_6.0.96 transcriptome, typically 5-10% more reads from cardiac samples mapped to this amended transcriptome.

### Data processing and quality control

Most data analysis was performed using the Terra cloud platform (app.terra.bio). BCL files for all datasets were processed using cellranger mkfastq (CellRanger 3.0.2, 10x Genomics) to demultiplex samples and generate FASTQ files, after first trimming reads (see Supplementary Methods). Quality control at the level of entire samples was performed by examining QC metrics produced by cellranger count, as well as UMAP plots and plots of log(UMI count) versus log(droplet ID) ranked by decreasing UMI count. 11 samples were identified as such strong outliers that they were deemed to be QC failures and subsequently removed (see Supplementary Methods), leaving 78 high quality datasets with approximately 5000 - 10000 nuclei each. Background noise was removed from count matrix data on a per-sample basis using CellBender ^23^, which also performed initial cell calling.

### Clustering

Details are contained in the Supplementary Methods. In brief, extensive droplet quality control was performed for each sample separately using several metrics, in order to eliminate doublets, low-quality nuclei, and cytoplasmic debris. Count matrices for passing nuclei from each sample were aggregated into one large count matrix in scanpy 1.8.2 ^24^. Batch effect correction was performed using scVI 0.6.5 ^25^ with the batch variable being individual rat (or tissue-pool). Batch-corrected latent embeddings of each nucleus from scVI were used to create a two-dimensional map using the UMAP algorithm ^26^. Leiden clustering was run at various resolutions, and the final resolution of 1.2 was chosen manually due to its parsimonious covering of the dataset and its suitability for biological interpretation. Clusters of fewer than 50 cells (there were two such tiny clusters) were excluded from downstream analyses due to their irreproducibility across samples. Cluster 20, which seems to be high mitochondrial contamination with cardiomyocyte and fibroblast signatures, suggesting the presence of remaining doublets after cell QC, was also excluded from further analyses.

### Subclustering

We first narrowed down to a subset of cells, for example, FBs. Additional QC was performed by looking for marker genes of other populous cell types (CMs and ECs) and scoring those gene sets. The top several percent of cells in terms of these scores were eliminated as a very conservative way of eliminating doublets. Subclustering was performed by recreating a neighborhood graph using the cell subset (same as above), and clustering that graph using the Leiden algorithm. UMAP was repeated to create a subcluster visualization, also using this graph.

### Differential expression testing

Differential expression tests were conducted using R limma ^27^. The recommendation of Lun and Marioni was followed ^28^, and so DE tests were performed after (1) summing count data over appropriate groupings (sample or individual rat or sample-by-cluster, etc. depending on the test), (2) normalizing using DESeq2 ^29^, and (3) correcting for the mean-variance trend using voom ^30^. Only genes with summed, DESeq2-normalized counts with a mean of >= 2 were tested. Multiple-testing correction was performed using the Benjamini-Hochberg method.

The matrix plots in **Supplementary** Figure 7e-f are created using the results of a differential expression test as above, and pulling out contrasts for every possible one-versus-one subcluster comparison. The magnitude of the differences in each test can be quantified and distilled down to a single number in a variety of ways, such as the number of genes meeting certain thresholds for p-values and log-fold-changes. However, methods based on arbitrary thresholds are not very robust to changes in those thresholds, and so we chose to quantify the magnitude of differences as the standard deviation of the list of log-fold-changes for each tested gene. This results in a single number which can be used to quantify relative transcriptional differences for the purposes of plotting and visualization.

### Marker gene discovery

Differential expression tests comparing one cluster or subcluster to all others were carried out using the above differential expression testing method. Contrasts of one cell cluster versus all others were fit in limma using the model (∼ 0 + cluster + tissue), along with duplicateCorrelation per individual rat (or tissue-pool), to extract an estimate of a log fold-change between the given cluster and all others. Marker genes shown in the dotplots are a subset of significant results, and are ranked in order of F0.5 score.

### Pathway analyses

Analyses of gene-sets were carried out using gene set enrichment analysis (GSEA), as implemented in the fgsea package in R ^31^. GSEA was carried out using the t-statistic from a relevant differential expression test to rank-order all genes tested. A million permutations were used to calculate a p-value using fgsea.

### Dendrograms and PCA plots

Dendrograms were created in scanpy, using the latent representation from scVI. PCA plots were created by summing expression per relevant cell group, using a variance stabilizing transformation (vst from DESeq in R), and computing PCA on the top 500 highly-variable genes.

### Imaging validation

Marker genes for imaging validation were chosen through a variety of methods, including genes’ known biological significance and differential expression results. Typically, genes were prioritized as potential imaging markers based on having a high positive predictive value for a subcluster of interest.

Frozen tissue was sectioned and mounted on Superfrost slides. After fixation and permeabilization, the samples were stained with RNAscope 4-plex kit reagents, according to manufacturer’s instructions (ACDbio) to validate EC and fibroblast markers or hybridized with primary and secondary labeled antibodies for VSMC validation and *Mpz*^+^ neuron experiments. Refer to the supplementary materials for a full list of reagents. Images were taken with a White Light Laser (WLL) confocal microscope in 3 Z-stacks at 20x (1024x1024 resolution) and further imaged at 63x with immersion oil in 4 Z-stacks to highlight regions of interest (Leica Sp8x).

### Cell-cell communication

Receptor-ligand interactions were inferred based on counts of receptor genes and ligand genes in all pairs of cell types, using the CellPhoneDB database ^5^ and the squidpy 1.0.0 software package’s “ligrec” permutation test ^32^, run separately on each sample, so that only nuclei within the same biological sample were ever tested for cell-cell communication (see **Supplementary Methods**). For tissue comparisons, samples were aggregated by tissue of origin. Differences in mean interaction strengths between tissues were assessed for statistical significance using Wilcoxon rank-sum tests, and p-values were adjusted for multiple testing using a Bonferroni correction.

### Rat and human joint analysis

Rat genes were mapped to human genes as detailed above in the Supplementary Methods. Only those genes which map uniquely from one species to the other were retained (15012 genes, see **Supplementary Table 8**). Human data from Tucker et al. ^1^ were re-processed from the FASTQ files using a pipeline identical to that used for the rat data. Once quality-controlled nuclei from each species were aggregated together, the data were jointly mapped by performing batch-effect correction using scVI with the batch variable being individual. This corrected batch effects due not only to inter-individual variability, but also due to species. Again, counts were not imputed by scVI, but instead the latent representation of each nucleus was used to create a UMAP and perform Leiden clustering with resolution 1.0.

## Abbreviations

| Abbreviation | Definition |
| --- | --- |
| LA | Left atrium of the heart |
| RA | Right atrium of the heart |
| LV | Left ventricle of the heart |
| RV | Right ventricle of the heart |
| SAN | Sinoatrial node of the heart |
| AVN | Atrioventricular node of the heart |
| PA | Pulmonary arteries |
| PV | Pulmonary veins |
| Ao | Aorta |
| FB | Fibroblast |
| EC | Endothelial cell |
| VSMC | Vascular smooth muscle cell |
| La-VSMC | Large artery vascular smooth muscle cell |
| VC-VSMC | Venous/cardiac vascular smooth muscle cell |
| CM | Cardiomyocyte |
| ACM | Atrial cardiomyocyte |
| VCM | Ventricular cardiomyocyte |
| snRNA-seq | Single-nucleus RNA sequencing |
| UMAP | Uniform manifold approximation and projection |
| GSEA | Gene set enrichment analysis |
| GO | Gene ontology |
| PCA | Principal component analysis |

## Acknowledgements

LX is supported by American Heart Association 20CDA35260081. PTE is supported by grants from the National Institutes of Health (1RO1HL092577, 1R01HL157635, 5R01HL139731), from the American Heart Association (18SFRN34230127, 961045), and from the European Union (MAESTRIA 965286). The authors would like to thank Carmen Korch for technical support with animal experiments.

