## Supplementary Information for "Transcriptional profile of the rat cardiovascular system at single cell resolution"

**cardiovascular system at single cell resolution**

#### Supplementary Results

##### S.1 Dataset overview

We performed a comprehensive analysis of the cellular and transcriptional diversity of the rat cardiovascular system, including tissues from 10 regions of the heart and major blood vessels: left ventricle (LV), right ventricle (RV), left atria (LA), right atria (RA), septum, atrioventricular node (AVN), sinoatrial node (SAN), pulmonary vein (PV), pulmonary artery (PA), and aorta (Ao) (**Figure 1a,b**). In total, we collected 89 samples, 78 of which passed quality control (**Supplementary Table 1**). This resulted in a map of 505,835 high quality nuclei. Tissue sampling was performed in such a way that several tissues were collected from each rat. The details of which tissues (and tissue-pools) were collected from each rat are shown in **Supplementary Table 1**. A total of 16 rats were used for snRNA-seq sample collection.

###### S.1.1 Cell types

29 cell clusters were identified by clustering the atlas at Leiden resolution 1.2. Two of these clusters had fewer than 50 cells, and so were not annotated as separate clusters at the global level. One cluster (cluster 20) was entirely excluded from downstream analyses because of the high presence of mitochondrial transcripts, and a high likelihood that it represents CM + FB doublets.

The complexity of each cell type (in terms of number of unique molecular identifiers [UMIs] captured per nucleus) is shown in **Supplementary Figure 1**. The mean number of UMIs per nucleus was 1151, with large variation across cell types, which largely reflects previously observed differences in transcriptional complexity across cell types ^2^. The non-myelinating Schwann cells express more transcripts than any other cell type. Cardiomyocytes are the second-highest ranked, followed closely by adipocytes, all of which express considerably more mRNA than most of the other cell types. Immune cells are among the lowest in terms of total UMIs.

A dendrogram was used to assess the transcriptional similarity of clusters (**Figure 2b**). With the exception of a cluster of *Top2a*^+^ EC, all EC types group together. However, fibroblasts are grouped in two distinct clades: one contains fibroblast clusters FB1, FB2, and FB3, and the other clade groups cluster 25 together with neuronal cells.

Differentially expressed genes were found using limma to compare each cluster versus all others (see Supplementary Methods: Marker gene discovery). Results are shown in **Supplementary Figure 2**, where we highlight the top 5 genes per cluster. A full table can be found in **Supplementary Table 2**.

T-statistics from these tests were used to rank genes for gene set enrichment analysis (GSEA) using gene ontology (GO) biological process terms, the results of which are also shown in **Supplementary Figure 2**. EC types are described by different GO terms, including “endothelium development”, “EC migration”, and “sprouting angiogenesis”. GO terms uniquely label other EC clusters, such as lymphatic EC (Lymph_EC) and *Top2a*^+^ EC. Specifically, the *Top2a*^+^ EC cluster is enriched in pathways involving DNA synthesis and cell cycle progression, suggesting these are cycling/proliferating EC. This finding was corroborated by an enrichment in genes upregulated in the mitotic state in the *Top2a*^+^ EC cluster compared to other cell clusters (**Supplementary Figure 2c**). GO terms also discriminate between VSMC clusters, such as structural and contractile terms, which are enriched in large artery VSMC (VMSC1), while “membrane depolarization” and “calcium channel activity” are enriched in VSMC2 (**Figure 2**). However, for other cell types, GO terms could not discriminate between cell subtypes, as observed for fibroblasts, which are described by common terms such as “ECM” and “collagen”. Of note, the *Apod*^+^ ganglionic plexi neurons in cluster 24 show a unique enrichment in “basement membrane”, consistent with the identification of this cell subtype in the PV (venous tissue) and the SAN, which is localized to the proximal portion of the superior vena cava. Similar GO terms were identified for global makers of ACM and VCM suggesting that genes differentiating these cell types from other cell types in the cardiovascular system may not strongly differ. In the immune cell compartment, we observed differentiating GO terms for T-cells, but overlapping terms for B-cells, macrophages, and other immune cell clusters. Interestingly, the three neuronal cell clusters (Neuro1, Neuro2 and Schwann cells) are described by distinct GO terms.

We further characterized fibroblasts, EC, and mural cells by performing GSEA using genes differentially expressed between cellular subtypes. We examined enrichment of gene sets from additional databases, including KEGG, Biocarta, and PID (**Supplementary Figure 3**). We observed an enrichment for a pathway relevant to “acute myocardial infarction”, as well as “intrinsic prothrombin activation” in fibroblast cluster 1 (FB1) compared to other FB subtypes. However, in fibroblast cluster 2 (FB2), which is more specific to the atria and SA node, we identified pathways specific to “androgen receptor activation”, and a reduced signal from cell adhesion pathway. Finally, fibroblasts localized to the vasculature (FB3) were enriched for Wnt and MAPK signaling pathways, and also sugar and amino acid metabolic processes. A rare subpopulation of fibroblast-like putative ganglionic plexi neuronal cells, cluster 24, displayed significant enrichment for a number of cell adhesion and integrin pathways.

We also identified differences among the mural cells, as well as in the EC compartment. Specifically, VSMC cluster 2 (VSMC2) shows enrichment in calcium and PIP signaling, while VSMC cluster 1 (VSMC1) has increased ECM-receptor interaction and integrin signaling. (ECM, extracellular matrix.) Among the ECs, we observed a signature for pathways relevant to PIP, VEGFR, and KIT in endothelial cluster 1 (EC1). In endothelial cluster 2 (EC2) we found an enrichment in “complement and coagulation cascade”, and Wnt and ribosome pathways enriched in endothelial cluster 3 (EC3). Consistent with the cycling state of cluster 21, the *Top2a*^+^ EC, we found an enrichment in ATR, Aurora B, and PLK1 signaling there. Overall, we identified distinguishing features of the different clusters that describe their cellular function.

###### S.1.2 Tissue distribution of cell types

70-85% of the volume of cardiac tissue is occupied by cardiomyocytes ^33^, yet cardiomyocytes represent 20-25% of cells present in the heart ^34^. Despite the emphasis on cardiomyocytes as the primary cell type present in the cardiovascular tissue, other cell populations, such as fibroblasts, endothelial cells, and rarer cell types such as macrophages and neuronal cells share an important role in regulating cardiac function ^35–37^. Single nucleus transcriptional profiling allows us to disentangle these extensive physical and molecular interactions across a large gradient of different cellular subtypes, and these data broaden our understanding of tissue-specific cell specialization.

We highlight in **Figure 2c** that ECs and FBs are the major cell types across all tissues (with the exception of the PA and Ao, where VSMC1 is a major constituent), and that there are differences in cellular composition across heart regions which are linked to anatomical or functional differences. The ratio of ECs to FBs is higher in the atria compared to ventricles, mostly explained by a greater luminal surface in the atria and large enrichment of endocardial EC. This is consistent with a higher proportion of EC in the atrial tissues, as observed by *in situ* hybridization (**Figure 2f**). In the large vasculature however, the proportion of mural cells (VMSC and pericytes) is larger. VMSCs are as abundant as ECs and FBs in the PA, and VSMCs and are the most abundant cell type in the Ao.

We observed compartmentalization of certain cell types, including: (1) the presence of two cardiomyocyte populations, which map to the atria and ventricles, (2) a large proportion of VSMC1 mapped to the arterial tissues and a more ubiquitous VSMC2, (3) an enrichment of the FB2 cluster in the RA and the embedded/neighboring tissues AVN and SAN, (4) a marked enrichment in FB3 in the large vasculature, and (5) higher pericyte abundance in ventricles and septum, and lower abundance in the large arteries.

From this analysis, we learned that EC2 is enriched in the atrial tissues and the SAN. To validate this finding, we sought to identify marker genes for each EC cluster. We observed ubiquitous expression of *Ptprb* across LA, RA, LV, RV for all EC clusters, and *Flt1* across all but lymphatic EC. We found *Cemip2* and *Bmp6* mostly enriched in the EC2 cluster (**Figure 3c**). Next, we performed RNAscope imaging experiments to validate the presence and distribution of the EC2 cluster on coronal sections of the rat heart. We found that the pan-EC marker *Flt1* was present similarly in atria and ventricles, while EC2 cell markers *Cemip2* and *Bmp6* are mainly localized to atrial tissue in the endocardial regions (**Figure 3d**).

Finally, by combining the information from the GO terms, the marker genes analysis, the top 100 ranked genes per cluster, and the imaging experiments, we defined the EC clusters as follows: capillary EC (EC1), endocardial EC (EC2), large vessel EC (EC3), lymphatic EC (LymphEC), and *Top2a*^+^ EC.

**Supplementary Figure S4** shows regional differences in EC composition between the four chambers of the heart. EC2, which is strongly atrial-enriched according to **Figure 2e**, expresses *Vwf* (shown in **Supplementary Figure S4a**). **Supplementary Figure S4b** shows that *Vwf* is observed in ECs in the atria and hardly at all in ECs in the ventricles. This is consistent with the regional distribution of EC subtypes found from snRNA-seq.

##### S.2 Differential expression by tissue

This dataset provides the opportunity to study the variation in gene expression across tissue regions within each cell type. **Figure 3** and the accompanying **Supplementary Figure 5** explore how the designation of “marker genes” for FBs, ECs, and VSMCs would be different in each tissue studied. We focused our analysis on FB, EC, VSMC; however, this same approach can be applied to other cell types. Limiting ourselves to one cell type and tissue region at a time (for example, FBs in the atria), we perform a statistical test of differentially expressed genes between FBs and all other cell types. Some of the resulting “marker genes” are found ubiquitously (including well characterized FB markers *Gsn* and *Dcn*), while others are tissue-specific.

In order to further understand the tissue-specificity of markers, we perform a second statistical test: for a given cell type, we test differences in expression across tissues (for example, FBs in the atria versus FBs in all other tissues). **Supplementary Figure 5** shows the results of these tests, displaying all the ubiquitous markers at the top, followed by genes which are markers only in one or two tissue regions. We emphasize that all the genes shown are “markers” of the given cell type in at least one tissue (meaning that the genes are significantly enriched in the given cell type compared to all other cell types). Variation in expression across tissues is evident, even among genes which are “markers” of the given cell type. Often, there are genes which are significantly upregulated in the arteries compared to the other tissues, suggesting specialized cellular functions in the vasculature. (Atria: LA and RA. Ventricles: LV and RV. Septum: septum apex and septum base. Nodes: SAN and AVN. Arteries: PA and Ao.)

Some FB marker genes have an obvious tissue-restricted pattern, such as *Csmd1* enriched in the atrial tissue and *Col1a2* enriched in the arteries (**Supplementary Figure 5a**). Among the list of robust marker genes for ECs, we identified *Flt1*, a commonly used EC marker gene, and *Ptprb*. *Nrxn3* is enriched in atrial tissues including the atria, the nodes, and the PV, while *Nav3* is enriched in ventricular tissues, and *Cytl1* in arterial tissues (**Supplementary Figure 5b**). Only *Myh11* and *Dmd* passed our significance thresholds as VSMC markers in all 11 tissues (**Supplementary Figure 5c**). Among VSMCs, *Mill1* and *Il34* have increased ventricular expression, and many other genes including *Fblim1* are enriched in the PA and Ao.

##### S.3 Cell-cell communication

Together with the results presented in connection with **Figure 4**, we explore ligands and receptors expressed in this dataset and their variation in expression across tissues and cell types. **Supplementary Figure 6** shows the top ligands (panels **a** and **b**) and receptors (panels **c** and **d**) that are differentially expressed by tissue (panels **a** and **c**) and by cell type (panels **b** and **d**). Some results are quite familiar, such as the elevated levels of *Nppa* and *Nppb* in ACMs. *Vegfa* and *Igf1* are mostly enriched in VCM. Interestingly, we found clear differences in ligand expression in EC clusters (*Kitlg* in EC1 and *Top2a*^+^ EC; *Vwf* and *Bmp6* in EC2 and EC3), but no clear differences in the FB clusters. Other examples might shed light on interesting mechanisms. For example, the presence of elevated *Notch3* in VSMC2 (cluster 19) as compared to VSMC1 (cluster 10) could shed light on the role of *Notch3* signaling discussed in Ragot *et al.* ^38^. In that work, Ragot *et al.* discuss how *Notch3* deletion is protective in pulmonary hypertension while deleterious in arterial hypertension. Here we observe that the cardiac VSMCs (VSMC2, cluster 19) have high levels of *Notch3* while VSMCs specific to the PA and Ao (VSMC1, cluster 10) have low *Notch3*. This suggests that differences between these two VSMC phenotypes, as well as their tissue localization, could have an impact on the observed outcomes for *Notch3* deletion models.

In addition to the prevalence of receptors and ligands in the dataset, we also show some of the top autocrine (**Supplementary Figure 6e**) and paracrine (panel **f**) interactions in the dataset, as computed from the CellPhoneDB dataset. Only the top interactions are shown.

We highlight the relevance of *Vegfa* from VCM and ACM toward several cell types, such as EC clusters and immune clusters, which receive the signal mainly through *Flt1*, *Npr1* and *Npr3*. Additionally, this analysis shows communication between ACM and many other cell types, including EC1, Lymphatic EC, monocytes, neural cells, and cycling ECs via the *Nppa*-*Npr1* pathway. In **Supplementary Figure 6e**, it is shown that EC2 and EC3 each communicate with themselves via *Bmp6* and its receptors *Bmpr2*/*Bmpr1a*/*Acvr2a*/*Acvr1*. *Bmp6* is known to be expressed in hepatocytes and has systemic functions, including regulating calcium homeostasis and promoting calcification in ECs and VSMCs in atherogenic conditions.^28^ *Bmp6* is expressed in EC2 and EC3 and may have an autocrine role in the local environment, possibly through regulation of angiogenesis.^29^ The full table of cell-cell communication results is included as **Supplementary Table 6**.

In **Figure 4b**, we show several of the top ligand-receptor interactions that are upregulated in specific tissues, prioritizing those interactions which differ most across tissues. We observe tissue-enriched cellular networks, with a particularly high interaction strength in the SAN between *Nppa* and *Npr1*/*2*/*3* receptors. Interestingly, we also identify communication routes specific to the arterial tissues, with a dominant role of the *Bmp6* ligand toward multiple receptors (*Acvr2a*, *Acvr1*, *Acvr2b*, *Bmpr1b*) in the Ao, and with *Wnt5a*-*Ror1* axes being strongest in the PA. The full dataset is included as **Supplementary Table 7**.

##### S.4 Cardiac conduction

###### S.4.1 Combined map of SA node and AV node

The AVN and SAN were carefully dissected after an optimized dissection protocol was arrived at. Details about the dissection of the nodes is included in **Supplementary Figure 28**, where an anatomical image of the dissection is shown. **Supplementary Figure 28b-c** shows immunofluorescence images for AVN and SAN, validating the presence of *Hcn4*^+^ pacemaker cells in the dissected areas.

Given the presence of a nodal-specific population of cardiomyocytes, we generated a UMAP from only the AV and SA nodal tissues to attempt to identify other unique populations of cells. We identified 26 distinct clusters of cells which are generally representative of the global UMAP clusters (**Figure 6a**). This is expected, as when we dissect the nodal regions from the whole heart, a 1mm^3^ section from several rat hearts is pooled (see Methods).

Most of these clusters were evenly distributed across the SAN and AVN, but some were specific, or biased towards a tissue. Brown and white adipocytes (cluster 9 and 14, respectively), as well as mesothelial cells (cluster 15) were predominantly found in the SAN, while ventricular cardiomyocytes (cluster 18), and the *Fmod*^+^ FBs (cluster 13) were found almost exclusively in the AVN (**Figure 6b**). Notably, we identified atrial cardiomyocytes positive for the automaticity associated channels *Hcn1* and *Hcn4* (cluster 16) and these are relatively evenly distributed across the AVN and SAN. *Hcn4* is highly specific to this cluster, while there are some *Hcn1* positive populations of cells in cluster 25 (non-myelinating Schwann cells) and cluster 10 (neurons) (**Supplementary Figure 8a**).

When we compared cluster 16 to cluster 4 (atrial cardiomyocytes), we identified numerous differentially expressed genes. Cluster 16 is defined by *Hcn1* and *Hcn4*, while cluster 4 expresses *Nppa* and *Nppb*, as well as *Ehbp1*, a gene thought to be involved in actin polymerization (**Figure 6c**). GSEA shows that these two clusters have a number of differentially regulated pathways (**Supplementary Figure 8b**), with cluster 16 cells being enriched for smooth muscle contraction, actin polymerization, as well as dilated cardiomyopathy gene sets. Cluster 4 cells were enriched in pathways regulating metabolism, protein translation, and electron transport.

We also explored whether there were any differences in the pacemaker *Hcn4* positive cardiomyocytes between the AVN and SAN (**Supplementary** **Figure 8c**). Genes upregulated in the SAN included *Tenm3*, a neuron connectivity protein, a gene involved in cholesterol homeostasis (*Gramd1b*), a homeobox protein often mutated in atrial fibrillation (*Shox2*), a glutamate receptor interactor (*Grip2*), and a putative neurotropic factor receptor (*Fstl4*). The upregulation of Shox2 in the SAN is consistent with Liu *et al.**^7^* Genes upregulated in AVN pacemaker CMs included actin polymerization (*Arhgap6*), cell-cell contact (*Pcdh7*), and myosin light chain 7 (*Myl7*).

###### S.4.2 AV node

We made separate maps of the AVN (this section) and SAN (next section) as well. The UMAP showing all 56,479 cells from AVN tissue is shown in **Supplementary Figure 10a**, along with counts of cells in each cluster.

Cluster 7 (atrial-like CMs), cluster 13 (ventricular-like CMs), and cluster 14 (pacemaker *Hcn4*+ CMs) are quite interesting, as they represent all three major groups of CMs, and they are cells obtained from the same samples. This allows us to carry out comparisons free from batch effects. **Supplementary Figure 10b** shows the results of such a differential expression test, and identifies genes that are significantly upregulated in the pacemaker CMs in the AVN (as compared to the atrial-like and ventricular-like CMs in the AVN). *Hst3sta1* is the top most upregulated gene, and seems to be novel from the standpoint of literature on marker genes in pacemaker CMs. **Supplementary Figure 10c** represents another way to characterize the differences between pacemaker CMs and other CMs in the AVN. The log fold-change on the y-axis is the same as in panel **b**, but here genes are prioritized in order of the number of gene sets (significant at FDR 0.3) in which the gene was part of the leading edge in the gene set enrichment analysis (GSEA) shown in panel **d**. Light blue bars show the number of upregulated gene sets in which the gene is implicated. This is meant to prioritize genes which are “well characterized”, in the sense that they are members of many annotated gene sets. Here we see more canonical genes like *Hcn1*, *Cacna1d*, and *Hcn4* rising to the top of the list. The difference between panel **b** and **c** suggests that there are several genes of interest in pacemaker CMs which may not yet be well characterized, or gene sets containing relevant functional pathways may be missing. **Supplementary Figure 10d** reveals that the gene sets most upregulated in pacemaker CMs include the KEGG pathways “vascular smooth muscle contraction” and “regulation of actin cytoskeleton”, as well as the GO molecular function term “tropomyosin binding”.

**Supplementary Figure 10e** shows the top 3 marker genes for each cluster in the AVN map. Clusters “23: *Apod*^+^ FB” and “25: *Sorbs1*^+^ FB” comprise the same group of cells labeled “Ganglionic plexi neurons” in the larger dataset.

###### S.4.3 SA node

The map of the SAN shown in **Supplementary Figure 11a** shows all 49,563 nuclei collected from SAN samples. The number of cells from each cluster is shown in **Supplementary Figure 11b**. FBs are the most common cell type, and most closely related to cluster FB2 from the larger map.

As in the AVN, we can do a direct comparison between expression of pacemaker CMs and atrial-like CMs from these samples. The results are shown in **Supplementary Figure 11c**, which highlights genes upregulated in pacemaker CMs, including *Unc13c* (predicted to be involved in glutamatergic synaptic transmission), *Hcn1*, *Cpne5*, *Prdm8*, *Gramd1b*, and *Robo1*, among many others. GSEA was performed to highlight gene sets differentially regulated in pacemaker CMs as compared to atrial CMs, and the results are shown in **Supplementary Figure 11d**, where we find that there is a relative upregulation of many terms having to do with ion channels in the pacemaker CMs: sodium ions in particular (GO “sodium channel activity” and “sodium ion transmembrane transporter activity”), as well as the KEGG pathway “vascular smooth muscle contraction” found in the AVN analysis above.

A dotplot of the top 3 marker genes for each cluster is shown in **Supplementary Figure 11e**. In the SAN, clusters 24 and 25 are the “ganglionic plexi neurons” from the larger map, and here they separate out quite well in the UMAP, suggesting that these cell subtypes are even more distinguishable in the SAN.

##### S.5 Pulmonary veins map

Because the pulmonary veins contain several unique populations of cells (a CM subcluster, a specialized FB subtype) and are also linked to ectopic foci in atrial fibrillation, we clustered just the PV samples to better characterize this unique blood vessel. In the PVs, we identified 36 clusters (**Supplementary Figure 9a**). Certain clusters (“34: Chondrocytes”, “22: Ciliated”, “12: Club”, and “32: FB Lgr5+”) as well as many of the white and brown adipocytes come from sample “PV2” (a and b) (**Supplementary Figure 9b**). The presence of chondrocytes, club cells, and ciliated cells likely indicative of a small amount of lung contamination specific to that “PV2” sample. Of note, PV samples #3 and #4 contain a greater quantity of non-myelinating Schwann cells (cluster 26) and the Tbx20+ FB subtype (“21: FB2 Tbx20+”).

**Supplementary Figure 9c** shows a dotplot of the top 5 marker genes for each of the neuronal-like cell clusters in the PV map (of which there are 8). The accompanying dendrogram shows that clusters 33, 20, and 16 (“Neur”, “Neur1”, and “Neur2”) are the most closely related, and also that the ganglionic plexi neuronal clusters 24 and 25 are closely related. The results of GSEA for GO molecular function pathways in these neuronal-like clusters are plotted in **Supplementary Figure 9d**.

Clusters 24 and 25, the ganglionic plexi neurons, have a cellular identity similar to neurons and fibroblasts, but they cluster separately from these groups. We highlight the differences between these related clusters in the volcano plot in **Supplementary Figure 9e**, which labels top differentially expressed genes. Cluster 24 expresses several genes typically expressed in the brain or nerve tissue (*Apod*, *Chrm3*, *Dclk1*, *Sox6*, *Trpm3*), while cluster 25 expresses genes involved in muscle function (*Utrn*, *Tnni1*) and cell-cell adhesion (*Itgb4*, *Itga6*, *Sorbs1*).

A dotplot of the top 3 marker genes for each cluster in the PV map is shown in **Supplementary Figure 9f**. Chondrocytes were labeled as such due to the significant enrichment of marker genes *Col2a1*, *Acan*, *Sox9*, *Sox5*, *Sox6*, *Col9a3*, *Hapln1*, and *Cytl1* ^39,40^.

##### S.6 Deep dives into specific cell types

The atlas of the entire dataset in **Figure 2a** is clustered at a relatively coarse resolution in order to provide an overview of cell types in the entire dataset. However, there is much detail contained in the dataset that is not visible at the coarse resolution shown in **Figure 2a**. Some of this detail was explored in **Figures 5-8** in the main text. Here we examine additional cell types in further detail by subclustering at higher resolution in order to understand the gene expression and tissue localization of subclusters. In each of the following sections, we focus on one cell type (and potentially a specific tissue region), re-clustering only those nuclei.

###### S.6.1 Cardiomyocytes

The major results from subclustering the CMs is shown in **Figure 5**. Additional information about these CM subclusters is contained in **Supplementary Figure 7**. **Supplementary Figure 7a** shows the same UMAP as **Figure 5b**, but the cells are colored by tissue of origin rather than subcluster identity. The nodal and PV-specific subclusters are highlighted by dotted line ovals. It can be seen that, within the atrial subclusters, there is a split between cells from the RA + SAN + AVN, and cells from the LA. However, the likelihood of this split being caused by a batch effect is mitigated not only by the use of scVI in performing batch effect correction, but also by the experimental design, where LA and RA tissues were collected from the same individual rats at the same time.

The subcluster “4: Atrial 2” is composed almost exclusively of cells from the PV. This also requires ruling out the possibility of a batch effect. In addition to the measures mentioned above (*in silico* batch effect correction, plus experimental design that collected PVs from the same individual rats as RA and LA, along with several other tissues), we observe that some of the CMs from individual PV samples belong to other CM subclusters, especially “5: Atrial 3” (see **Figure 5c**). The fact that not *all* the CMs from the PVs look different from other samples lends credibility to the hypothesis that this separation is not a batch effect.

We further examined the cells in the PV-specific subcluster “4: Atrial 2” via differential expression analysis and RNAscope imaging. **Supplementary Figure 7b** shows a volcano plot highlighting the top differentially expressed genes when “4: Atrial 2” cells are compared to all other CMs from LA and RA samples. The genes *Cntn3*, *Kcnma1*, *Cpeb1*, *Cpeb3*, *Auts2*, and *Cacna1a* are all overexpressed in the PV-specific “4: Atrial 2” subcluster, while *Nppa*, *Fgf14*, *Kcnip4*, and *Bmp10* are all strongly downregulated compared to other atrial CMs. The UMAP in **Supplementary Figure 7c** shows raw counts of *Cacna1a* in all CMs, highlighting the gene’s overexpression in the “4: Atrial 2” subcluster, in agreement with the volcano plot. **Supplementary Figure 7d** displays a single contiguous section of PV and adjoining LA tissue (same section as shown in **Figure 6i**). RNAscope probe for *Cacna1a* is shown in red, while the DAPI counterstain is shown in blue. The enrichment of red *Cacna1a* is visible in the PV compared to the LA. *Cacna1a* expression is specific to CMs in a lookup from the entire atlas, and so we conclude that this *Cacna1a* enrichment in the PV is consistent with the presence of a tissue-specific *Cacna1a*-enriched CM subcluster in the PVs.

**Supplementary Figure 7e-f** shows a heatmap that displays every one-versus-one differential expression test among CMs by subcluster (panel **e**) and by tissue of origin (panel **f**). Lighter color indicates that the two conditions are more similar. The nodal pacemaker CMs appear relatively unique, while the atrial and ventricular clades are visible. Similarly, atrial and ventricular tissues as a whole (still looking only at CM expression) group into separate clades.

We performed an analysis of cell-cell communication involving subclusters of CMs. We find that communication differs by CM subtype (main results in **Figure 5g**). Clusters “18: B cells” and “22: Antigen presenting cells” each communicate with nodal pacemaker CMs via the *Cd74–App* interaction. Atrial CMs communicate with ECs, and ventricular CMs communicate with a mixture of ECs and FBs. Specific interactions reflect *Vegfa* and *Nppa* signaling, mainly between CMs and ECs. Additional plots providing more detail are included in **Supplementary Figure 7g-k**, which show top directed (i.e. one “receptor” and one “ligand” from CellPhoneDB) interactions involving atrial CMs (panel **g**), nodal CMs (panel **h**), ventricular CMs (panel **i**), the PV-specific “4: Atrial 2” CM subcluster (panel **j**), as well as the top undirected interactions (panel **k**). All panels use the same color scale and dot size scale, and so are visually comparable. The lack of enriched interactions in the nodal pacemaker CMs is notable. Among the undirected interactions, the *Nrg1–Erbb4* interaction between ECs and various CM subclusters is ubiquitous.

###### S.6.2 Neuronal cells

In our atlas, there are nearly six thousand cells which have some type of a neuronal identity (**Supplementary** **Figure 12a**). We identified 6 subclusters: the epicardial inputs to the heart (“ganglionic plexi”, subcluster 2), three distinct populations of neurons, including myelinating Schwann cells (positive for Mpz), and a group composed of non-myelinating Schwann cells, and Th positive cells (**Figure 6d**). All 6 of the identified clusters are present in the PV, SAN, and AVN regions (**Figure 6e**). The AVN region has proportionally fewer non-myelinating Schwann cells. The atria and ventricles are generally devoid of neural cells, with the exception of the right atrium (RA) which has neuronal 1 and 2 populations, as well as non-myelinating Schwann cells (**Figure 5d**).

The epicardial sympathetic nervous system inputs (cluster 2) are predominantly marked by transcription factors (*Bnc2*, *Ebf2*), and proteins involved in cell-cell contact (*Tenm2*, *Gpc3*). The main marker for myelinating Schwann cells is *Mpz* (**Supplementary Figure 12d**), alongside the other myelin components *Mbp* and *Prx*. Non-myelinating Schwann cells are defined by *Cntn5*, the neural-specific RNA binding protein *Elavl2*, the small nucleolar RNA *Snhg11*, a member of the fibroblast growth factor family *Fgf14*, and a heparin sulfotransferase *Hs6st3*. *Th*^+^ neuronal cells are defined by hypoxia inducible genes (*Rgs5*, *Higd1c*), synaptic vesicles (*Syt1*), monoamine vesicle transport (*Slc18a1*), dopa decarboxylase (*Ddc*), and carbonic anhydrase (*Car12*). *Apod* is specific to a subset of the epicardial inputs to the heart (**Supplementary Figure 12c**), while *Tenm2* marks those same cells, as well as non-myelinating Schwann cells (**Supplementary Figure 12e**).

In **Supplementary Figure 12f**, we plot the expression of markers of the sympathetic and parasympathetic nervous system. *Th*^+^ cells most strongly expressed two markers of the sympathetic nervous system, while non-myelinating Schwann cells expressed two parasympathetic marker genes (and one sympathetic marker gene). Clusters 0,1,2 are similar in up- and down-regulated gene sets, and this pattern is nearly the opposite seen in clusters 4 and 5 (**Supplementary Figure 12g**).

###### S.6.3 Endothelial cells

In **Supplementary Figure 14** we show a subclustering of all ECs from the study. The UMAP in panel **a** shows that we find 10 *de novo* subclusters of ECs, and this is the same UMAP used to project the ECs from the vasculature in **Figure 7f**. Panels **b** and **c** show the tissue distribution of EC subclusters and their relatedness via a dendrogram, respectively. Panel **d** shows the principal components of variability in pseudo-bulk EC expression across tissues, where we see that the top principal component separates tissues into three similar compartments: PA and Ao; the atria and SAN and PV; and the ventricles and AVN and septum.

The ECs show quite a bit of heterogeneity. This fascinating heterogeneity is consistent with previous reports describing a EC diversity across tissues in other species ^41,42^. **Supplementary Figure 14e** quantifies where the transcriptional heterogeneity is coming from, and explores whether tissue of origin or EC subcluster has a larger effect. PCA shows the transcriptional variability when data are pseudo-bulked by (sample, subcluster). When dots (sample, subcluster) are colored by tissue (right panel), we see that tissues do not separate from one another. However, when colored by EC subcluster, we see that subcluster identity explains nearly all of the transcriptional variability we see in the first two principal components.

Marker genes for the EC subclusters are shown in **Supplementary Figure 14f**, and their similarity by differential expression testing is shown in panel **g**. Panel **i** explores GO biological process terms enriched in each EC subcluster (compared to all other EC subclusters). Finally, **Supplementary Figure 14h** displays the usage of EC subclusters in each tissue visually on UMAPs. Differences between tissues are quite striking in terms of EC subtype composition.

###### S.6.4 Fibroblasts

We find that FBs display a broad continuum of transcriptional profiles. FBs are highly plastic cells that during normal physiology contribute to many functions, and adapt and specialize according to the specific tissue environment ^43^. Here we explore the FBs from heart tissue, excluding PV, PA, and Ao, which are examined separately in the section on cells of the vasculature.

We collected nearly 142,000 FB nuclei from heart tissue (**Supplementary Figure 24a**) and these can be made to subcluster into 4 main types (panels **b** and **c**). Marker genes are shown in **Supplementary Figure 24d**, and we see that marker genes specific to subclusters 0 and 1 are hard to find, as the distinction between 0 and 1 is a gradual gradient of transcription. Panel **e** shows that subcluster 3 is the most different from the others. It matches the main cluster of “*Fmod*+ FBs” from the entire atlas, and we see that these FBs are coming almost entirely from the AVN region (panel **c**). **Supplementary Figure 24f** shows that subcluster 3 FBs express *Postn*, and may represent an activated state. In panel **g**, we compute differentially expressed genes among FBs (any subcluster) between the atria and ventricles. In the atrial FBs, we see a relative upregulation of *Ccn1*, *Ccn2*, *Ccn5*, *Fos*, *Alpl*, *Epas1*, and *Egr1*, among other genes, and we see a relative downregulation of L ribosomal genes (*Rpl41*, *Rpl15*, *Rpl37a*) and *Atp5f1b*. In panel **h**, we explore differences in FB expression between the right and left chambers of the heart, and find that *Plcxd3* is upregulated on the right-hand side, while *Ntrk3* is upregulated on the left-hand side. We also explore gene sets enriched in each tissue among all FBs (**Supplementary Figure 24i**) and gene sets enriched in each FB subcluster (**Supplementary Figure 24j**). We see that the KEGG_RIBOSOME pathway is upregulated in ventricular tissues and downregulated in atrial tissues, in agreement with the above differential expression tests.

###### S.6.5 Immune cells

In the main atlas, we found that immune clusters (macrophages, monocytes, T cells, B cells, and antigen presenting cells) were quite evenly distributed across tissues, with some enrichment of B cells in the ventricular tissues and septum compared to atrial tissues. Here we take a deeper dive into the immune cells, and subcluster them separately to achieve finer resolution. In total, we explore over 61,000 immune cells from across the cardiovascular system.

**Supplementary Figure 25b** shows that, at the pseudo-bulk level, immune cells look different in the Ao and in the PA than they do in other tissue regions, which largely look rather similar. A *de novo* subclustering of all the immune cells is shown in **Supplementary Figure 25c**, where we have enough resolution to identify 13 clusters. The tissue distribution of these subclusters is shown in panel **d**. We confirm that B cells are slightly enriched in ventricular tissues as well as the PV. Many other cell types are relatively uniformly present across tissues. Cycling T cells (subclsuster 12) are absent from the PA and Ao, but present in all other tissues. Subcluster 8 (*Cd8a*+ T cells) are present chiefly in the LV and the SAN, with some additionally in the AVN and the vasculature (PV, PA, and Ao). **Supplementary Figure 25e** shows gene sets enriched in each immune subcluster, while panel **f** shows the top 3 marker genes of each cluster. We see that the subclusters have clear and distinct markers.

###### S.6.6 Cells of the vasculature

**Supplementary Figure 13** provides an overview of all the cells collected from the vascular tissues: PV, PA, and Ao. The vascular system accomplishes multiple physiological functions to maintain organismal homeostasis: it provides molecules necessary for cell and tissue function, enables intra- and inter-organ communication, distributes circulating cells to peripheral tissues, and plays a key role in thermoregulation. Blood vessels are biologically diverse, due to both a complex developmental origin and different physiological contexts. ^44,45^ The vascular lumen is composed mainly of endothelial cells (EC), which sense and respond to both mechanical and chemical cues, such as shear stress and oxygen availability ^46,47^. The medial part of most blood vessels has a key role in the regulation of vascular tone, and compositional differences in both extracellular matrix (ECM) and the abundance of vascular smooth muscle cells (VSMC) differentiate small and large blood vessels ^48,49^. The composition of the adventitia, the outer layer of a blood vessel, is far more complex, consisting mostly of fibroblasts, pericytes, immune and mesenchymal cells ^50^.

Cell physiology across vascular beds is heterogeneous, and this is partially explained by different genetic and epigenetic programs or because of adaptation to specific molecular cues ^44^. The pulmonary circulation responds to hypoxic conditions by vasodilation, while most blood vessels respond by long-term constriction; and calcification of VSMC is heterogeneous across blood vessels ^51,52^. For instance, certain cellular phenotypes are conserved *ex vivo*, as observed in human coronary artery EC compared to saphenous vein EC after stimulation with oxLDL ^53^. As postulated by Florey in 1966, there are tissue-intrinsic behaviors that may be explained on the basis of differing cellular composition, or the existence of different cellular phenotypes and their functions ^54^. However, a molecular characterization of the cell types and discrete phenotypes across a panel of large blood vessels is incomplete, and warrants investigation at single cell resolution.

Recently, single-cell RNA-seq (scRNA-seq) studies have dramatically improved our understanding of cellular diversity. Initial scRNA-seq studies of mouse aortic tissue described a cellular landscape of ~10 major cell types ^55,56^. A similar experimental approach has been used to define cell types from aortas of pig, monkey, and human, followed by the identification of 10-12 cell types ^57–59^. These early discoveries suggest that biological heterogeneity could be far more complex than currently described. We completed a methodologically refined large-scale analysis of the cardiovascular system of the rat at single cell resolution and obtained a high-quality dataset that we used to explore cell types and cell-cell communication across cells and tissues. We leveraged this information to perform a systematic analysis of the large arterial and venous beds. Our aim was to identify the cell types of the large vasculature and characterize cell types and subtypes that are intrinsically linked to large blood vessel physiology. These analyses further our understanding of blood vessel biology, enabling better modeling of the cellular context for human disease and a more precise interpretation of preclinical studies.

An overview of extracellular-matrix and oxygen response and metabolic genes in the vasculature at the bulk tissue level is shown in **Supplementary Figure 26**.

###### S.6.6.1 Vascular endothelial cells

ECs specific to the vasculature were characterized in **Figure 7**. Co-expression of the *Klf2* and *Nf-kb* program in EC subclusters is restricted to the large vasculature. The *de novo* subclustering analysis is shown in **Figure 7b**, identifying 7 EC subclusters from the large blood vessels. We defined subcluster names/types based on tissue distribution, highly expressed genes and variable genes. We assigned the following nomenclature: capillary-EC (Cap-EC), Large artery-EC (La-EC), Large vein-EC (Lv-EC), Cycling EC (Cyc-EC) and Neuronal/Mesenchymal-like-EC (Nml-EC). In Cap-EC we observe higher expression of *Myo10* and *Dach1*, genes involved in EC migration ^60,61^. La-EC is distinguished by the expression of *Meis1* and *Eya4*, key regulators of cell proliferation and differentiation ^62^. In Lv-EC we found high expression of *Slit3*, a secreted pro-angiogenic and migratory molecule ^63^. Lymph-EC marker genes comprise canonical markers for lymphatic vessels, such as *Reln* and *Ccl21*. Among the small subclusters, Cyc-EC were distinguished by the presence of *Top2a* and a large number of genes related to cell cycle (not shown), while Nml-EC were distinguished by a wealth of genes related to neuronal functions and stemness, such as (*Lsamp*, *Grid2*, *Syt1*, *Msi2*).

Of note, Cap-EC were relatively abundant in PV (**Figure 7e**), a distribution that resembles EC from the heart, a tissue with a dense capillary network (**Figure 7f** panel 1 *vs* 2). We also noted that capillary EC heterogeneity is reduced in the large blood vessels. As expected, the large vasculature does not include endocardial EC, which provides some validation of our subclustering strategy. In agreement with a previous study, we also found a lack of lymph-EC heterogeneity across tissues ^64^. The large vasculature and the heart have overlapping EC subtypes, but also have tissue-restricted EC types that likely have functional implications for cardiac and vascular physiology.

We also validated the identity of these EC subtypes in the PV and Ao using RNAscope (**Supplementary Figure 15**). We imaged Cap-EC as *Flt1*+/*Ptprb*+/*Vwf*- cells, La-EC as *Ptprb*+/*Vwf*+/*Flt1*-, and Lv-EC as *Ptprb*+/*Vwf*+/*Flt1*+. We identified Cap-EC present mostly in PV and to a lesser degree in Ao (**Supplementary Figure 15b,d**). This could reflect regional anatomical promiscuity between PV and LA, but most likely reflects a higher density of vasa vasorum network in venous versus arterial tissue ^65^. We also successfully identified Lv-EC from the PV (but also in Ao to a lesser degree) mostly present on the luminal endothelium **(Supplementary Figure 15a-d**). However, using this selected probe combination we observed that subcluster 1 (*Ptprb*+/*Vwf*+/*Flt1*- for La-EC) was exceedingly rare, as *Flt1* was present in most ECs. Thus, this probe combination cannot discriminate between La-EC and Lv-EC, which are probably present as a continuum of phenotypes. A more inclusive probe set (>3) may allow a precise discrimination of these clusters.

Finally, we explored the phenotype of these subclusters, focusing on defined physiological responses: mechanical stress, activation state, and oxygen response and metabolism (**Supplementary Figure 16**). ECs respond to laminar shear stress by an adaptive transcriptional program, which includes the expression of *Klf2*, Klf2-regulated genes *Thbd* and *Nos3*, and other genes previously shown to respond to mechanical stimulation, such as *Klf4* and its target *Ass1* ^66,67^. We identified a higher expression of *Klf2*, *Klf4*, and *Ass1*, mirrored by lower expression of *Dach1*, *Cxcl12*, and *Angpt2*, indicative of activity of a mechanical stress program in La-EC and Lv-EC subclusters (**Supplementary Figure 16a**). We also observed a higher expression of EC activation markers *Vcam1*, *Icam1*, and *Vwf* in La-EC and Lv-EC, the same EC types where we identified higher *Klf2* and *Klf4* (**Supplementary Figure 16b**) ^67,68^.

###### S.6.6.2 Vascular fibroblasts

PCA of pseudo-bulk FB expression in each sample reveals that Ao and PA FBs clearly segregate from all other cardiac FCs along the top principal component (PC1) (**Supplementary Figure 21a**). Contrary to the EC observation (**Figure 7a**), FBs from the PV occupy an intermediate location along PC1, between arterial and cardiac FBs.

A *de novo* subclustering analysis identified 4 FB subclusters which show up as rather continuous in the UMAP representation (**Supplementary Figure 21b**). This suggests that during normal physiology, fibroblasts in the large vasculature probably display a continuum of phenotypes, instead of discrete, highly specialized phenotypes. The per-subcluster composition of PV, PA, and Ao show differences in the rank order of these fibroblast subclusters, providing further evidence for the role of cell-environment interaction (**Supplementary Figure 21h**). One example is a higher proportion of Fib_s0 in Ao and PA, while Fib_s1 is the most abundant subcluster in PV. From the subcluster dendrogram, we find that fibroblasts from subclusters 0 and 1 are very similar, while the small subcluster 3 (266 nuclei) is more distant and in a separate clade (**Supplementary Figure 21c**). Similarly, differential expression tests between subclusters highlight a large transcriptional difference between subcluster 3 and other fibroblast subclusters (**Supplementary Figure 23a**). The expression of several extracellular matrix genes and signaling genes are shown in **Supplementary Figure 21f-g**.

Fibroblasts from the heart largely match the distribution of fibroblasts from all rat cardiovascular tissues on a UMAP, suggesting that the heart encompasses most identified phenotypes (**Supplementary Figure 21i**, panel 1). However, we observe that large artery fibroblasts (Fib_s0) are transcriptionally homogeneous and cluster in a smaller region of the UMAP (**Supplementary Figure 21i,** panels 3 and 4). PV fibroblasts are transcriptionally more heterogeneous, with a higher proportion of Fib_s1, which overlaps with heart fibroblasts (**Figure 7h,** panel 2, left region of the UMAP). From the analysis of the entire cardiovascular dataset, we identified a larger proportion of Fib_s3 in atrial tissue (data not shown), a tissue adjacent to the PV. While Fib_s3 looks rare in the context of the PV, PA, and Ao, it is actually common in the heart (**Supplementary Figure 21i**, panel 2 red cells compared to panel 1).

A differential expression analysis is shown in **Supplementary Figure 21d**. Although Fib_s0 and Fib_s1 were transcriptionally distinguishable from the rest, we could not identify highly specific marker genes for Fib_s0, apart from a higher expression of *Eln*, reinforcing the observation that FB subclusters display a continuum of phenotypes, in contrast to EC subclusters. We instead identify marker genes for the other subclusters, such as *Smoc2*, a regulator of fibroblast-to-myofibroblast transition ^69^ in Fib_s1, and the Mmp2 inhibitor *Pi16* ^70^ in Fib_s2. Interestingly, we observed more distinctive marker genes for Fib_s3, a subcluster with higher levels of *Tbx20*, *Smad6*, *Nr4a1*, and *Ccbe1*, genes that either drive tissue repair or act as inhibitors of TGFb1 signaling.

Next, we used RNAscope to validate the presence of FB subclusters in both PV and Ao (**Supplementary Figure 22**). Pdgfra has been found to be a marker for most FBs ^71^, and appears to be a marker for FBs in our data (see **Supplementary Figure 21e**). We imaged Fib_s0+s1 as *Pdgfra*+/*Adamts16+* cells, and Fib_s2 as *Pdgfra*+/*Pappa1+ or Pdgfra+/Tspan11+*. Based on our analysis, discrimination of clusters 0 and 1 would be achievable only by inclusion of additional markers. We performed imaging at both 20x and 63x resolution to observe general localization of the marker genes across large tissue sections, and also assess the presence of given combinations of marker genes in cells. As expected, we observed enrichment of FBs in the adventitia of PV and Ao. We also validated the presence of subcluster 0+1 and 2 in both tissues (**Supplementary Figure 21b,d**). Interestingly, we observed a high dynamic range of *Pdgfra* and *Tspan11* in fib_s0+s1, which could represent the existence of several cell states.

We also assessed the expression of genes and gene programs to explore biological roles of fibroblast subclusters. We performed GSEA (**Supplementary Figure 23b**) and identified increased expression of the NPP1 pathway, responsible for bone mineralization, in Fib_s0. A cytokine-receptor and integrin signaling signature is statistically higher in Fib_s1, sugar and glycan biosynthesis signature in Fib_s2, and nitrogen metabolism in Fib_s3. An *ad-hoc* analysis of key biological functions of fibroblasts, including collagens, non-collagen and ECM remodeler genes, highlights a higher expression of *Eln* in Fib_s0, *Col1a1* and *Col3a1* in Fib_s1, *Fn1* and *Fbn1* in Fib_s2, and *Lamc1* in Fib_s3 (**Supplementary Figure 23c-e**). A small overview of metalloproteinases and their inhibitors yielded a limited number of genes with reasonable expression in our dataset. Of note, we found a higher expression of *Mmp2* in Fib_s1 and a lower expression of *Timp3* in Fib_s2 (**Supplementary Figure 23e**). These results suggest that different subtypes of fibroblasts make unique contributions to the synthesis of ECM in the large vasculature. We also analyzed the gene expression signature of the Tgfb and Bmp pathways, which play a major role in the specification of the fibroblast phenotypes. Our analysis uncovered a very complex scenario, whereby *Tgfbr2* is expressed at higher levels in Fib_s1 and Fib_s2, while the Tgfb-pathway transcription factors *Smad2* and *Smad3* are more expressed in Fib_s3, though the differences are rather subtle (**Supplementary Figure 21g**). Our data indicate that the Smad1/5/8 pathway may be less active in Fib_s0, inferred by a lower expression of *Bmpr1a*, *Bmpr2* and *Smad1*.

###### S.6.6.3 Vascular smooth muscle cells

We find that VSMCs exhibit transcriptional differences between large arteries and other tissues (**Figure 8**). VMSCs are responsible for maintaining the vascular tone through pulsatile contraction and production of ECM proteins. As expected from their biological roles, La-VSMCs (subcluster 0) show enrichment of several pathways related to both cellular and extracellular structural components that have a role in cell adhesion and tensile strength, while VC-VSMCs (subcluster 1) show enrichment of several processes related to ion and transmembrane transporter activity (**Supplementary Figure 17a**). In panels **b** and **d**, we show the expression of contractile, synthetic, and mechanical response genes at the pseudo-bulk tissue level for VSMCs. In addition to the VSMC subcluster imaging validation shown in **Figure 8h**, we performed immunostaining of Myh11 and Acta2 on tissue sections from PV, PA, and Ao. Those results are shown in **Supplementary Figure 17e**, and we can see that Myh11 and Acta2 colocalize in VSMCs in all three tissues.

Current knowledge supports heterogeneity and plasticity in the VSMC compartment, which reflects both diverse embryonic origin and the exposure to different environmental cues ^72^. In this study, we identified two discrete VSMC subclusters with marked transcriptional differences and a different distribution across blood vessels. In the arterial tissues, the La-VSMC phenotype (*Cnn1^+^/Cdh6^-^*) was dominant, while VC-VSMC (*Cdh6^+^/Cnn1^-^*) was enriched in the PV. La-VSMC are phenotypically distinct, with a higher expression of contractile genes. In addition, we observed a differential distribution of ion channels across the VSMC subclusters, suggesting a different regulation of VSMC physiology in the large arteries compared to PV or cardiac tissue. It has previously been described that *Kcnq5* is expressed in smooth muscle cells of the mesenteric artery, a blood vessel of large diameter ^73^. We observed expression of *Kcnq5* in the La-VSMC (yet almost zero expression was detected in VC-VSMC), suggesting a role for *Kcnq5* in large arteries across the body. Conversely, in the VC-VSMC we found a clear enrichment in several subunits of the L-type and T-type subunits of calcium voltage-gated channels, regulators of VSMC excitation, contraction, transcription, and proliferation ^74^.

###### S.6.6.4 Vascular pericytes

We find that pericytes in *vasa vasorum* differ from those in the heart. Pericytes have a complex physiological role, acting as pseudo-VSMCs and nourishing and protecting the integrity of small blood vessels by regulating EC physiology. We use PCA to illustrate the spectrum of pericyte transcriptional profiles across tissues of the cardiovascular system, and we observe that pericytes from Ao and PA segregate to the top right of the plot, differing in both of the top two principal components (**Supplementary Figure 18a**). Although it is known that pericytes from different tissues may have different physiological roles, it is not known whether pericytes in the large blood vessels are transcriptionally heterogeneous or play roles beyond controlling small blood vessels (*vasa vasorum*).

We performed *de novo* subclustering of 2,087 pericyte nuclei from the PV, PA, and Ao, and we identified two pericyte subclusters, a larger Peri0 and Peri1 (~7% of pericytes) (**Supplementary Figure 18b**). In the PV, the proportion of Peri0 is much larger than Peri1, while Peri0 and Peri1 have a nearly equal representation in Ao (**Supplementary Figure 18e**). The PA is intermediate. We then asked whether these two pericyte subclusters are unique to the large vasculature or if they represent common cell phenotypes identified in the cardiovascular system. Combining nuclei of pericytes from the heart and from the large vasculature in a single UMAP plot, we highlight that the distribution of pericytes from the rat heart (black dots) overlap the entire global cardiovascular pericyte (**Supplementary Figure 18f**, panel 1), while pericytes from the PA and Ao have a more restricted distribution (**Supplementary Figure 18f**, panel 2-4). Conversely, PV pericytes comprise both subclusters, with more of Peri0 (**Figure 6e**, panels 2-4). Interestingly, we also observed an enrichment of Peri1 in SAN tissue (shown as subcluster 3 in **Supplementary Figure 20d**), which contains a portion of the superior vena cava. We suggest that Peri1 is a large blood vessel pericyte cluster, probably derived from the *vasa vasorum* network in these tissues, and that Peri0 is heart pericytes.

GSEA identifies “passive transmembrane transporter activity” as the pathway enriched in Peri1 (**Supplementary Figure 19a**). A marker gene analysis reveals differences in the transcriptional profiles of the two subclusters (**Supplementary Figure 18c**). In Peri0 there is an enrichment in membrane structural and communication proteins (*Lama2*, *Pid1*, *Ank3*, *Ctnna3*), and in Peri1 we identified phosphodiesterase (Pde) enzymes (*Pde4d*, *Pde10a*) (**Supplementary Figure 18d**). Previous work suggests that phosphodiesterase family members are tissue-specific and that some Pde isoforms are specifically expressed in rat brain pericytes ^75^. We find a relative over-expression of *Pde3a*, *Pde1a*, *Pde7b* in Peri0 and of *Pde4d*, *Pde10a*, *Pde5a* in Peri1 (**Supplementary Figure 18d**). Among these genes, *Pde3a, Pde4d*, and *Pde10a* show strong enrichment (see **Supplementary Figure 19b** for UMAPs). The enrichment of different Pde isoforms in the two pericyte phenotypes suggests different biochemical events in these cells, which may help elucidate the behavior of these cells in the response to extracellular stimuli ^76^.

Pericytes have pleiotropic roles in vascular biology, supporting vasomotion, angiogenesis and neovascular stabilization, basement membrane homeostasis, and the regulation of the endothelial barrier ^77^. It is known that pericytes are heterogenous across organs ^78^, however it is currently unclear whether pericytes acquire different phenotypes in a defined tissue. In the human heart, a single pericyte cluster has been previously identified ^78^, and in the brain, there is no conclusive evidence to support the existence of pericyte heterogeneity ^79,80^. Here, based on all 26,796 pericytes in our atlas, we identified two pericyte subclusters (**Supplementary Figure 18f**), which suggest a rather limited heterogeneity in this cell compartment. Both identified pericyte types express the general pericyte markers *Rgs5*, *Abcc9*, and *Plcl1*. Interestingly, we observed that the large arteries are composed almost exclusively of Peri1. This subcluster is characterized by expression of *Pde4d*, a gene not previously identified in cerebral pericytes ^75^. From a broader analysis of phosphodiesterases, we observed a skewed expression of several genes across pericyte types, which suggests that functional specialization of pericytes differs across tissues.

###### S.6.7 Mural cells

We additionally sub-clustered all the mural cells (VSMCs plus pericytes) from all tissues excluding the large blood vessels PV, PA, and Ao (over 27,000 cells; see **Supplementary Figure 20a**). *De novo* subclustering revealed four subclusters: the same two VSMC subclusters and two pericyte subclusters found in the preceding sections. **Supplementary Figure 20h** lends additional evidence that “subcluster 3” here (which corresponds to Peri1, the pericytes in *vasa vasorum*) is present in the SAN region due to some unintended sampling of the superior vena cava (mentioned above). We can see from the breakdown of cells by sample that only the samples from tissue pool 2 (and not pools 1 or 3) contained this group of cells.

##### S.8 Rat and human transcriptional similarities

An important question raised by this dataset is translatability to human biology, both in terms of knowledge as well as use in preclinical models of disease. We attempted to address the validity of the rat as a model organism for human cardiovascular research by comparing orthologous genes across our healthy rat dataset and a published human snRNA-seq atlas of the four chambers of the human heart ^1^. We re-ran all sequencing data through our most recent analysis pipeline, followed by a combined clustering of rat and human nuclei (**Supplementary Figure 27a**). The result was the generation of a comprehensive UMAP of 19 cell types, which describes overall similarities across the rat and human cellular composition (**Supplementary Figure 27b**). Each individual cluster from the UMAP is composed of both rat and human nuclei. We compared the distribution of cell types across the two species (**Supplementary Figure 27c,d**), and identified overall similarities in the proportion of certain cell types (e.g. FBs, pericytes, macrophages and neuronal 2), and species-specific enrichment of cell types such as VSMCs, lymphatic ECs, and adipocytes. The differences in lymphoid cell distribution may reflect both biological divergence and perfusion at the time of tissue collection.

Beyond this initial analysis, we attempted to describe the conservation of molecular features across species in each cell type (**Supplementary Figure 27e**). Globally, we observed that most cell types have conserved molecular function pathways (GO molecular function gene sets) in both species. We find an exception in selected molecular pathways in immune cells, which may reflect differences in rat and human immune cell biology ^12,81^. Finally, we studied conservation of the transcriptional landscape in each cell type by calculating a correlation matrix using the pseudo-bulk expression of each cell type (**Supplementary Figure 27f**). We included the top 2,000 most variable genes, which are a close representation of cell type identity. This approach identified similarities (*r*>0.5) across specific cell types in rat and human. We highlight high levels of transcriptional conservation in ECs, neuronal cells, ventricular CMs, and T cells. The transcriptional profiles of adipocytes are the least conserved across species, and this reflects the underlying biological heterogeneity in mouse and rat adipocytes (white, brown, beige) as opposed to a larger pool of white adipocytes in humans ^82^. Overall, this analysis exemplifies similarities in cardiac tissue composition that go beyond overall anatomy, with widespread conservation of molecular functions and transcriptional signatures across many prevalent cell types.

##### S.9 Removal of noise in snRNA-seq data

Removal of background noise (ambient RNA) was important for cleaning up the transcriptional profiles of cells in this dataset. We used CellBender remove-background ^23^ to remove contaminant ambient RNA. The difference in transcriptional profiles of cells before and after cleanup is visualized in the dotplot in **Supplementary Figure 29**. The dotplot of the raw data shows a low level of expression of the FB marker genes Gsn and Dcn, as well as the CM marker genes Ttn and Ryr2, in all cell types. However, after running CellBender, much of this non-specific expression has been removed.

##### S.10 Concluding remarks

This rat cardiovascular atlas features systematic tissue sampling from different regions of the cardiovascular system. Bulk RNA-seq experiments have long demonstrated that transcription varies widely based on tissue of origin. Single-cell experimental techniques have revealed that differing cell-type proportions across tissues can explain much of this transcriptional variability. However, cell-type proportions are not the only source of transcriptional variability. Not only can cell-type composition vary, but also the expression profile of a given cell type can vary in different tissue contexts. The extent to which the variability of expression within a single cell type is explained by differing proportions of cell subtypes with fixed expression profiles, versus a situation where a cell type simply has a fluid expression profile in different tissue contexts, has not been thoroughly explored. Our dataset sheds some light on this matter. ECs seem to be a good example of the first scenario: cell subtypes with well-defined expression profiles that are mixed in different ratios in different tissues (**Supplementary Figure 14**); while FBs seem to be an example of the second scenario, as their transcriptional profiles are quite continuous, and they seem to vary rather subtly in different tissue contexts (**Supplementary Figures 21-24**).

#### Supplementary Methods

##### Nuclei isolation and single-nucleus RNA-seq

Frozen tissues were mounted on OCT and sectioned at 60μm at -20°C using a cryotome (Leica CM 1950). Tissue slices were transferred to 4mL of ice-cold NIB (Hepes 20 mM, Sucrose 0.25M, MgCl_2_ 3mM, KCl 25mM, Igepal-630 0.01%, BSA 0.5%, pH 7.2). All tissue mixtures were *dounce* homogenized on ice with 10 strokes of a loose fit pestle followed by 10 minutes incubation, followed by 10 strokes of a tight fit pestle and 10 min incubation. Homogenate was transferred to a 15mL conical tube, filled to 8mL with *nuclei wash buffer* (NWB = NIB without detergent), and centrifuged at 40*g* x 4’, at 4°C (Beckman Coulter Allegra X-15R swinging bucket centrifuge). Supernatant was filtered through sequential 40μm and 10μm meshes (Pluriselect, Germany) in 50mL conical tube, filtrate topped up to 10mL with NWB and centrifuged at 600*g* x 5’, at 4°C. Supernatant was discarded and pellet resuspended, transferred to 15mL tube, topped up with 8mL NWB centrifuged (600g x 5’, 4°C). Final pellet was resuspended in 150μL *nuclei resuspension buffer* (NRB = NWB with 1:80 murine RNAse inhibitor, NEB). All procedures were performed on ice. Nuclei, stained with Trypan blue, were manually counted using a hemocytometer (inCyto.com). 7,000 nuclei input (5,000 calculated recovery) per sample were used for droplet generation and library construction according to the manufacturer's protocol (10x Genomics, single-cell 3-prime V2 chemistry), with minor modifications. First, after emulsion generation, nuclei were incubated on ice for 15 minutes to promote nuclear lysis. Second, the reverse transcription protocol was extended to enable the retention of longer transcripts.

##### Augmentation of Wistar rat reference transcriptome

The rat transcriptome from Ensembl (Rattus norvegicus, Rnor_6.0.96) ^22^ lacks full-length *Ttn* as well as large stretches of other important cardiac-related transcripts including *Ryr2*. Many other transcripts are annotated with extents shorter than the read alignment would suggest, resulting in low read-mapping to the Ensembl transcriptome. We therefore created an augmented reference transcriptome for the rat which was used for this study.

First, bulk RNA sequencing was generated by strand specific, long insert whole transcriptome sequencing as offered by the Genomics Platform of the Broad Institute (genomics.broadinstitute.org). Briefly, poly-adenylated RNA was isolated from the aorta, AV node, and all four cardiac chambers of two male Wistar rats and converted to sequencing-ready Illumina TruSeq libraries according to manufacturer's protocols. Libraries were subjected to paired end 50bp sequencing to a mean depth of ~47,000,000 dually mapping reads per library. A de novo reference transcriptome was created from the bulk RNA-seq data using StringTie unguided ^83^. Only transcripts with at least 5 TPM read evidence were kept.

Our augmented reference transcriptome was created by starting with Ensembl Rnor_6.0.96. Given that we performed nuclear 3’ scRNA-seq, all transcripts were collapsed to the level of a gene body as we expected to find retained introns in our reads. We added annotations for *Ttn* and several other genes from the RGD rat60 reference transcriptome, downloaded from ftp://[ftp.rgd.mcw.edu/pub/data_release/GFF3/Gene/Rat/rat60/](http://ftp.rgd.mcw.edu/pub/data_release/GFF3/Gene/Rat/rat60/) ^84^ and expanded each of the annotations in the Ensembl reference based on two rules: (1) if there is an overlapping gene on the same strand with the same name in RGD, and it does not cause a conflict with another protein-coding Ensembl gene on the same strand, expand the gene definition to match RGD; and (2) if there is an overlapping transcript in the unguided StringTie reference, and it does not cause a conflict with any other Ensembl gene on the same strand, expand the gene definition, in whichever direction(s) possible. Compared to the Ensembl Rnor_6.0.96 transcriptome, typically 5-10% more reads from cardiac samples mapped to this amended transcriptome.

##### Data processing

Most data analysis was performed using the Terra cloud platform (app.terra.bio). BCL files for all datasets were processed using cellranger mkfastq (CellRanger 3.0.2, 10x Genomics) to demultiplex samples and generate FASTQ files. These FASTQ files were trimmed using cutadapt^4^ to remove the template switch oligo adapter sequence and its reverse complement [AAGCAGTGGTATCAACGCAGAGTACATGGG, CCCATGTACTCTGCGTTGATACCACTGCTT] (max_error_rate=0.07, min_overlap=10) and all four homopolymer repeats [A30, C30, G30, T30] (max_error_rate=0.1, min_overlap=20). The trimmed FASTQ files were used as input to cellranger count (CellRanger 3.0.2) in order to obtain count matrices, using our modified rat reference transcriptome for nuclear scRNA-seq, and setting --expect-cells to 5000.

##### Sample-level quality control

Quality control at the level of entire samples was performed by examining QC metrics produced by cellranger count, as well as UMAP plots and plots of log(UMI count) versus log(droplet ID) ranked by decreasing UMI count. 11 samples were identified as such strong outliers that they were deemed to be QC failures and subsequently removed. CellRanger count metrics, which were part of the considerations, in addition to the UMI curves, are provided for all samples in Supplementary Table 1.

##### Removal of ambient RNA and cell calling

Background noise was removed from count matrix data on a per-sample basis using CellBender remove-background (<https://github.com/broadinstitute/CellBender>) ^23^ version 0.2.0, with the following parameters: expected-cells=5000; fpr=0.01; total-droplets-included=20000; epochs=150; z-dim=100. This produced a cleaned count matrix per sample, and additionally removed empty droplets. To visualize the reduction of ambient RNA, see **Supplementary Figure 29**.

##### Quality control for nuclei

The number of reads per nucleus mapping to introns, exons, and junctions was tabulated using scR-Invex (Aaron Graubert, François Aguet; https://github.com/broadinstitute/scrinvex). Quality control at the level of individual nuclei was performed separately for each sample. QC metrics calculated per nucleus included log(fraction of reads from mitochondrial genes), fraction of reads mapping to exons, entropy of gene expression (python ndd package), and doublet score as calculated by scrublet ^85^. Outlier nuclei were detected using a 4-dimensional outlier detection algorithm using the above four QC metrics, fitted on those nuclei with fraction of reads from mitochondrial genes < 85th percentile, fraction of reads from exons < 90th percentile, doublet score < 30th percentile, and entropy of gene expression > 5 and < 9. Outlier detection was performed using the scikit-learn function LocalOutlierFactor (contamination=0.02, n_neighbors=10, novelty=True). The following additional hard cutoffs were applied after outliers were removed: n_gene > 100, entropy > 4, scrublet_score < 0.3, exon_fraction < 0.4, mito_fraction < 0.1 except for aorta samples where mito_fraction < 0.2. Experiments typically retained between 4000 and 10000 cells after this quality control procedure.

##### Creation of map and clustering

Count matrices for passing nuclei from each sample were aggregated into one large count matrix in scanpy 1.8.2. ^24^ Highly variable genes were computed using Seurat 3 (method=vst, n_genes=2000). Batch effect correction was performed using scVI 0.6.5 (latent_dimension=50, max_epochs=150, early_stopping=True, only using highly variable genes) ^25^ with the batch variable being individual rat (or tissue-pool). Counts were not altered using scVI. Batch-corrected latent embeddings of each nucleus from scVI were used to create a two-dimensional map using the uniform manifold approximation and projection for dimension reduction (UMAP) algorithm ^26^. Nuclei in the aggregated map were clustered using the Leiden algorithm, computing nearest-neighbor distances using Euclidean distance in the space of the scVI latent representation. Leiden clustering was run at various resolutions.

##### Differential expression testing

Differential expression tests were conducted using R limma ^27^. Testing was carried out as per the recommendation by Lun and Marioni ^28^, after (1) summing count data over appropriate groupings (sample or individual rat or sample-by-cluster, etc. depending on the test), (2) normalizing using DESeq2, and (3) correcting for the mean-variance trend using Voom ^30^. Only genes with summed, DESeq2-normalized counts with a mean of >= 2 were tested. Multiple-testing correction was performed using the Benjamini-Hochberg method.

Background RNA was largely removed during pre-processing by CellBender. However, noise removal is necessarily imperfect, and even residual counts from extremely highly-expressed genes (e.g. *Nppa*) can show up as “significantly differentially expressed” due to ambient/background RNA. These cases are flagged using the following heuristic to calculate a “background probability” for each differential expression result. A per-gene background probability is computed by summing counts over all nuclei in the experiment and taking the value of the empirical cumulative distribution for each gene. A per-test background probability is computed by examining the expression of each gene within the groups being tested versus outside the tested groups. For example, in a test between atrial and ventricular FBs, the CMs (containing *Nppa*) would be out-of-group. We compute the probability that each gene is contamination from out-of-group cells as ${p^{test}}_{g}=(1 -\frac{PP{V_{g}}^{0} + PP{V_{g}}^{1}}{2}) p_{g}$, where $PP{V_{g}}^{0}=\frac{{f_{g,0}}^{T}}{{f_{g,0}}^{T}+{f_{g,0}}^{N}+ 1e-5}$ is the positive predictive value for expression > 0 in the target cell type. Here ${f_{g,0}}^{T}$ is the fraction of cells in the tested cell group expressing > 0 counts of gene $g$, and ${f_{g,0}}^{N}$ is the same for nuclei that were not included in the DE test. $PP{V_{g}}^{1}$ is the analogous quantity when counting cells based on expression > 1 count (a noise mitigation strategy). Intuitively, $(1 -\frac{PP{V_{g}}^{0} + PP{V_{g}}^{1}}{2})$ represents the probability that the gene belongs to cells that were *not* included in the DE test either as case or control. $p_{g}$ is an overall noise probability per gene, computed by converting the empirical counts per gene observed in the entire dataset to an empirical cumulative distribution function. The most highly-expressed gene has a $p_{g} \approx1$, while lowly-expressed genes have values near zero, reflecting the fact that lowly-expressed genes are hardly ever going to contribute to noise under any circumstances. ${p^{test}}_{g}$ then represents a likelihood that a particular gene resulted in spurious “significant” differential expression due to contamination from nuclei outside the tested group. In the running example, due to the high expression of *Nppa*, in particular in atrial CMs, it would be assigned a high background probability ${p^{test}}_{g}$ in a test of atrial versus ventricular FBs.

##### Marker gene discovery

Differential expression testing was performed for each gene by comparing expression in a given cluster to all other clusters using the differential expression testing framework above, and summing count data per sample per cluster. Sample-cluster combinations with fewer than 25 cells were excluded from analysis. Contrasts of one cell cluster versus all others were fit in limma using the model (~ 0 + cluster + tissue), along with duplicateCorrelation per individual rat (or tissue-pool), to extract an estimate of a log fold-change between the given cluster and all others. Tens to hundreds of genes were found to be significantly differentially-expressed in each cluster (false discovery rate 0.001). Cell types were named by examination of the top up-regulated genes in a cluster and manual searching of the literature (in combination with highest-expressed genes and gene set enrichment analyses). In **Supplementary Figure 2**, marker genes are ranked by sorting based on log2-fold-change times mean-counts-per-cell in the cluster of interest. This ensures the prioritization of genes with both high counts as well as large over-representation in the cluster of interest.

##### Pathway analyses

Analyses of pathways / gene-sets were carried out using gene set enrichment analysis, as implemented in the fgsea package in R ^31^. Human gene sets were downloaded from MSigDB v7.1: c2.cp.biocarta, c2.cp.kegg, c2.cp.pid, and c5.all (for GO analyses). Rat genes were mapped to human genes using biomaRt ^86^, manually adding genes from our transcriptome that lacked Ensembl IDs, and removing genes with multiple mappings. The final mapping is included as **Supplementary Table 8**. GSEA was performed using the t-statistic from a relevant differential expression test to rank-order genes. Genes with probability > 0.4 that a differential expression result was caused by ambient RNA contamination were removed before running GSEA. A million permutations were used to calculate a p-value using fgsea.

##### Cell-cell communication analyses

Receptor-ligand interactions were inferred based on counts of receptor genes and ligand genes in all pairs of cell types, using the CellPhoneDB database ^5^ and the squidpy 1.0.0 software package’s “ligrec” permutation test ^32^, which computes a mean interaction strength and a p-value that a particular receptor-ligand pair is enriched in a certain cell-type-pair as compared to all other cell-type-pairs, similar to CellPhoneDB’s own software. The permutation test was run separately on each sample, meaning that only nuclei within the same physical vicinity in a biological sample were ever tested for cell-cell communication. Results from all samples were aggregated by taking the mean of the mean interaction strength over all samples, and a new p-value that a result was significant in at least one sample was computed as the product of p-values. For tissue comparisons, samples were aggregated by tissue of origin. Differences in mean interaction strengths between tissues were assessed for statistical significance using Wilcoxon rank-sum tests.

### Supplementary Figures

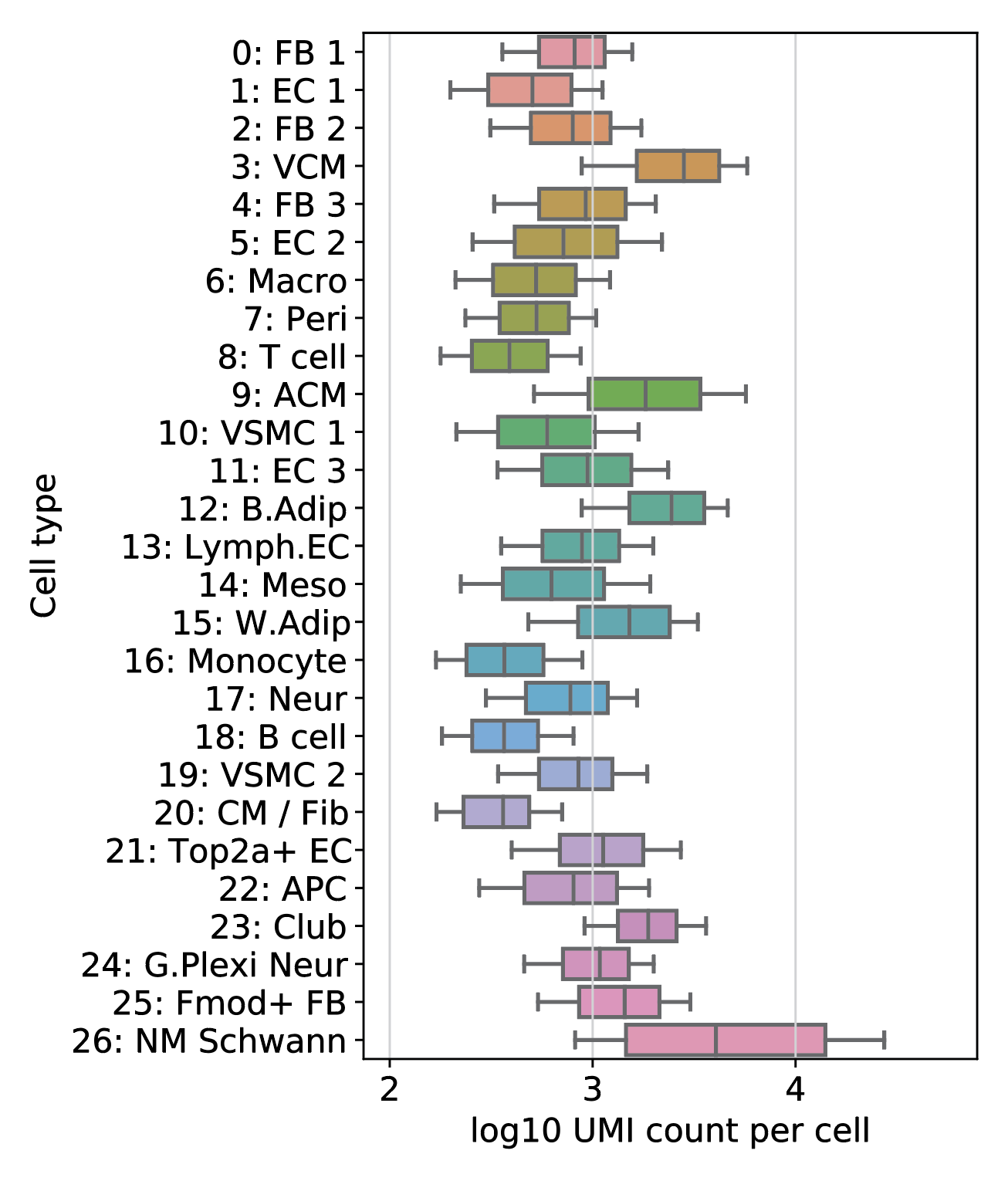

**Supplementary Figure 1. Complexity of each cell type.** Boxplot shows the distribution of UMI counts per nucleus, broken down by cell type. Boxes show the quartiles of the distribution of UMI counts for each cell type on a log scale, while whiskers extend from the 10th to the 90th percentile of the distributions. NM Schwann cells, cardiomyocytes, and adipocytes are among the cell types producing the most mRNA.

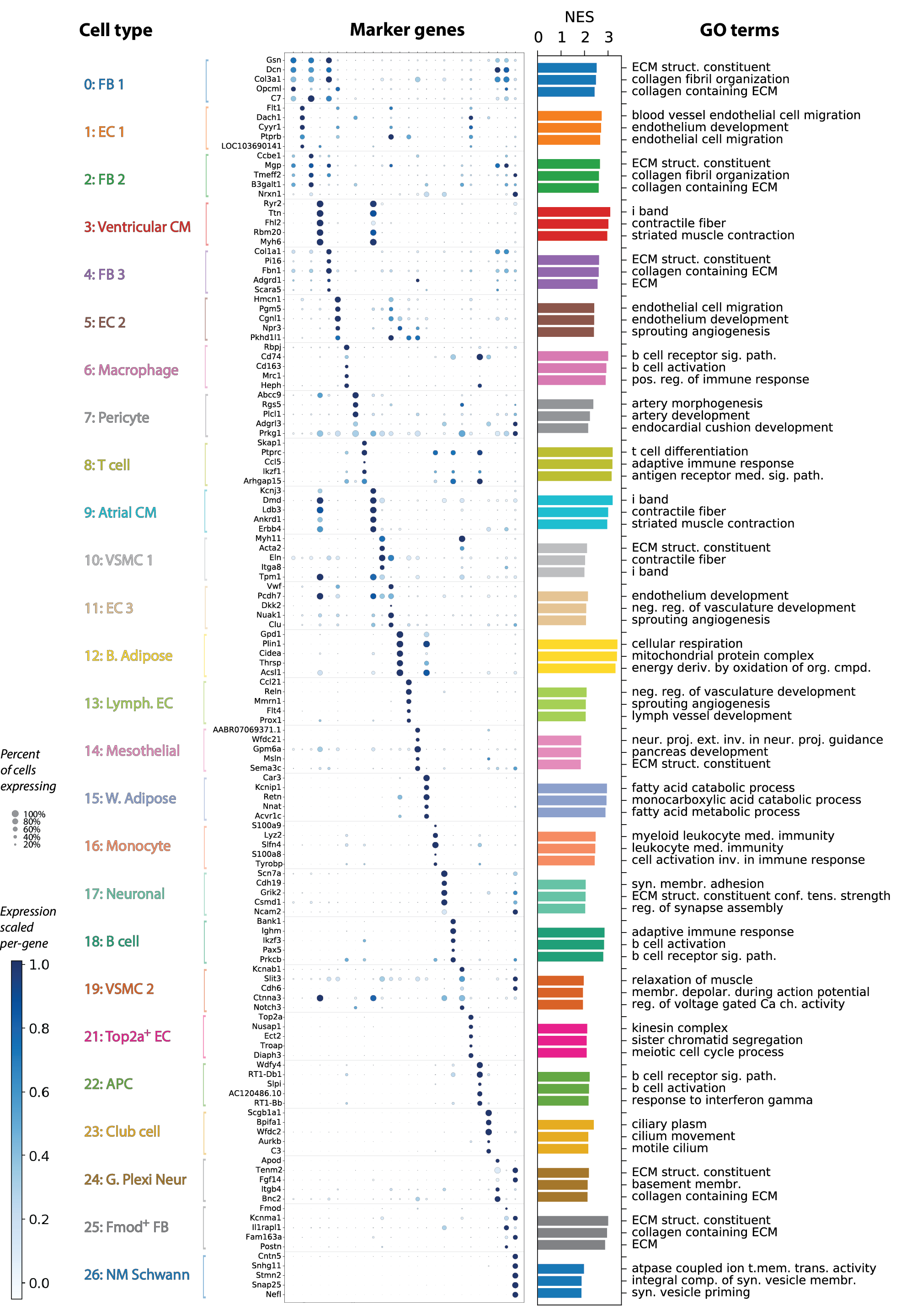

**Supplementary Figure 2. Cluster annotation in detail, by marker genes and GO-term enrichment.** Top five marker genes are displayed for each cluster, where marker genes are prioritized based on the results of a differential expression test. The dotplot uses dot size to convey the percent of cells expressing a gene, and color is proportional to the expression level, here scaled to a maximum of 1.0 for each gene. To the right of the dotplot, a bar chart displays the normalized enrichment score (NES) from a gene set enrichment analysis using GO terms (c5.all.v7.1 gene sets from MSigDB). The top three GO terms (by NES) for each cell type are annotated. All the displayed GO terms are significantly enriched in a single-cluster-versus-all comparison at a false discovery rate of 0.05.

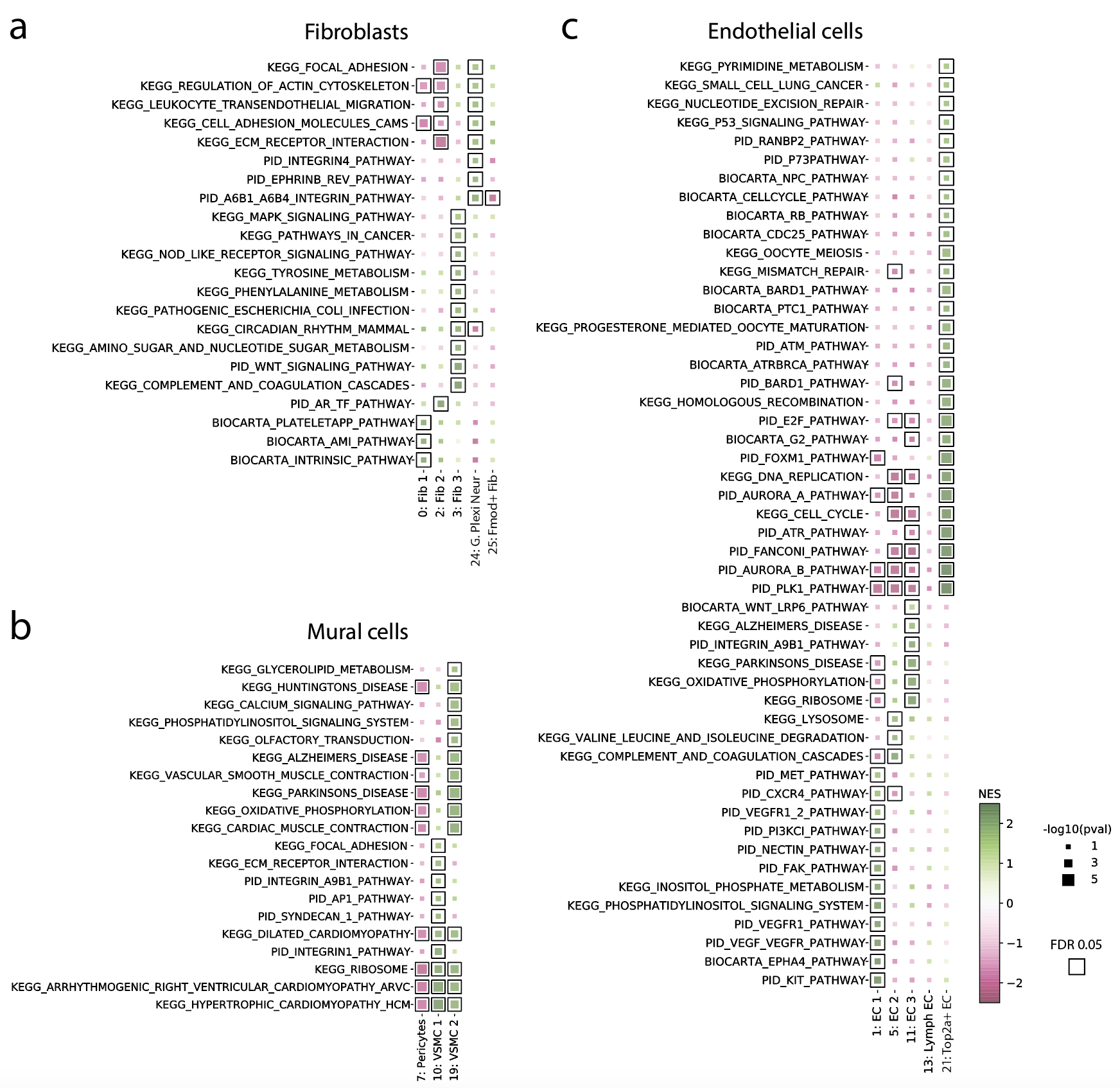

**Supplementary Figure 3. Differentially enriched pathways among related cell types.** Pathway enrichment within three major “meta-clusters”: (a) fibroblasts, plus cluster 24 which is nearby on the global UMAP, (b) mural cells, and (c) endothelial cells. These pathways highlight differences between cell types in the same meta-cluster. Differential expression tests were carried out for each cell type versus all others in the same meta-cluster. Using these differential expression results (t-statistics), gene set enrichment analysis was performed for Biocarta, Kegg, and PID pathways (c2.cp.biocarta, c2.cp.kegg, and c2.cp.pid from MSigDB, v7.1). All pathways where at least one cell type reached FDR 0.05 significance are included, and significance is denoted by the black boxes. The color of the squares corresponds to the normalized enrichment score versus other cell types in the same meta-cluster, while the size corresponds to the Benjamini-Hochberg adjusted p-value.

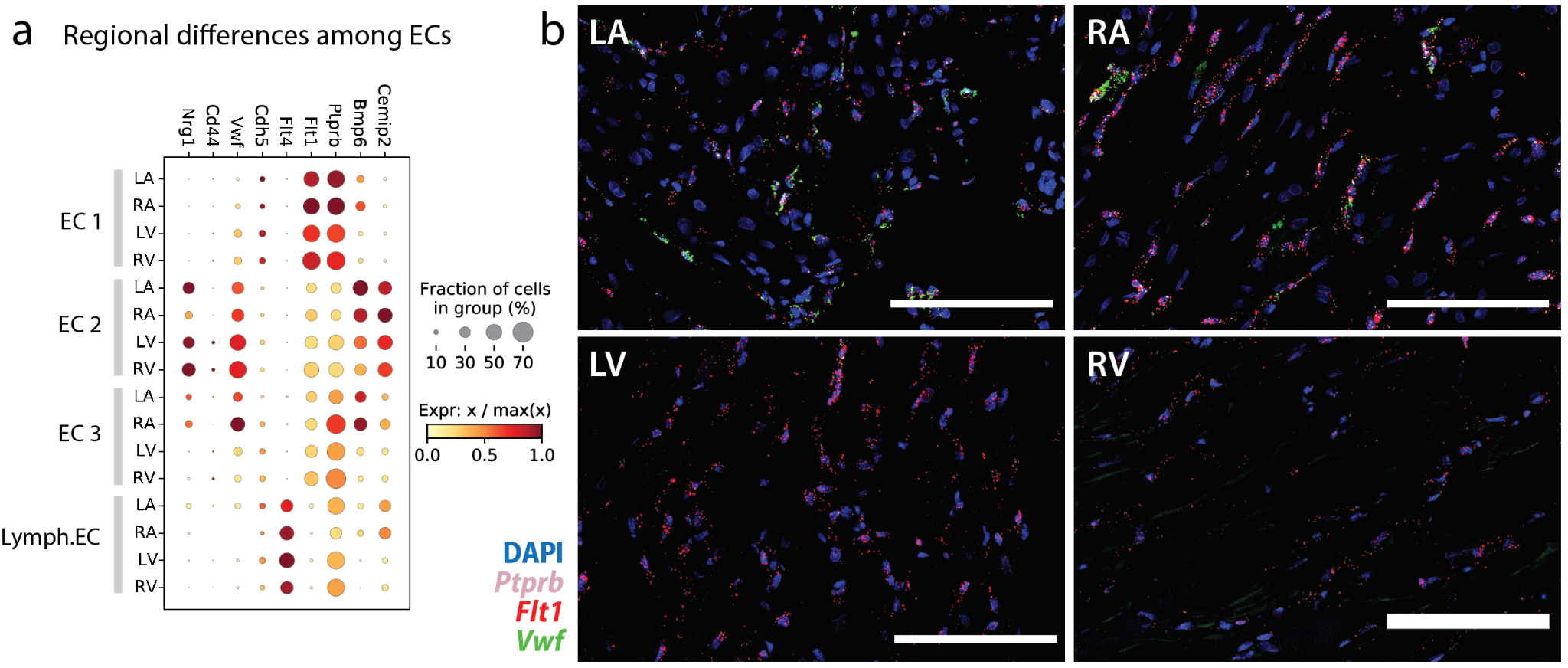

**Supplementary Figure 4. Different EC composition in atria versus ventricles.** (See Figure 2d-e for an overview of the cell-type composition of each tissue.) **(a)** Marker genes which capture some of the variability among EC clusters. *Flt1* and *Ptprb* are present in EC1, EC2, and EC3. Expression values (dot colors) are normalized so that, for each gene, 0 denotes zero expression and 1 denotes the maximum expression in any of the conditions along the y-axis. **(b)** RNAscope imaging of the distribution of different EC types in the 4 chambers of the heart (RA, LA, LV, RV). Tissue was counterstained with DAPI. All scale bars are 100 microns.

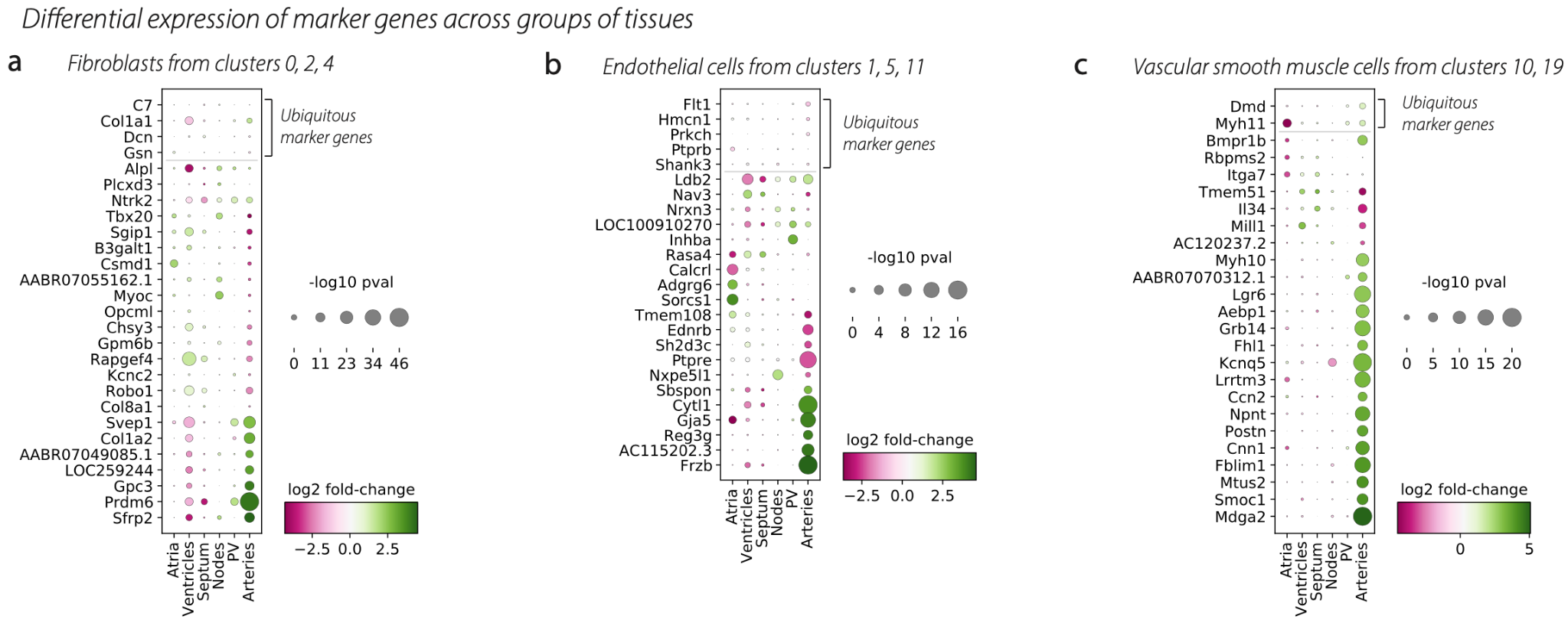

**Supplementary Figure 5. Cell type marker genes differ by tissue of origin.** Summary statistics from differential expression tests for one tissue versus all others. Genes are restricted to those that are marker genes for the given cell type in at least 1 tissue. Dotplots show fold changes and p-values for tests among **(a)** FBs, **(b)** ECs, and **(c)** VSMCs.

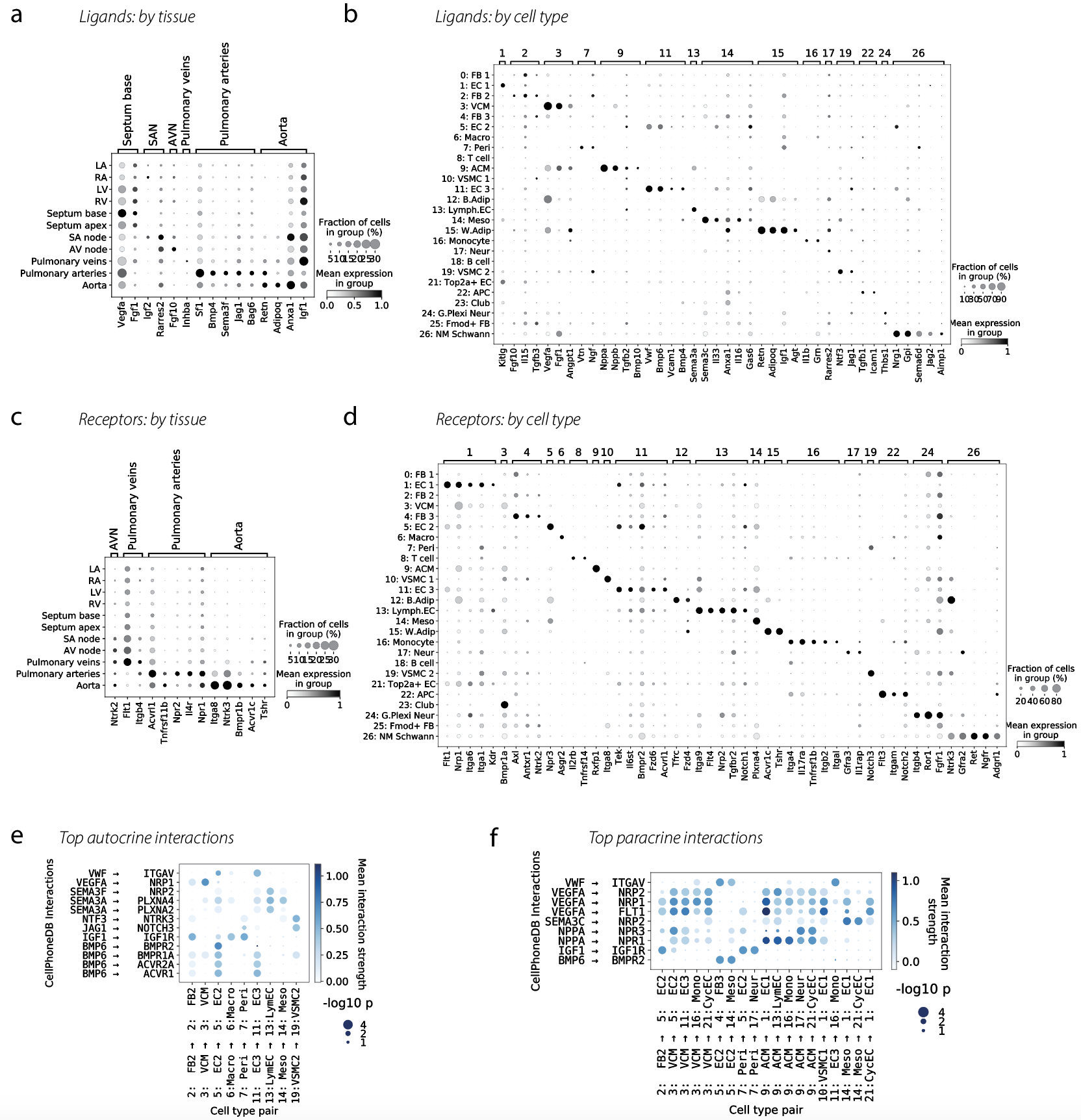

**Supplementary Figure 6. Ligands and receptors and top cell-cell communication pathways.** Dotplots show selected **(a-b)** ligands and **(c-d)** receptors that are differentially expressed either by tissue or by cell type. Dotplots are normalized per-gene so that the maximum expression across groups is 1 and zero expression is 0. Receptors and ligands are taken from the CellPhoneDB database of interactions. Brackets at the top of each dotplot serve to group genes by the condition in which the differential expression effect size was the largest, but the brackets do not indicate exclusivity of expression. **(e)** Top paracrine interactions across the entire dataset. **(f)** Top autocrine interactions across the entire dataset. Interaction strengths are computed using CellPhoneDB.

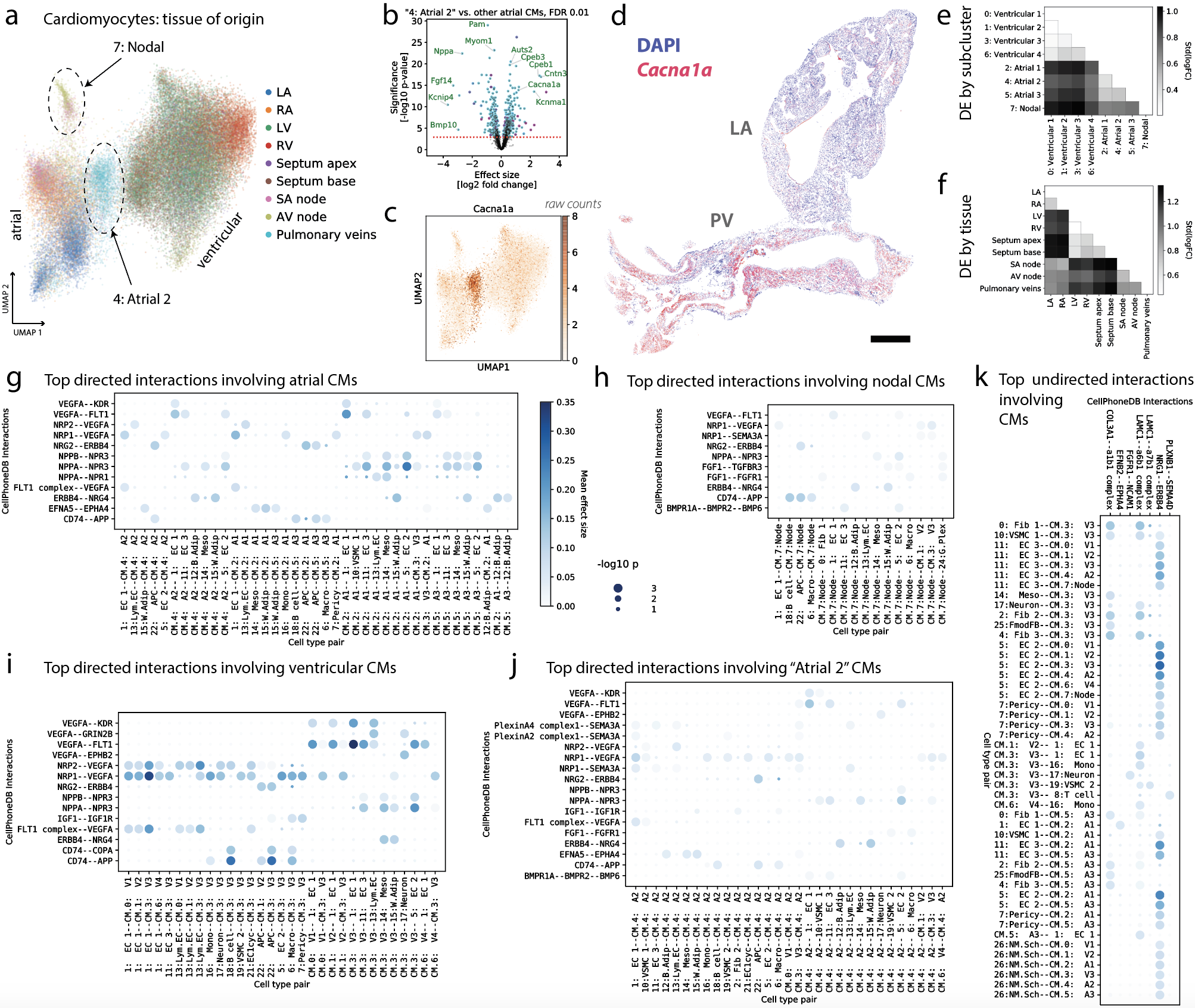

**Supplementary Figure 7. Further detail about CM subclustering. (a)** UMAP showing the *de novo* CM subclustering (see **Figure 5b**), here coloring each cell by tissue of origin. The labels “4: Atrial 2” and “7: Nodal” reflect subcluster names. **(b)** Volcano plot showing top differentially expressed genes between the PV-enriched subcluster “4: Atrial 2” and all other CMs from LA and RA. *Cacna1a* is upregulated in the PV-specific CM subcluster. **(c)** UMAP of CMs colored by counts of *Cacna1a*, showing higher expression in the PV-specific “4: Atrial 2” subcluster. **(d)** RNAscope image of *Cacna1a* in a section of PV and LA tissue, confirming the snRNA-seq results in panels **b-c**. Tissue is counterstained with DAPI (nuclei). **(e)** Heatmap shows aggregate summary statistics about differential expression between tissues. Each entry in the heatmap corresponds to a separate pairwise differential expression test between all the CMs in two tissues. The color value is the standard deviation of the log fold changes for all tested genes, and is a proxy for how dissimilar two tissues are. Larger values mean that CM transcription in the two tissues is more different. **(f)** Similar heatmap as in **e**, showing all pairwise comparisons between CM subclusters. **(g-k)** Dotplots showing the top significant cell-cell communication interactions involving the CM subclusters. Colorbar and dot size legend applies to all panels. “Undirected” interactions (panel **k**) in the CellPhoneDB database are those which do not follow the pattern that one interacting partner is designated “ligand” and the other “receptor”. Each interaction shown is FDR significant at 0.01 in at least one x-axis condition.

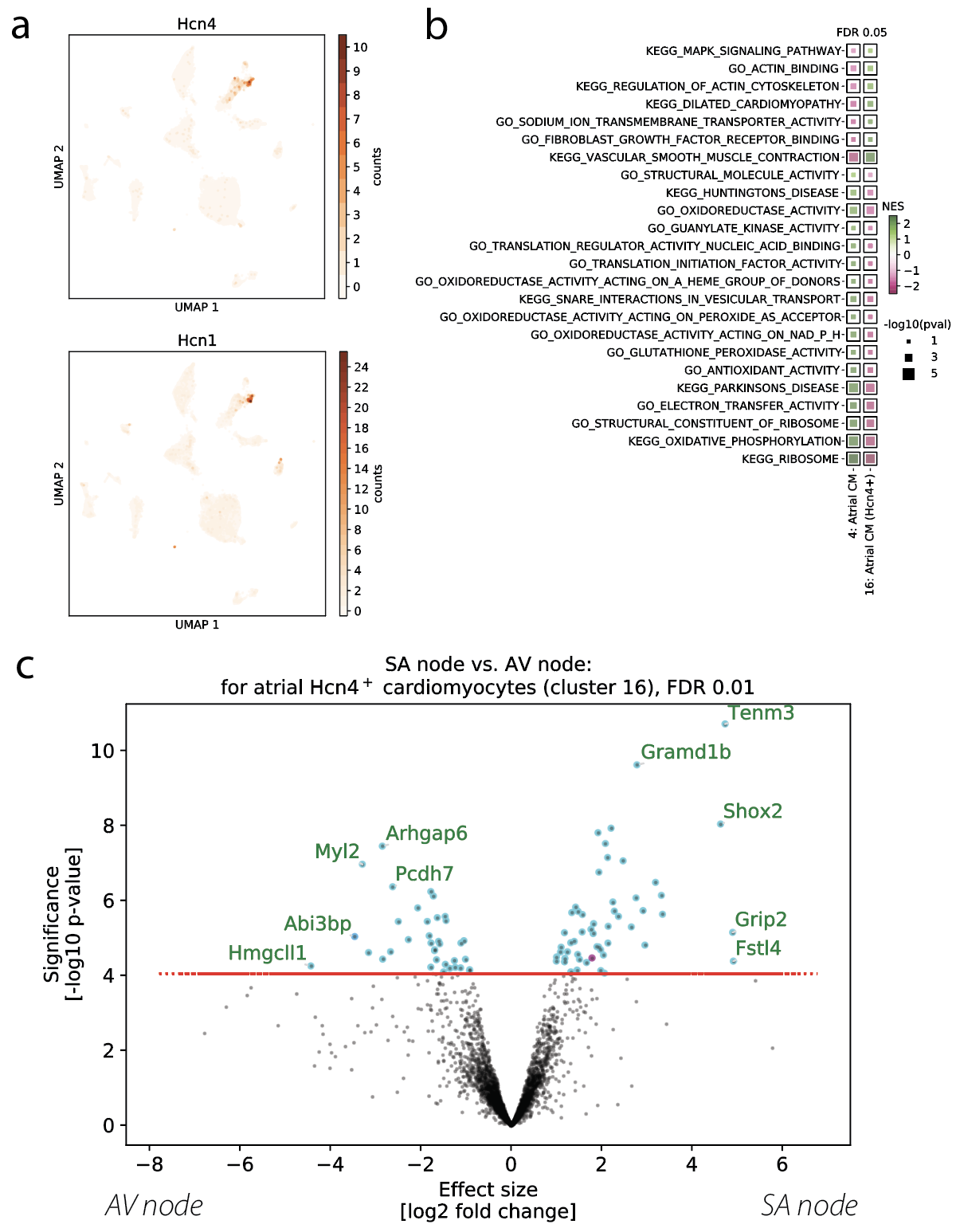

**Supplementary Figure 8. Additional detail about the map of the SAN and AVN.** (Companion to Figure 6.) **(a)** UMAP plots show the expression of *Hcn1* and *Hcn4*, indicating its enrichment in cluster 16, the pacemaker atrial CM cluster. **(b)** All KEGG, Biocarta, PID, and GO molecular function pathways that are identified as having significantly different enrichment in pacemaker CMs as compared to atrial CMs (MSigDB pathways v7.1). **(c)** Volcano plot shows results of a differential expression test between pacemaker CMs in the SAN versus the AVN.

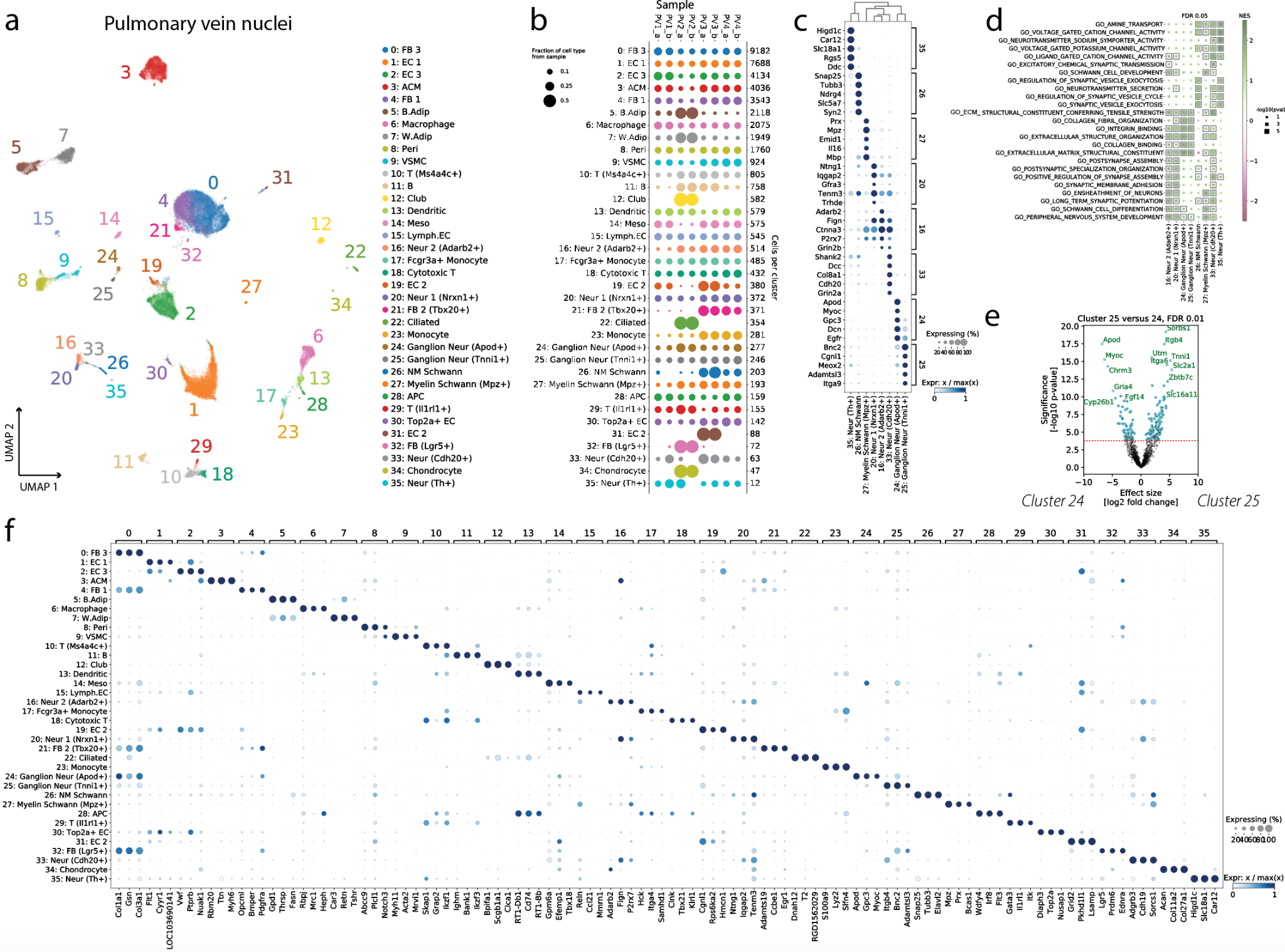

**Supplementary Figure 9. Cells of the pulmonary veins. (a)** UMAP of 46,099 nuclei from the pulmonary veins. **(b)** Breakdown of the representation of each cluster in each sample. Both technical replicates of sample PV2 contain all cells from clusters 12, 22, 32, and 34 (presumably of pulmonary origin, representing a small amount of contamination). **(c)** Neuronal cell types in the PV samples. Dendrogram at top shows relatedness between clusters, and the dotplot shows the top five marker genes for each cluster. **(d)** Top GSEA results for GO biological process and molecular function pathways, run using results of differential expression tests of each neuronal cell type versus all other clusters. Boxes indicate a result is significant at FDR 0.05. **(e)** Volcano plot highlights differences between the closely-related clusters 24 and 25. **(f)** Dotplot shows the top three marker genes for each PV cluster.

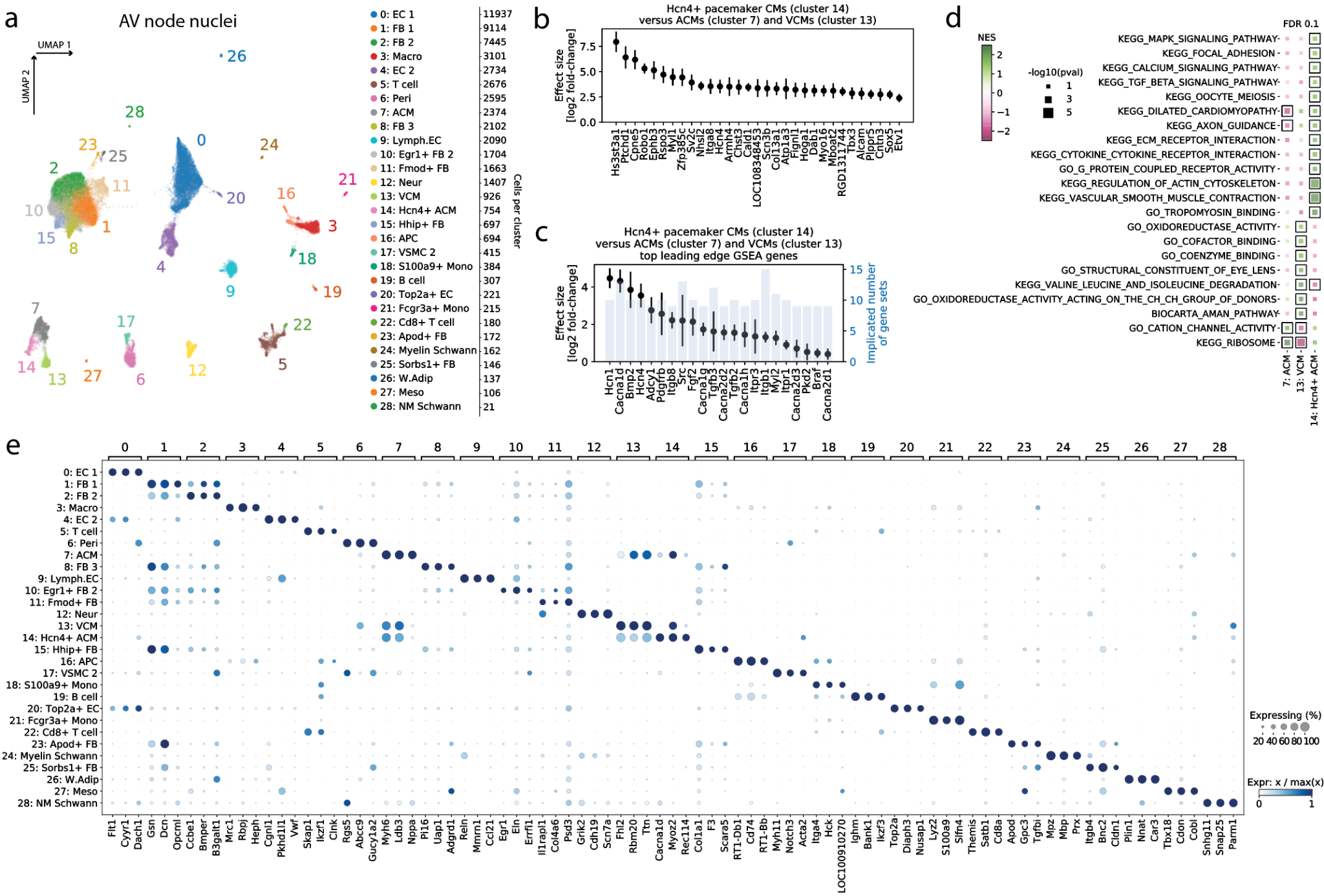

**Supplementary Figure 10. Cells of the AV node. (a)** 56,479 nuclei from 10 AV node samples comprising 3 pooled-tissue library preps. Numbers along the right-hand side denote cells per cluster. **(b)** Upregulation of genes in pacemaker CMs as compared to atrial CMs. **(c)** Same as **b**, but prioritizing a short list of genes based on the number of gene sets (significant at FDR 0.3) in which the gene was part of the leading edge in the gene set enrichment analysis (GSEA) analysis shown in panel **d**. Light blue bars show the number of upregulated gene sets in which the gene is implicated. **(d)** GSEA results significant at FDR 0.1 after Benjamini-Hochberg multiple testing correction, comparing in each case one cluster versus the other two. **(e)** Dotplot of the top 3 marker genes for each cluster.

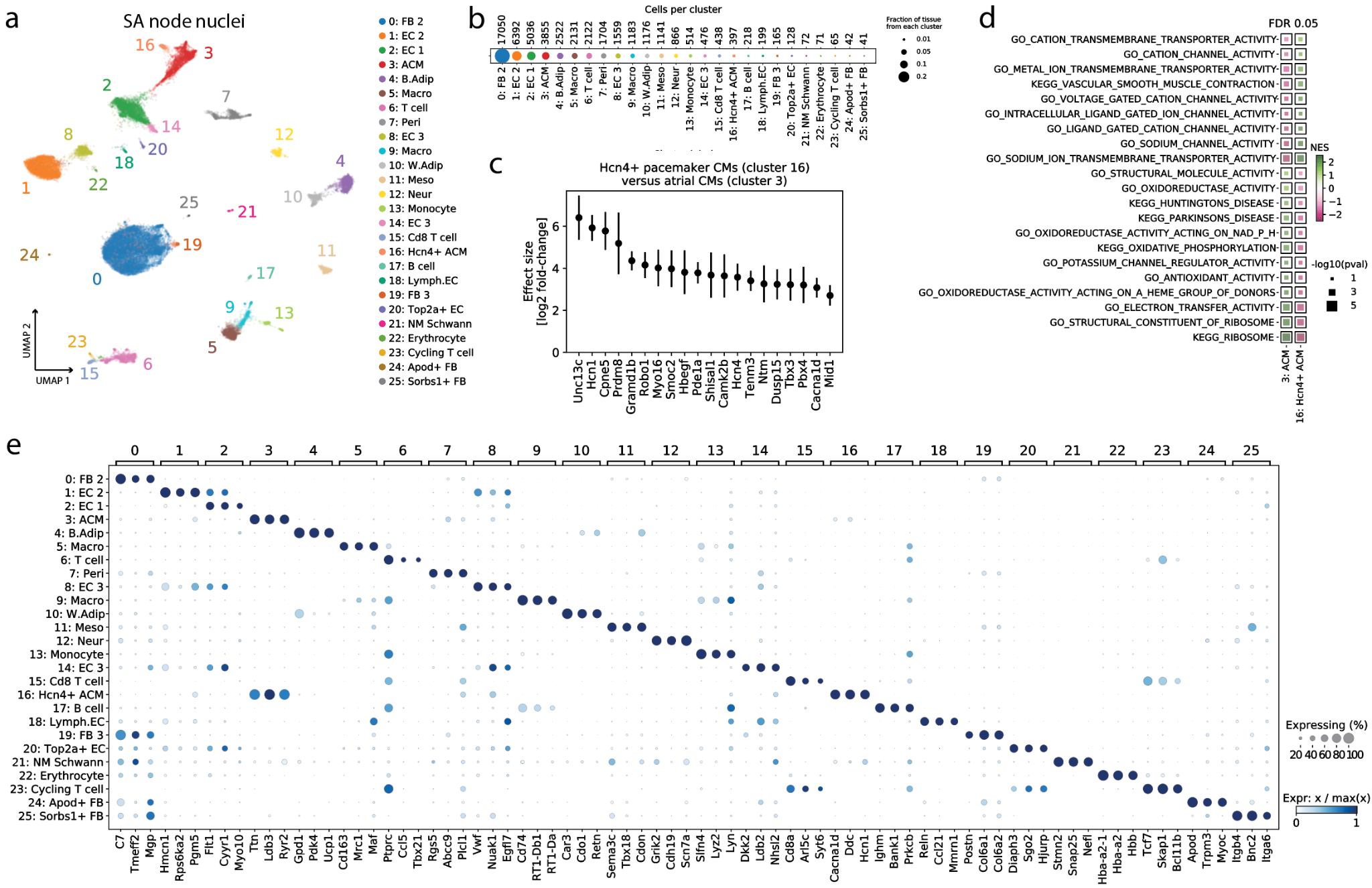

**Supplementary Figure 11. Cells of the SA node. (a)** 49,563 nuclei from 9 SA node samples comprising 3 pooled-tissue library preps. **(b)** Composition of the SA node by cell type. **(c)** Upregulation of genes in pacemaker CMs as compared to atrial CMs. **(d)** Gene set enrichment analysis results significant at FDR 0.05 after Benjamini-Hochberg multiple testing correction. **(e)** Dotplot of the top 3 marker genes for each cluster.

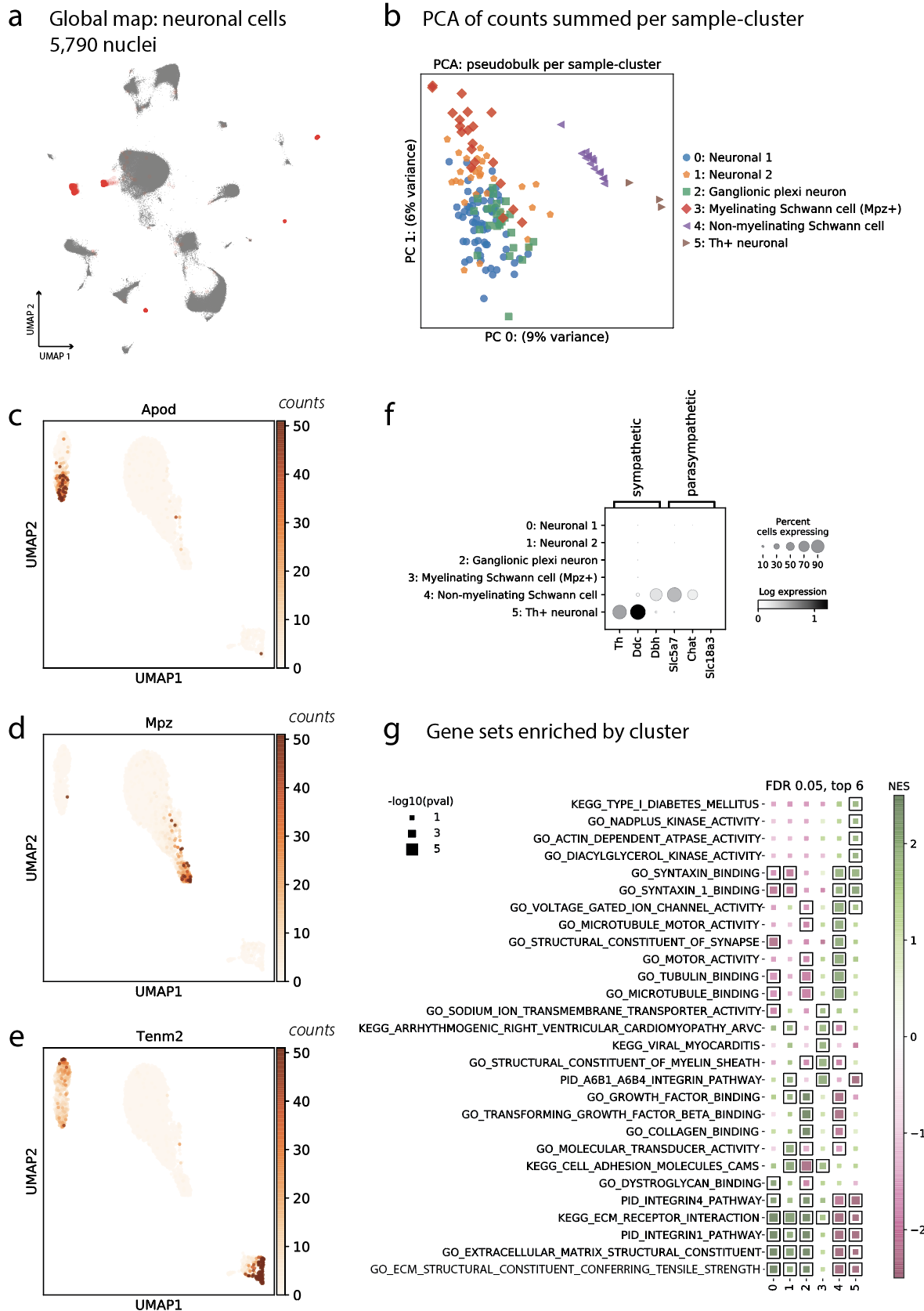

**Supplementary Figure 12. Subclustering of neuronal cell populations. (a)** UMAP of all cells in the experiment, with 5790 neuronal cells highlighted in red. **(b)** Top two principal components of variation among neuronal cells, showing that cells segregate by subcluster. Subclusters are further defined in **Figure 6e,f,h**. **(c-e)** UMAP (same coordinate system as **Figure 6e**, where subclusters are labeled) showing counts of the genes *Apod*, *Mpz*, and *Tenm2*, which have striking and very different distributions across subclusters. **(f)** Dotplot shows the expression of a few canonical markers of the sympathetic and parasympathetic nervous system. **(g)** GSEA results for pathways enriched in one neuronal subcluster versus all other neuronal subclusters. Pathways significant at FDR 0.05 are highlighted with a box.

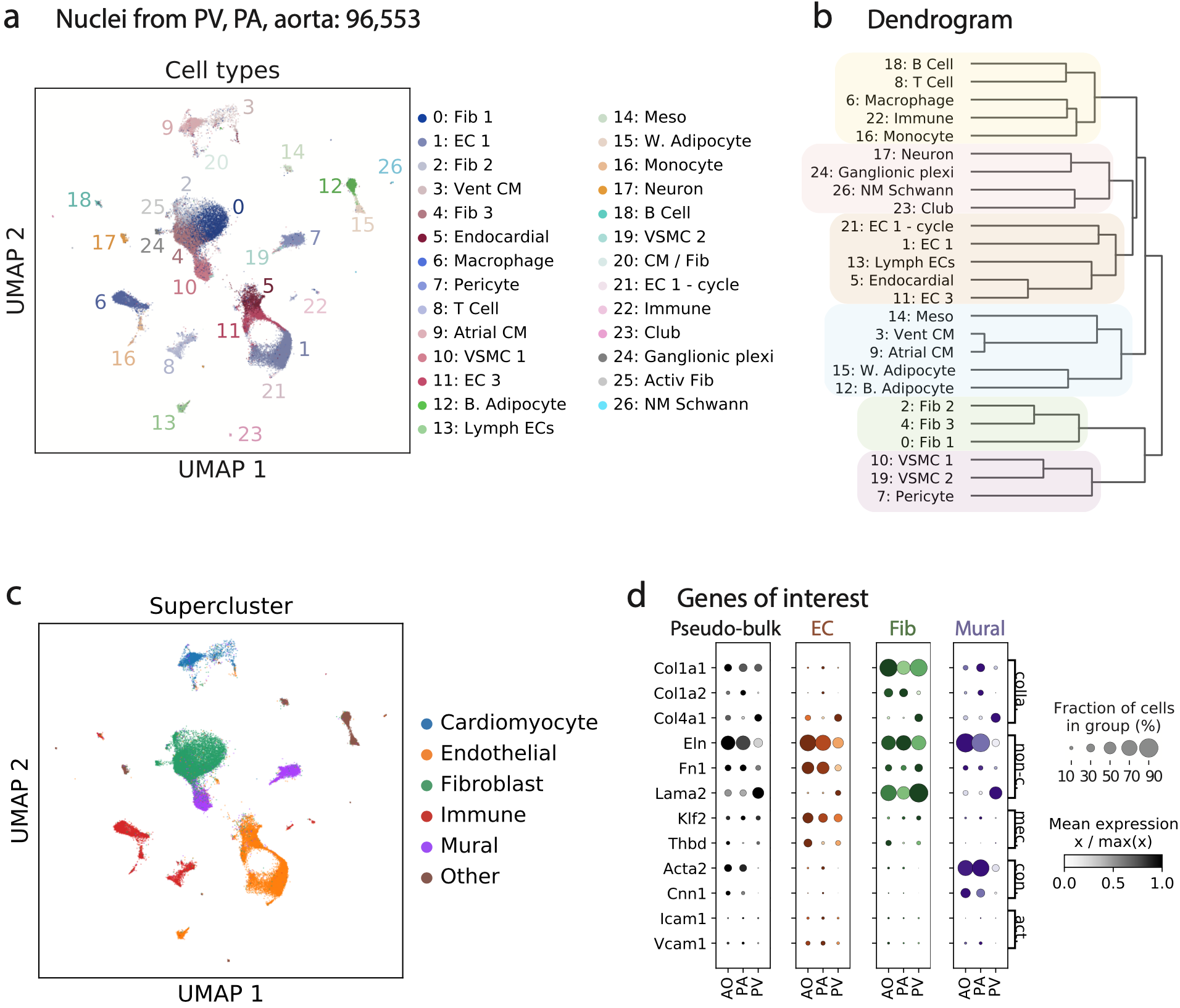

**Supplementary Figure 13. Overview of the cells of the vasculature. (a)** UMAP of 96,533 cells from the PV, PA, and Ao, using the same UMAP coordinate system as the global map in **Figure 2a**. **(b)** Dendrogram showing transcriptional similarity between the various clusters. Cell types cluster neatly into related clades, denoted by the colors. **(c)** Same UMAP as in panel **a**, but labeled by a low-resolution cell type category, or “supercluster”. **(d)** Dotplot showing lookups of several genes of interest both at the pseudo-bulk level and for cell type superclusters including ECs, FBs, and mural cells. Tissue specificity is apparent in some cases.

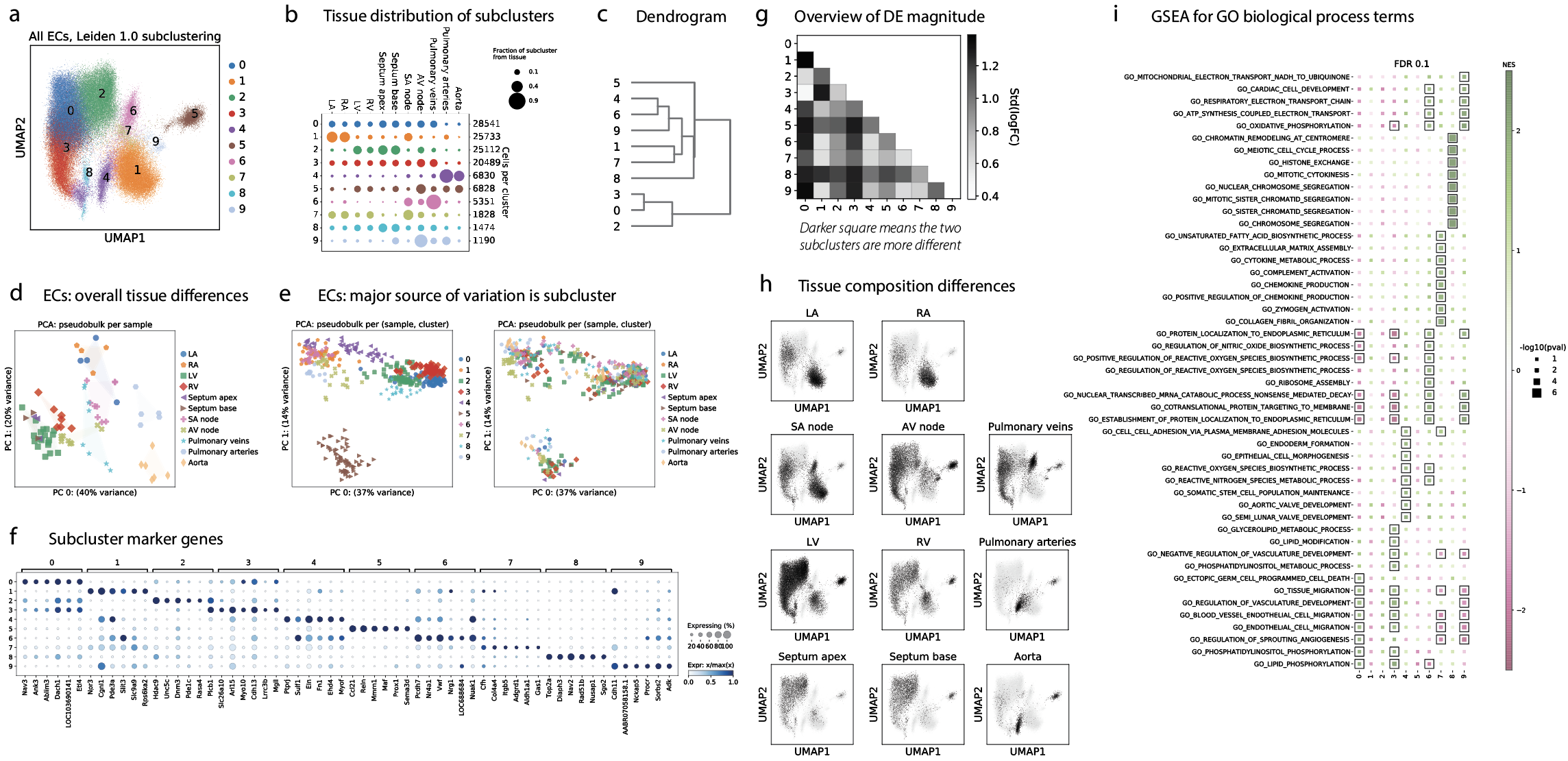

**Supplementary Figure 14. Subclustering of all ECs. (a)** UMAP of 123,416 ECs from all tissues, subclustered at Leiden resolution 1.0. **(b)** Tissue distribution of each EC subcluster. Dot sizes are first normalized so that all cells sum to one each tissue, and then normalized by row, so that each row sums to one. **(c)** Dendrogram showing transcriptional similarly between subclusters. **(d)** PCA plot showing pseudo-bulk EC expression. Each dot is the summed EC expression from one sample. **(e)** Similar PCA plot, this time summing expression per sample per EC subcluster. Each dot is the summed expression from one EC subcluster in one sample. Coloring by EC subcluster (left panel) as compared to tissue (right panel) reveals that the principal components of variation seem to be explained by subcluster rather than tissue of origin. **(f)** Top six marker genes for each EC subcluster. **(g)** Every pairwise differential expression test between EC subclusters, similar to **Supplementary Figure 7b**. Lighter color means the subclusters are more similar. **(h)** Overview of the differences in the presence of each subcluster in each tissue. Each tissue is one panel. UMAPs show all cells in light gray, overlaid in black by cells from the specific tissue. It is visually quite apparent that the tissues use the EC subclusters in very different proportions. **(i)** Top GO biological process terms significant at FDR 0.1 (boxes) in GSEA comparisons of one EC subcluster versus other EC subclusters.

#### S15: imaging of vascular EC subclusters

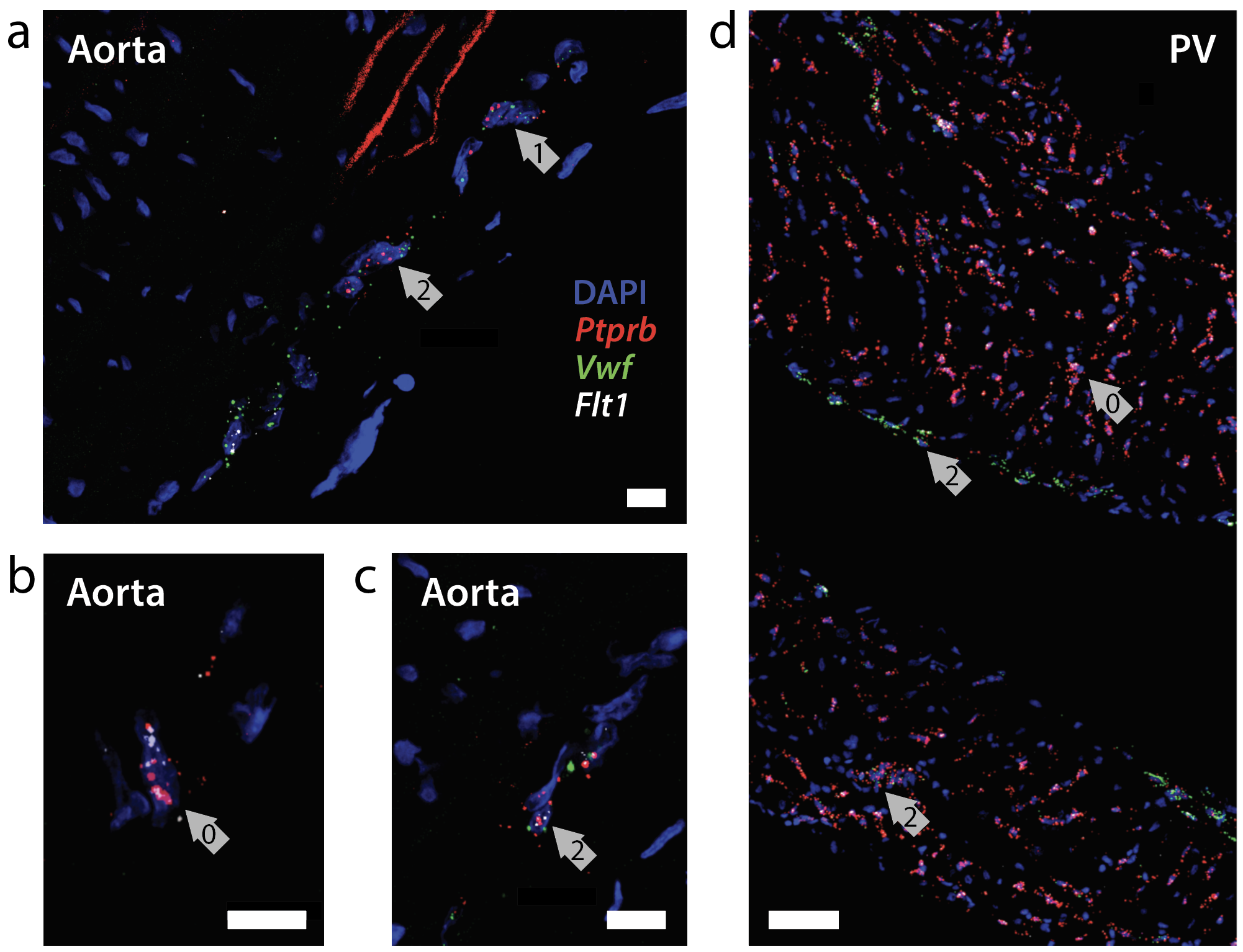

**Supplementary Figure 15. Imaging validation of EC subclusters.** RNAscope imaging validates the presence of EC subclusters 0, 1, and 2 in the Ao, and subclusters 0 and 2 in the PV. **(a)** Cells from EC subclusters 1 and 2 in the Ao. **(b)** Cell from EC subcluster 0 in the Ao. **(c)** Cell from EC subcluster 2 in the Ao. **(d)** Cells from EC subclusters 0 and 2 in the PV.

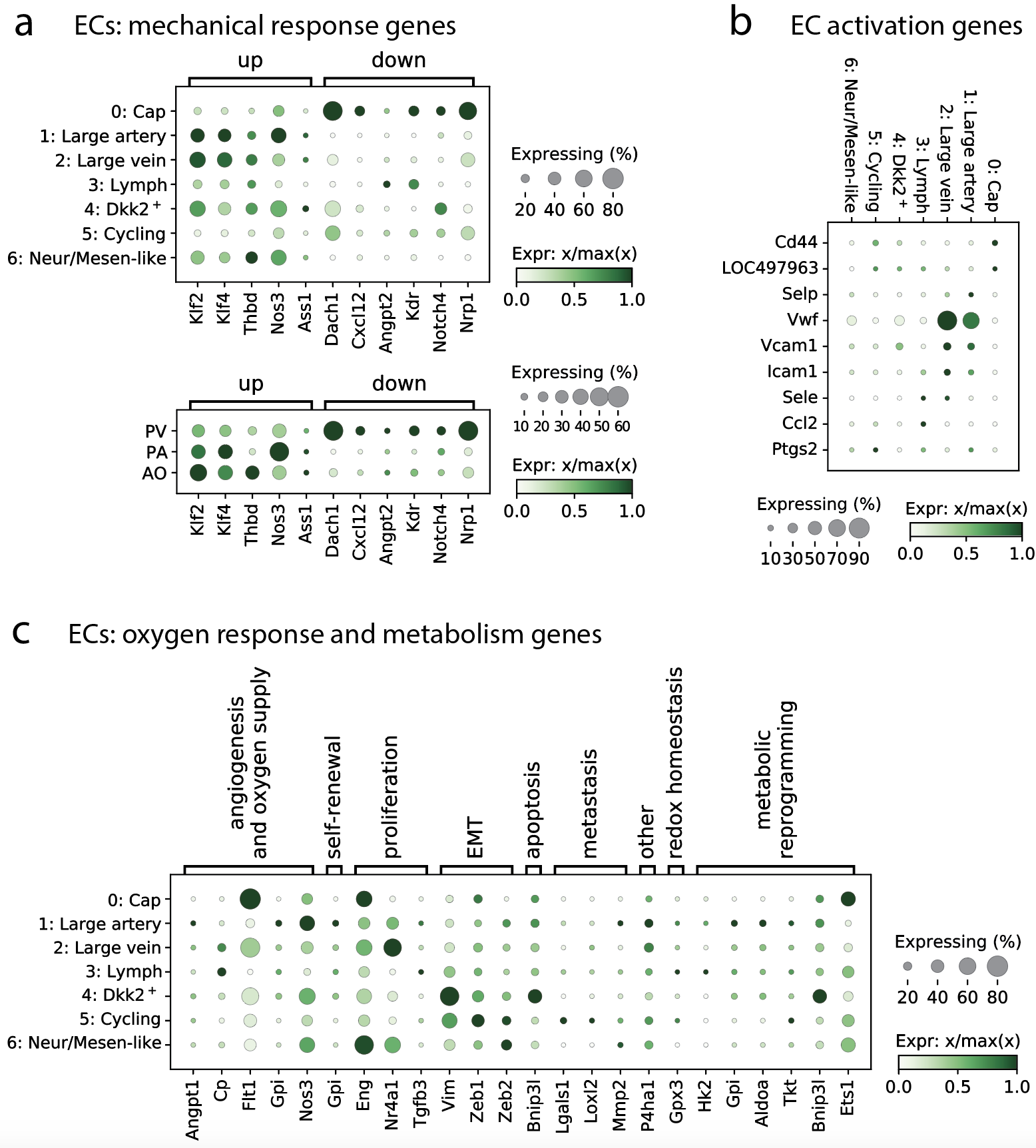

**Supplementary Figure 16. Gene lookups in the vascular EC subclusters. (a)** Mechanical response genes across EC subclusters and tissues (ECs only). **(b)** EC activation genes across EC subclusters. **(c)** Oxygen response and metabolism genes across EC subclusters.

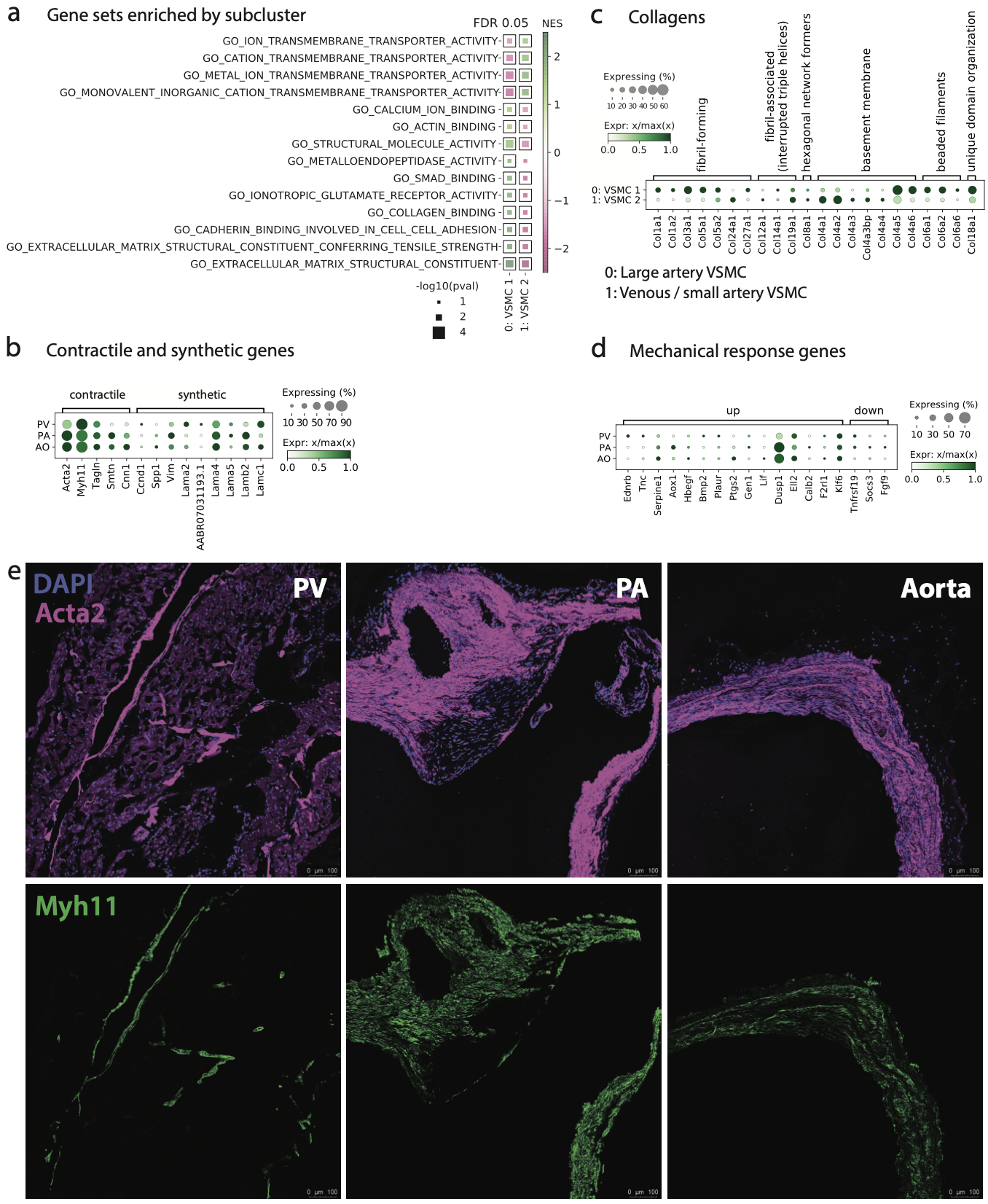

**Supplementary Figure 17. VSMCs from the vasculature. (a)** GO molecular function terms found to be enriched in each VSMC subclustering using GSEA. Black box borders highlight results significant at FDR 0.05 after Benjamini Hochberg multiple testing correction. **(b)** Expression of contractile and synthetic genes across tissues. Plot showing subcluster expression is shown in **Figure** **8d**. **(c)** Expression of collagen genes in each subcluster. **(d)** Mechanical response genes in VSMCs, by tissue. **(e)** Immunofluorescence staining of Myh11 and Acta2 (both of which are present in both VSMC subclusters) in sections from PV, PA, and Ao.

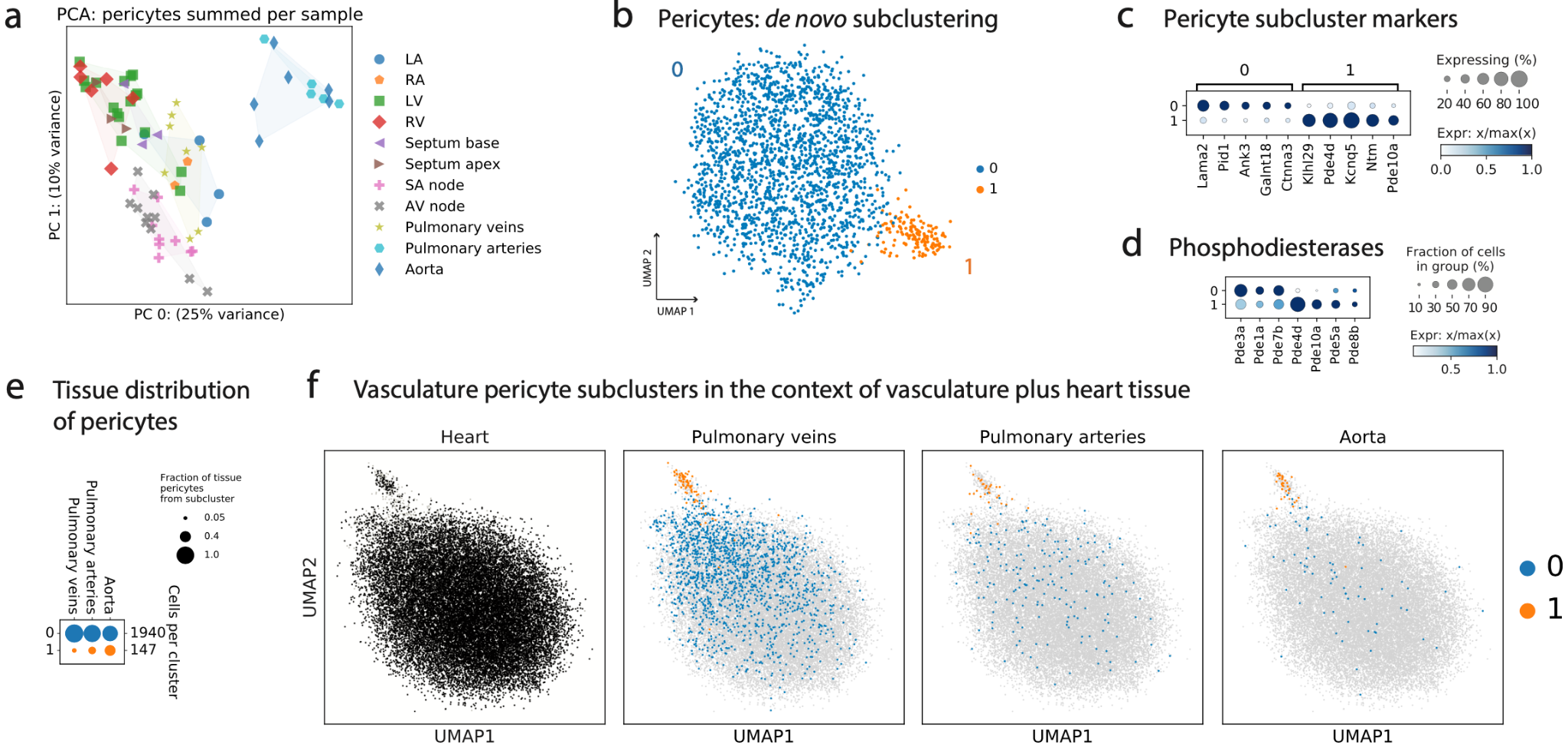

**Supplementary Figure 18. Subclustering pericytes from PV, PA, and Ao. (a)** PCA plot shows how pseudo-bulk pericyte expression differs among samples. Each dot is the summed expression of all pericytes in a single sample. Dots are colored by tissue of origin. **(b)** Subclustering of pericytes from the vasculature. UMAP where each dot is a cell. **(c)** Marker genes for pericyte subclusters. **(d)** Some phosphodiesterases that are expressed in the dataset show specificity for one pericyte subcluster. **(e)** Breakdown of pericyte subcluster composition of PV, PA, and Ao. **(f)** UMAPs show all pericytes from the entire dataset in light gray. Heart panel shows pericytes from the heart overlaid in black. Other panels show pericytes from PV, PA, and Ao respectively, colored by subcluster.

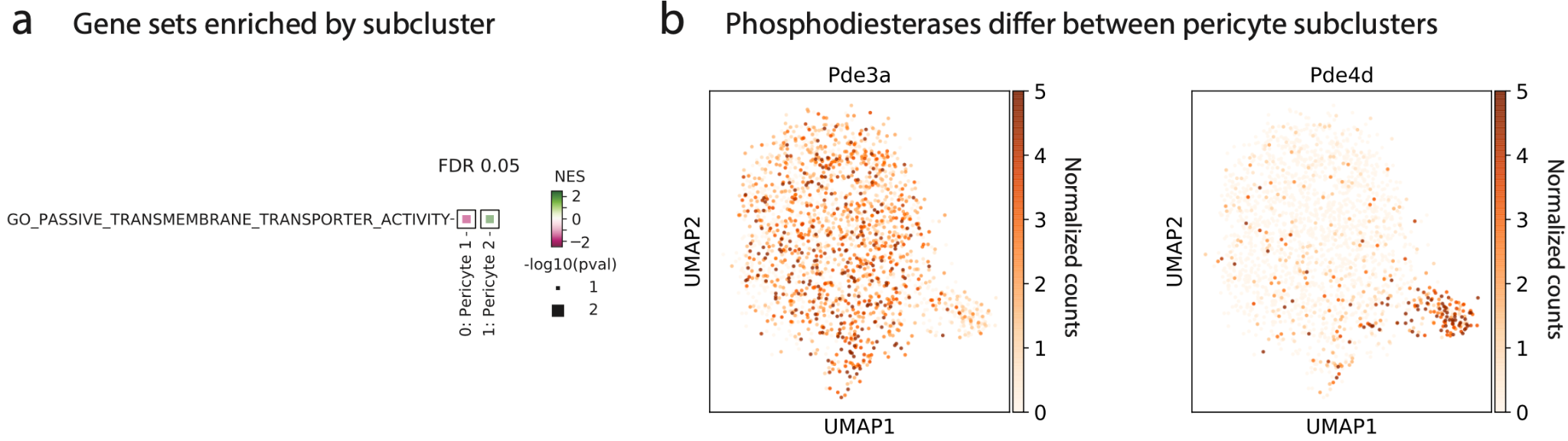

**Supplementary Figure 19. Differences between vascular pericyte subclusters. (a)** Only one GO molecular function gene set meets an FDR 0.05 significance threshold via GSEA. **(b)** UMAPs of two phosphodiesterases whose expression differs strongly between pericyte subclusters (UMAP subcluster labels are shown in **Supplementary Figure 18b**).

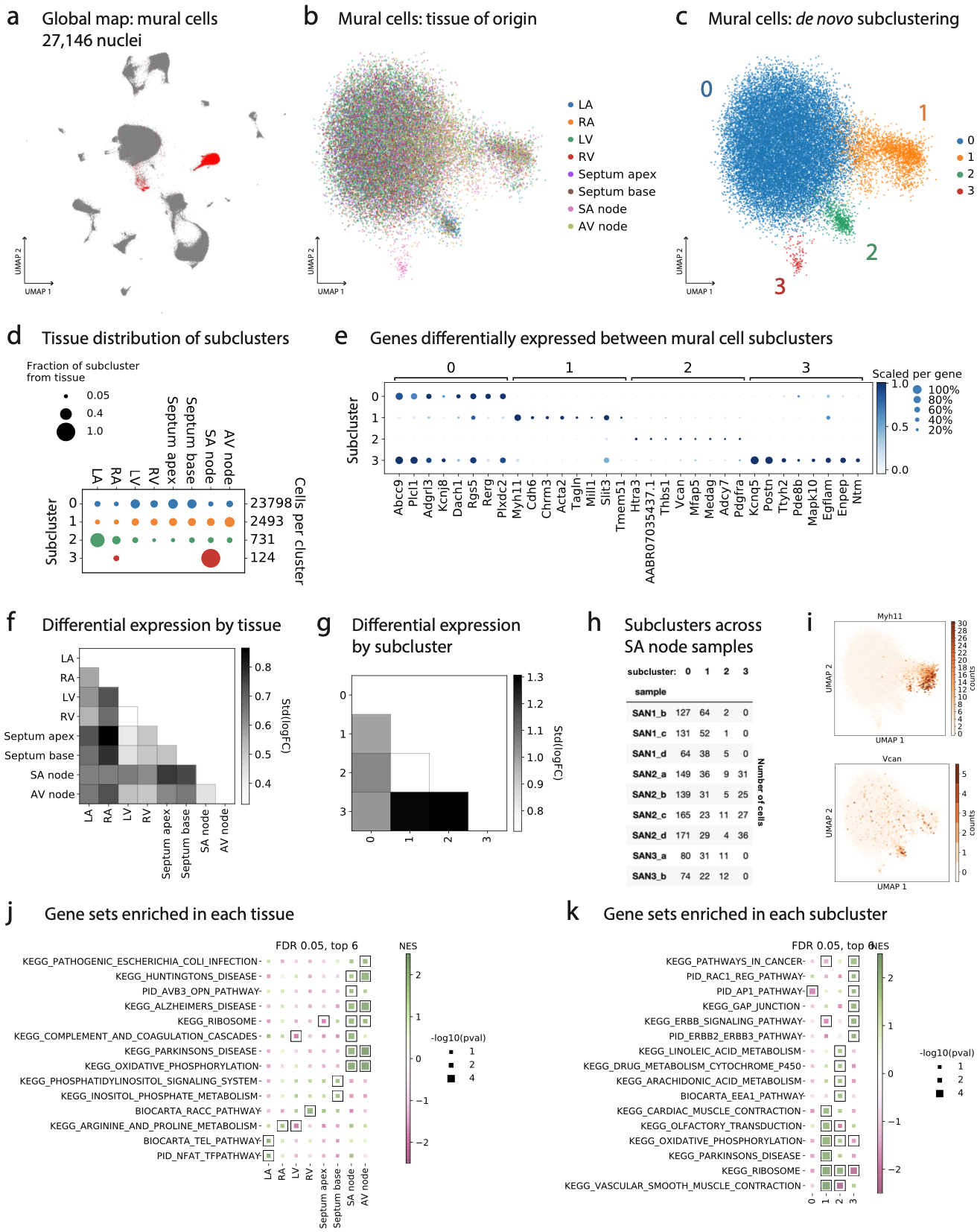

**Supplementary Figure 20. Subclustering of mural cells from the heart.** Includes pericytes and VSMCs. Excludes PV, PA, and Ao tissue samples. **(a)** UMAP showing 27146 relevant cells in red. **(b)** UMAP showing the tissue of origin of mural cells not from PV, PA, or Ao. **(c)** UMAP showing subclustering of the mural cells. **(d)** Tissue distribution of the subclusters. Dot sizes sum to one in each row. **(e)** Top marker genes of each subcluster. Subclusters 0 and 3 are pericytes, while subcluster 1 is VSMCs. **(f)** Summary of pairwise differential expression tests between tissues, for all mural cells. Lighter color means the tissues are more similar. **(g)** Same as panel **f**, showing pairwise differences between subclusters. **(h)** Due to the presence of subcluster 3 specifically in the SA node, we show the number of cells from each sample. All the subcluster 3 cells are coming from 4 replicates of one SA node tissue pool. **(i)** UMAP showing the expression of two marker genes, *Myh11* (in VSMCs) and *Vcan*. **(j)** KEGG, Biocarta, and PID gene sets enriched in each tissue (all mural cells included). **(k)** Same as **j** for each subcluster.

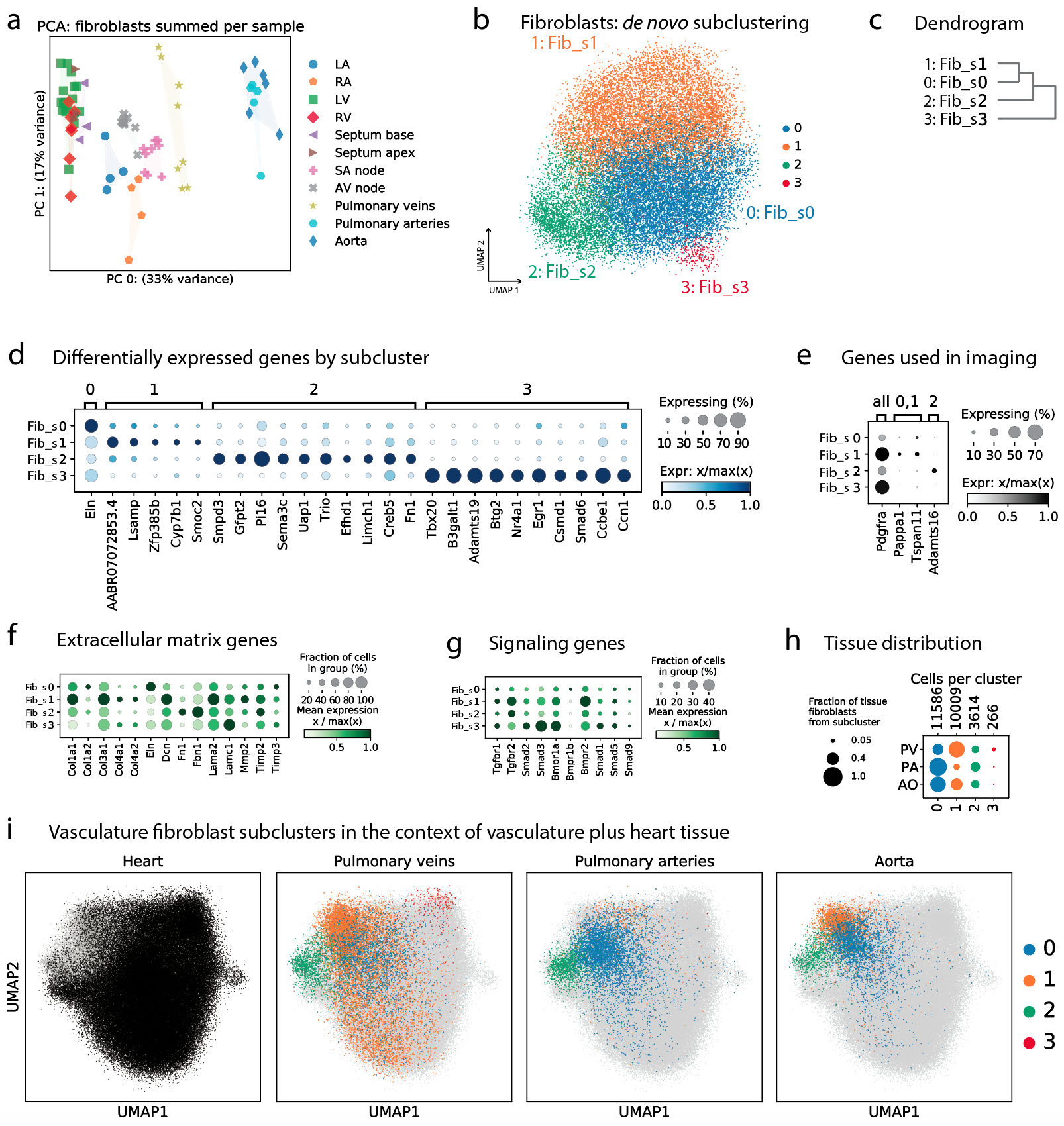

**Supplementary Figure 21. Subclustering of fibroblasts from PV, PA, and Ao. (a)** PCA plot showing the principal components of variation of pseudo-bulk FB expression in all samples. PV, PA, and Ao tissue samples clearly look different from other tissues. **(b)** Subclustering FBs from PV, PA, and Ao samples (25475 cells). FBs largely display a continuum of transcriptional profiles. **(c)** Dendrogram shows transcriptional similarity between subclusters. **(d)** Top differentially expressed genes in each subcluster. Even these top genes are not highly specific to one subcluster. **(e)** Genes used for imaging validation in **Supplementary Figure 22**. **(f)** Dotplot of a few extracellular matrix genes across subclusters. **(g)** Dotplot of a few signaling genes across subclusters. **(h)** Fraction of the total FBs in each tissue that come from each subcluster. Row dot sizes sum to 1. **(i)** UMAP panels showing all FBs from all tissues in light gray, overlaid by FBs from the heart (black) or the PV, PA, or Ao (color matches subclustering in panel **b**).

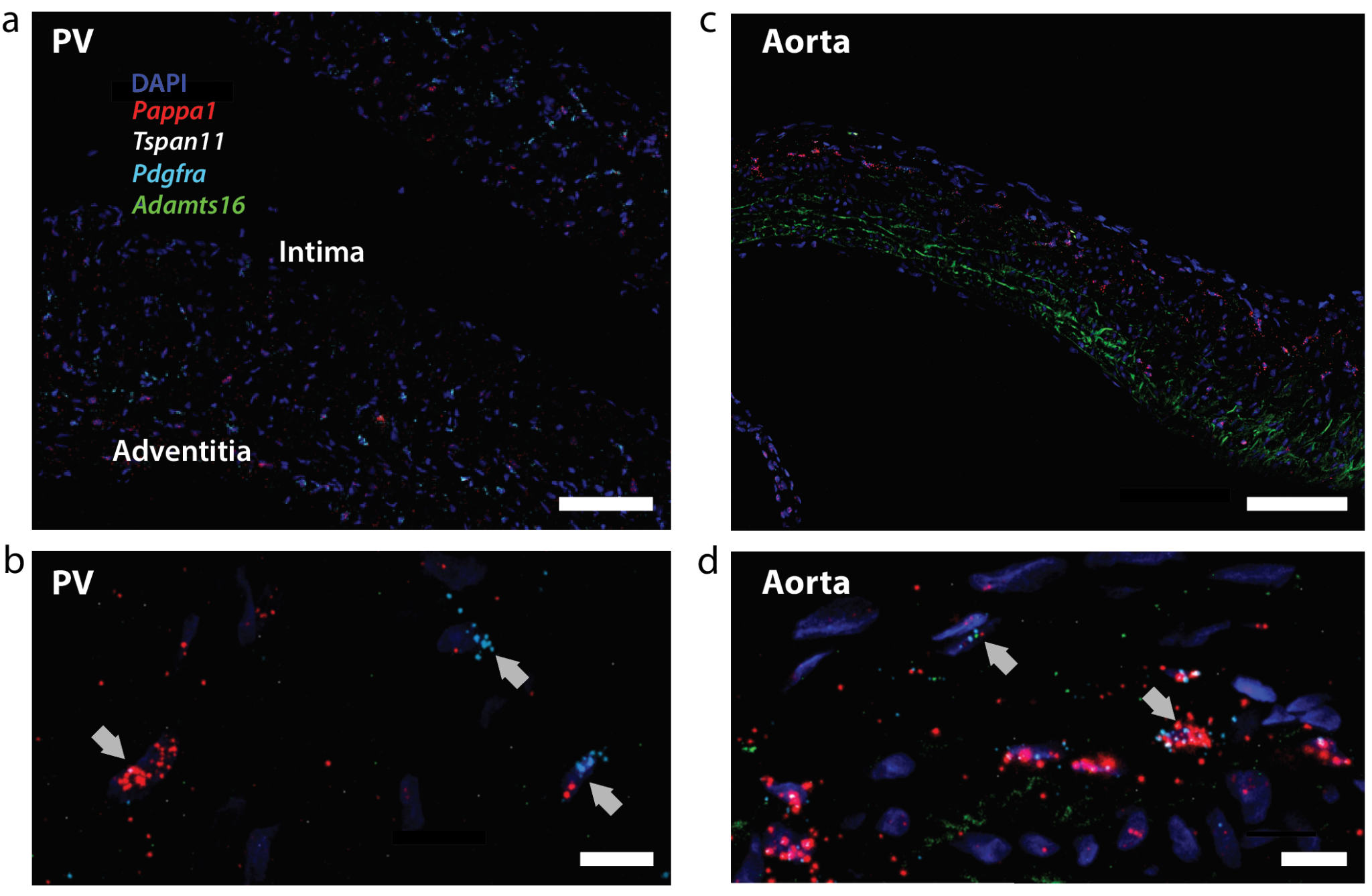

**Supplementary Figure 22. RNAscope imaging of vascular FB subtypes. (a)** Regional distribution of selected FB genes in PV tissue. Scale bar is 100 microns. **(b)** FBs in PV tissue are consistent with subclusters 0 and 1. Scale bar is 10 microns. Upward arrows show cells expressing *Pappa1* and *Pdgfra*. Downward arrow shows a cell with *Pappa1* and *Tspan11*. **(c)** Regional distribution of selected FB markers in Ao tissue. Scale bar is 100 microns. Green is *Adamts16*, which marks FB subcluster 2. **(d)** Downward arrow highlights a cell consistent with FB subclusters 0 and 1, expressing *Pappa1*, *Tspan11*, and *Pdgfra*. Upward arrow highlights a FB expressing *Adamts16*, consistent with subcluster 2. Scale bar is 10 microns.

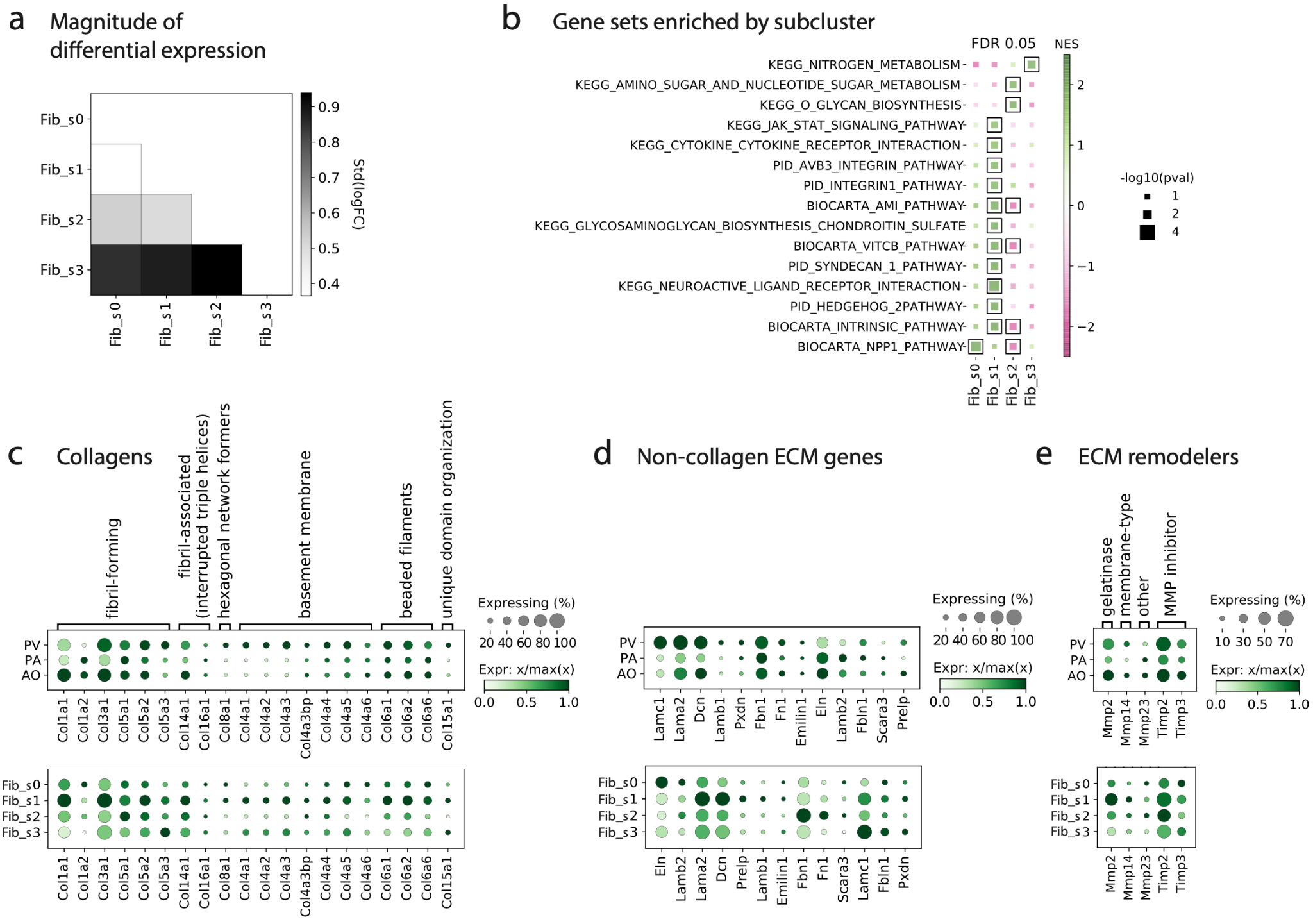

**Supplementary Figure 23. Characterization of vascular FB subtypes. (a)** Summary of magnitude of differential expression for all pairwise comparisons between FB subclusters from Supplementary Figure 21b. Lighter color indicates the two subclusters are more similar. **(b)** GSEA results for enriched KEGG, Biocarta, and PID pathways among the vascular FB subclusters. Boxed results are significant at FDR 0.05 after Benjamini-Hochberg multiple testing correction. **(c)** Expression of various collagens in FBs across vascular tissues and subclusters. **(d)** Other extracellular matrix (ECM) genes. **(e)** Genes involved in remodeling of the ECM.

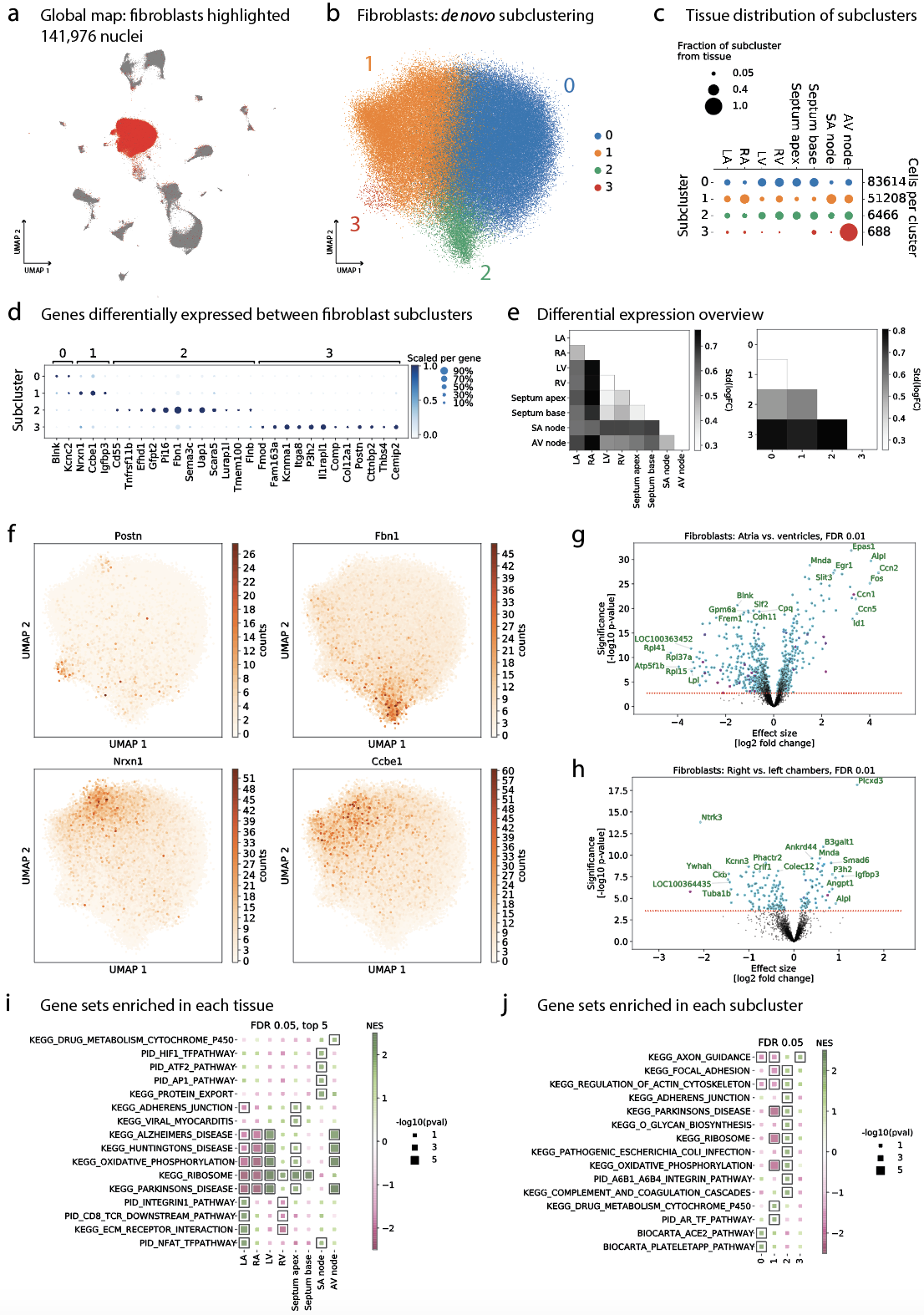

**Supplementary Figure 24. Subclustering FBs from the heart.** PV, PA, and Ao tissue samples are excluded. **(a)** Red color highlights 141,976 FBs on a UMAP of all cells from the heart. **(b)** Subclustering of the FBs (independent of the results in Supplementary Figure 21). **(c)** Tissue distribution of these FB subclusters. **(d)** Top differentially expressed genes between subclusters. **(e)** Overview of differential expression results for each one-versus-one comparison of all the FBs in each tissue (left panel) and subcluster (right panel). Lighter color denotes more similarity. **(f)** Four marker genes from panel **d**, plotted on the FB UMAP, show a considerable amount of variation. **(g)** Volcano plot shows the differential expression test results for all FBs from the atria versus all those from the ventricles. Magenta color denotes genes which were excluded based on being potential background noise. **(h)** Same as **g** for RA+RV versus LA+LV. **(i)** Top 5 (at most) significant GSEA results for KEGG, Biocarta, and PID pathways when FBs from each tissue were compared. **(j)** All significant GSEA results for KEGG, Biocarta, and PID pathways when heart FB subclusters were compared to each other.

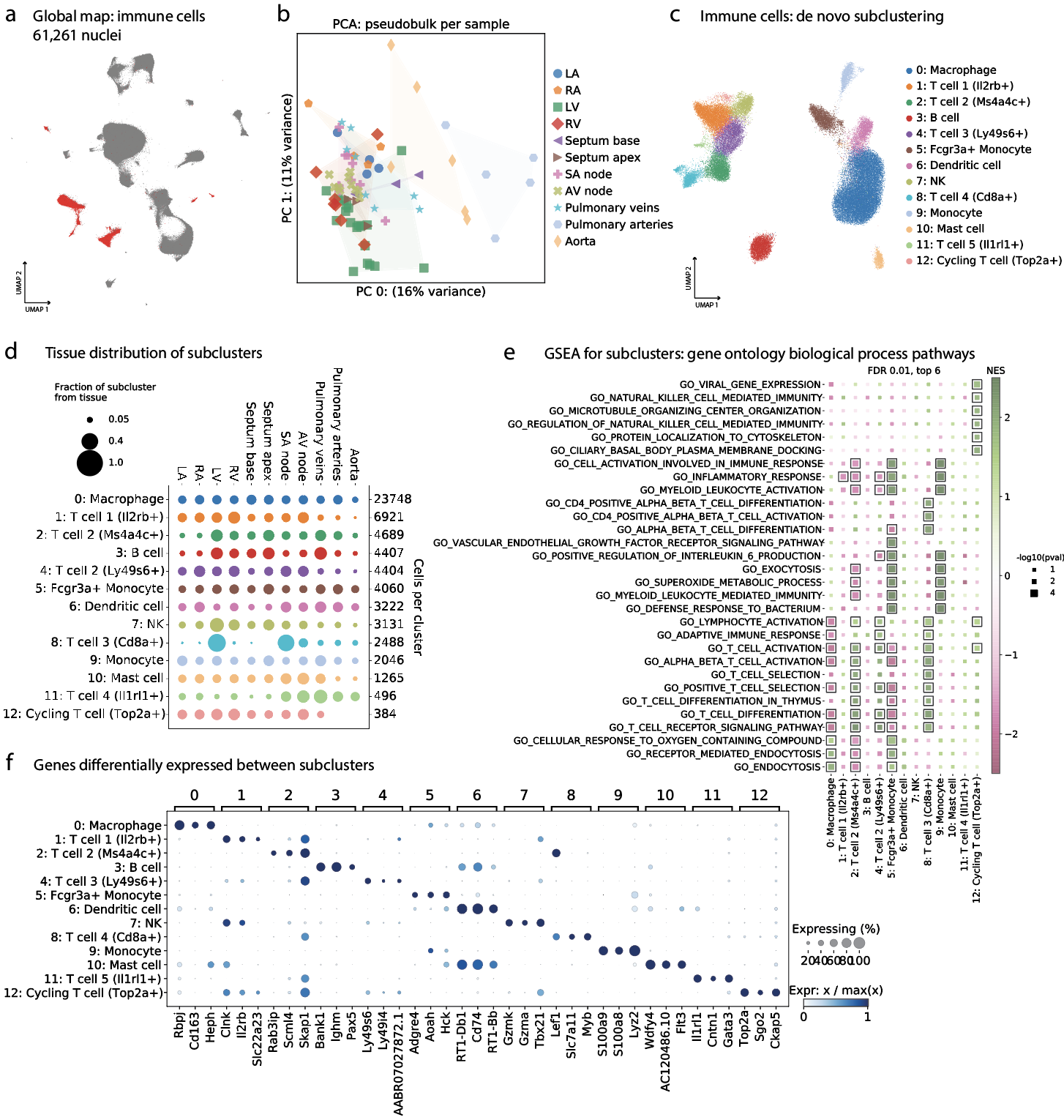

**Supplementary Figure 25. Subclustering of all immune cells.** **(a)** UMAP highlighting 61,261 immune cells in the overall UMAP in red. **(b)** Pseudo-bulk PCA plot obtained by summing expression of all immune cells in each sample. Each dot is the immune cells from a single sample, and dots are colored by tissue. **(c)** Subclustering of immune cells allows for more fine-grained resolution. **(d)** Distribution of immune cell subclusters across tissues. **(e)** Enrichment of GO biological process pathways identified by GSEA. **(f)** Top 3 differentially expressed genes in each immune cell subcluster.

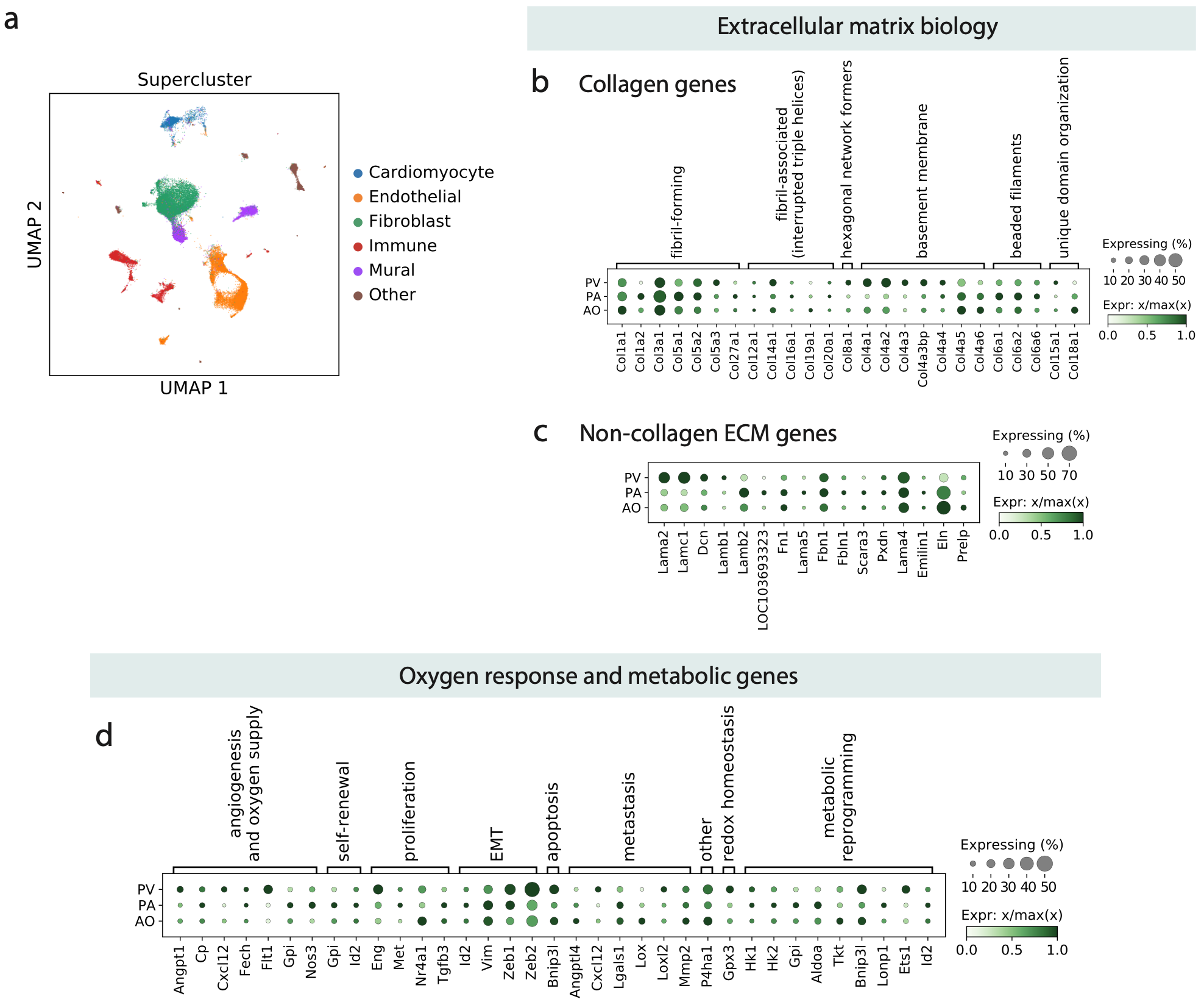

**Supplementary Figure 26. ECM and oxygen response genes in PV, PA, and Ao at a bulk level.** **(a)** Overview of UMAP of all cells from PV, PA, Ao where cells are labeled by major cell type group. **(b)** Dotplot of the expression of collagen genes by tissue. **(c)** Expression of non-collagen ECM genes by tissue. **(d)** Expression of Hif-regulated genes (oxygen response and metabolism) by tissue.

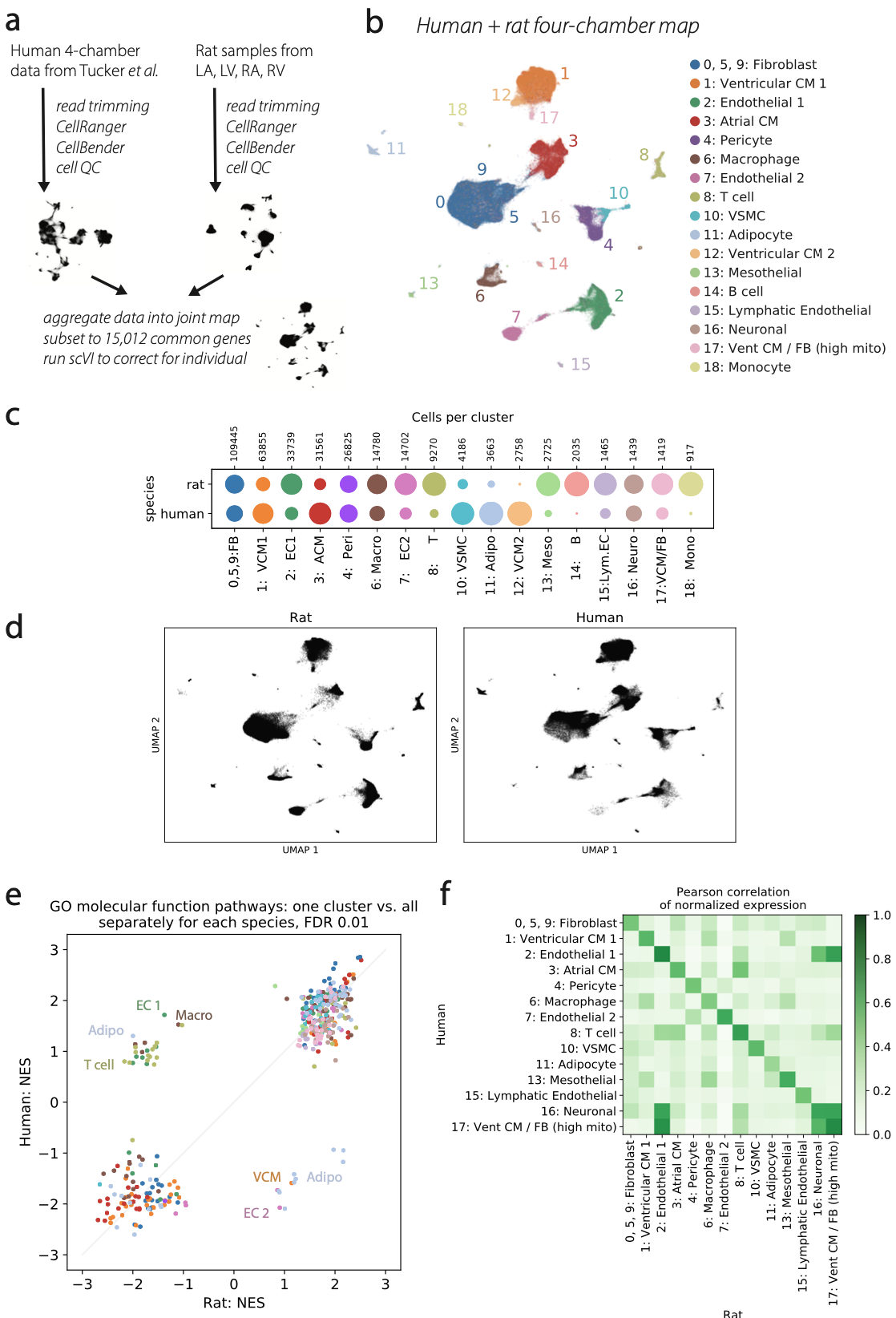

**Supplementary Figure 27. A combined rat and human healthy four-chamber cellular map. (a)** Data analysis pipeline includes rat data from LA, LV, RA, and RV, along with human data from Tucker *et al.* ^1^, reanalyzed here using an identical data processing pipeline. **(b)** UMAP of all human and rat cells that passed quality control. **(c)** Representation of each cluster in each species. Dot sizes represent the relative fraction of the cluster coming from each species. **(d)** UMAPs provide another way to visualize which cells come from which species. **(e)** Meta-analysis of GSEA results for GO molecular function pathways (c5.go.mf from MSigDB). GSEA was performed for each cluster on a species by species basis. Normalized enrichment scores (NES) from GSEA are then plotted, with one species on each axis, to try to understand concordance between the species. **(f)** Concordance of the cellular transcriptional profiles themselves, plotted as a heatmap of Pearson correlation of normalized expression. Clusters 12, 14, and 18 are excluded due to their near-absence in one species.

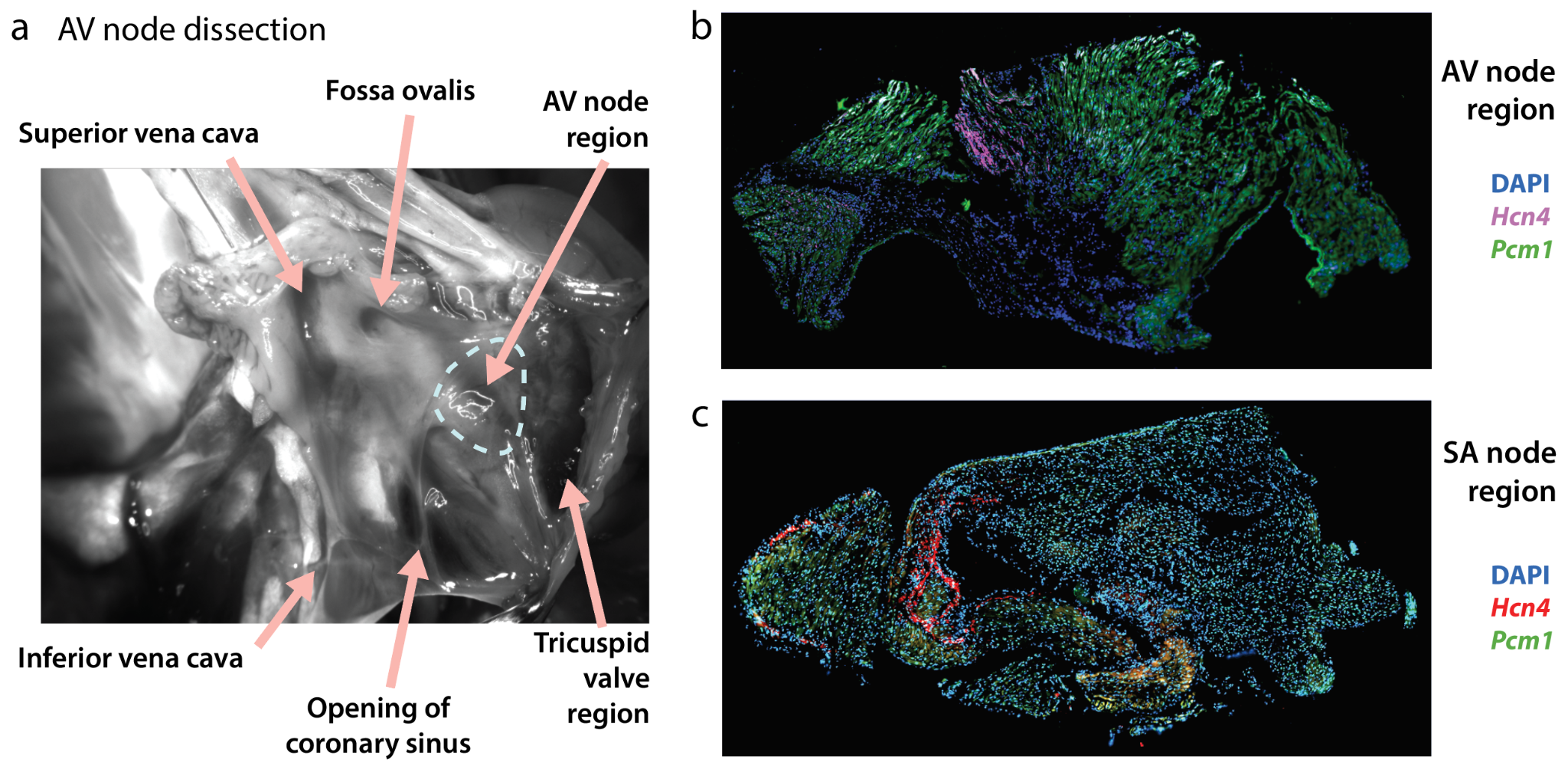

**Supplementary Figure 28. Obtaining nodal region samples containing pacemaker cells.** **(a)** Dissection of the AV node. Microscope image shows rat heart with opened right atrium, from a right lateral view. **(b)** Immunofluorescence image of the AV node region, staining *Hcn4* (purple; nodal pacemaker cardiomyocytes) and *Pcm1* (green), where nuclei are counterstained with DAPI (blue). **(c)** Same as **b** for the SA node region. *Hcn4* is in red. Both regions contain groups of *Hcn4*-positive cells, giving us confidence in our dissection of the AV and SA nodes. The dissected sample size is approximately 0.5mm^3^.

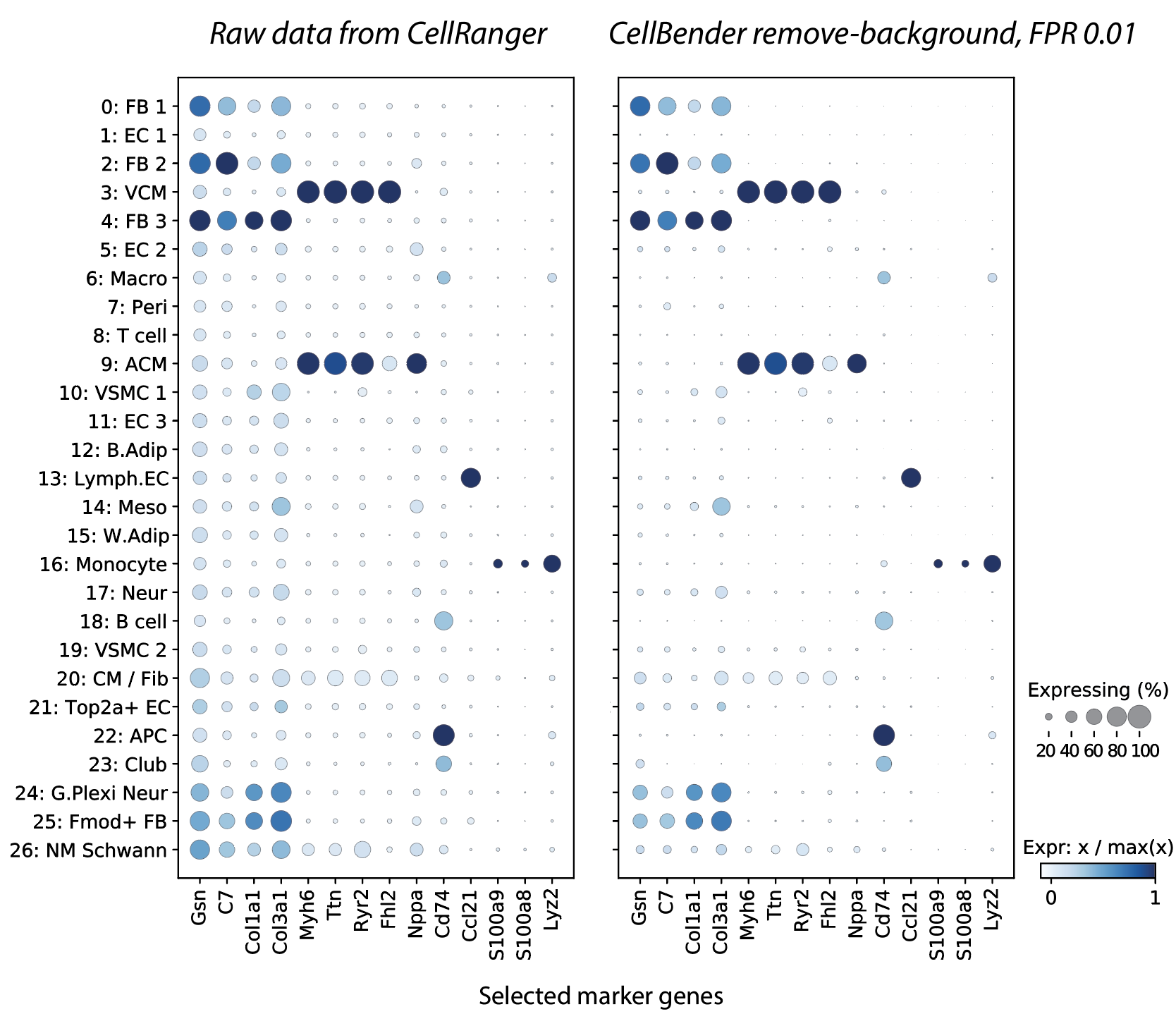

**Supplementary Figure 29. Removal of background noise by CellBender.** Effect of ambient RNA removal by cellbender remove-background v0.2.0. The side-by-side dotplots show the same selection of marker genes for (left) the raw data and (right) the data after CellBender preprocessing. We see improved specificity for these and other marker genes, and a diminution of the background noise. Each gene’s expression (dot color) is scaled by dividing a cluster’s mean expression by the minimum value of the mean expression of that gene in any cluster. Note that we are not scaling so that the minimum cluster expression is assigned the value zero on the color axis, as is common practice.

### Supplementary Tables

#### T1: CellRanger QC metrics and experimental design

#### T2: Cluster DE

#### T3: Tissue DE per cluster

#### T4: Cluster GSEA

#### T5: Intra-supercluster GSEA

#### T6: Cell-cell communication

#### T7: Tissue enrichment of cell-cell communication

#### T8: Rat and human gene linker
